# *Helicobacter pylori* attachment-blocking antibodies protect against duodenal ulcer disease

**DOI:** 10.1101/2023.05.24.542096

**Authors:** Jeanna A. Bugaytsova, Kristof Moonens, Artem Piddubnyi, Alexej Schmidt, Johan Olofsson Edlund, Gennadii Lisiutin, Kristoffer Brännström, Yevgen A. Chernov, Kaisa Thorel, Iryna Tkachenko, Oleksandra Sharova, Iryna Vikhrova, Anna Butsyk, Pavlo Shubin, Ruslana Chyzhma, Daniel X. Johansson, Harold Marcotte, Rolf Sjöström, Anna Shevtsova, Göran Bylund, Lena Rakhimova, Anders Lundquist, Oleksandra Berhilevych, Victoria Kasianchuk, Andrii Loboda, Volodymyr Ivanytsia, Kjell Hultenby, Mats A. A. Persson, Joana Gomes, Rita Matos, Fátima Gartner, Celso A. Reis, Jeannette M. Whitmire, D. Scott Merrell, Qiang Pan-Hammarström, Maréne Landström, Stefan Oscarson, Mario M. D’Elios, Lars Agreus, Jukka Ronkainen, Pertti Aro, Lars Engstrand, David Y. Graham, Vladyslava Kachkovska, Asish Mukhopadhyay, Sujit Chaudhuri, Bipul Chandra Karmakar, Sangita Paul, Oleksandr Kravets, Margarita Camorlinga, Javier Torres, Douglas E. Berg, Roman Moskalenko, Rainer Haas, Han Remaut, Lennart Hammarström, Thomas Borén

## Abstract

The majority of the world population carry the gastric pathogen *Helicobacter pylori*. Fortunately, most individuals experience only low-grade or no symptoms, but in many cases the chronic inflammatory infection develops into severe gastric disease, including duodenal ulcer disease and gastric cancer. Here we report on a protective mechanism where *H. pylori* attachment and accompanying chronic mucosal inflammation can be reduced by antibodies that are present in a vast majority of *H. pylori* carriers. These antibodies block binding of the *H. pylori* attachment protein BabA by mimicking BabA’s binding to the ABO blood group glycans in the gastric mucosa. However, many individuals demonstrate low titers of BabA blocking antibodies, which is associated with an increased risk for duodenal ulceration, suggesting a role for these antibodies in preventing gastric disease.

## INTRODUCTION

Chronic life-long infection by *Helicobacter pylori* is the main cause of chronic active gastritis, which can lead to severe clinical conditions including gastric and duodenal ulcer disease and gastric cancer. All these diseases except gastric cancer have become curable by *H. pylori* eradication ^1, 2^; however, gastric cancer remains a major threat to human life with about 1.2 million new cases every year with a poor survival rate, with ∼25 million deaths since the year 2000 ^3–5^. About 5–10% of *H. pylori*-infected individuals experience duodenal ulcer disease, which, although curable, can perforate causing bleeding with a high mortality ^6^. Patients with ulcers or gastric cancer typically carry the more virulent “triple-positive infections” characterized by *H. pylori* bearing the CagA onco-effector protein, the VacA toxin, and the Blood group Antigen Binding (BabA) attachment protein (adhesin) ^7, 8^. BabA is a member of the *H. pylori* outer membrane protein family and binds to the ABO/Lewis b (Leb) blood group antigens expressed in the gastro-intestinal lining, including the gastric epithelium ^9–12^. BabA exhibits rapid and diversifying selection through amino acid substitutions in its Leb-<u>c</u>arbohydrate <u>b</u>inding <u>d</u>omain (CBD) that adapts its binding preferences for different human populations and ABO blood group phenotypes ^11, 13, 14^. The rapid mutation and selection in BabA also allow for adaptation to the individual and optimal binding to the different locations and microenvironments in the stomach, including adaptation to shifts in acidity caused by diseases such as peptic ulcer disease and gastric cancer and adaptation by shifts in its binding affinity ^14^. As a consequence, BabA exhibits extensive global diversity where nearly every infected individual will harbor a strain of *H. pylori* possessing a unique BabA sequence. *H. pylori* infections induce high antibody titers against several dominant antigens, such as its urease enzyme, but these antibodies fail to clear the infection. Hence, *H. pylori* infections instead become persistent and life-long. By contrast, most *H. pylori* outer membrane proteins, including BabA, are reported to be weakly antigenic ^15–17^. Immunoglobulin G is induced by gastro-intestinal (GI) pathogens also in the gut and by *H. pylori* virulence factors in both the gastric antrum and the corpus mucosa ^18, 19^. Therefore, we tested ∼1000 patient sera and found that the vast majority exhibited blocking IgG antibodies (Abs) that inhibited BabA-mediated Leb binding. Surprisingly, the patient sera turned out to efficiently block Leb-binding of world-wide *H. pylori* strains, suggesting the presence of <u>b</u>roadly <u>b</u>locking IgG <u>Abs</u> (bbAbs). We also found that patients with duodenal ulcer disease (DU) exhibited low titers of bbAbs. This result suggests that the IgG-mediated blocking activity is protective throughout the GI tract. To identify the bbAb binding epitopes in BabA, we cloned a human antibody, <u>A</u>nti-<u>B</u>a<u>bA</u>, denoted as ABbA, from healthy *H. pylori* carriers and co-crystalized it with BabA. The co-crystal structure showed that ABbA and Leb bind to the same residues in the BabA CBD. Thus, ABbA is resistant to microbial antigen variation and instead performs fulminant glycan mimicry to block and compete with BabA-mediated Leb binding to reduce mucosal attachment of *H. pylori*. Our results indicate that the host’s blocking antibody response has co-evolved with *H. pylori* as a way of tolerating chronic infection and reducing the risk for overt gastric disease by mitigating mucosal inflammation. We believe that the bbAbs reported here offer translational applications in treatment and predictive diagnostics for individuals at risk for severe gastric disease.

## RESULTS

### Human serum antibodies block ABO/Leb binding and attachment of *H. pylori* bacteria to the gastric mucosa

To test if human immune responses during chronic *H. pylori* infection raise factors that reduce bacterial adherence, we tested sera from six individuals, four of whom were ELISA positive for *H. pylori* and two of whom were *H. pylori* negative. We found that three of the four positive sera samples fully inhibited or reduced epithelial attachment of the clinical isolate *H. pylori* J166 to the human gastric mucosa (**Figure 1A**). Two tests were used to identify the nature of the serum samples’ inhibitory activity. First, the inhibitory activity was scored as the serum dilution at which half maximal *H. pylori* bacterial binding to Leb was lost, i.e., the 50% Inhibition Titer (IT50). Hence, the IT50s tightly reflected the strength of the serum samples in inhibition of *H. pylori* attachment to the gastric mucosa (**Figure 1B**). Second, the IgG antibodies were specifically desorbed from the serum sample, which resulted in >95% loss of the inhibitory activity (**Figure 1C**), suggesting that the broadly blocking activity resides in the IgG antibody fraction. These results show that individuals with *H. pylori* infection can raise attachment-blocking IgG antibodies that inhibit Leb binding and thus reduce *H. pylori* adherence to the gastric mucosa.

**Figure 1.**
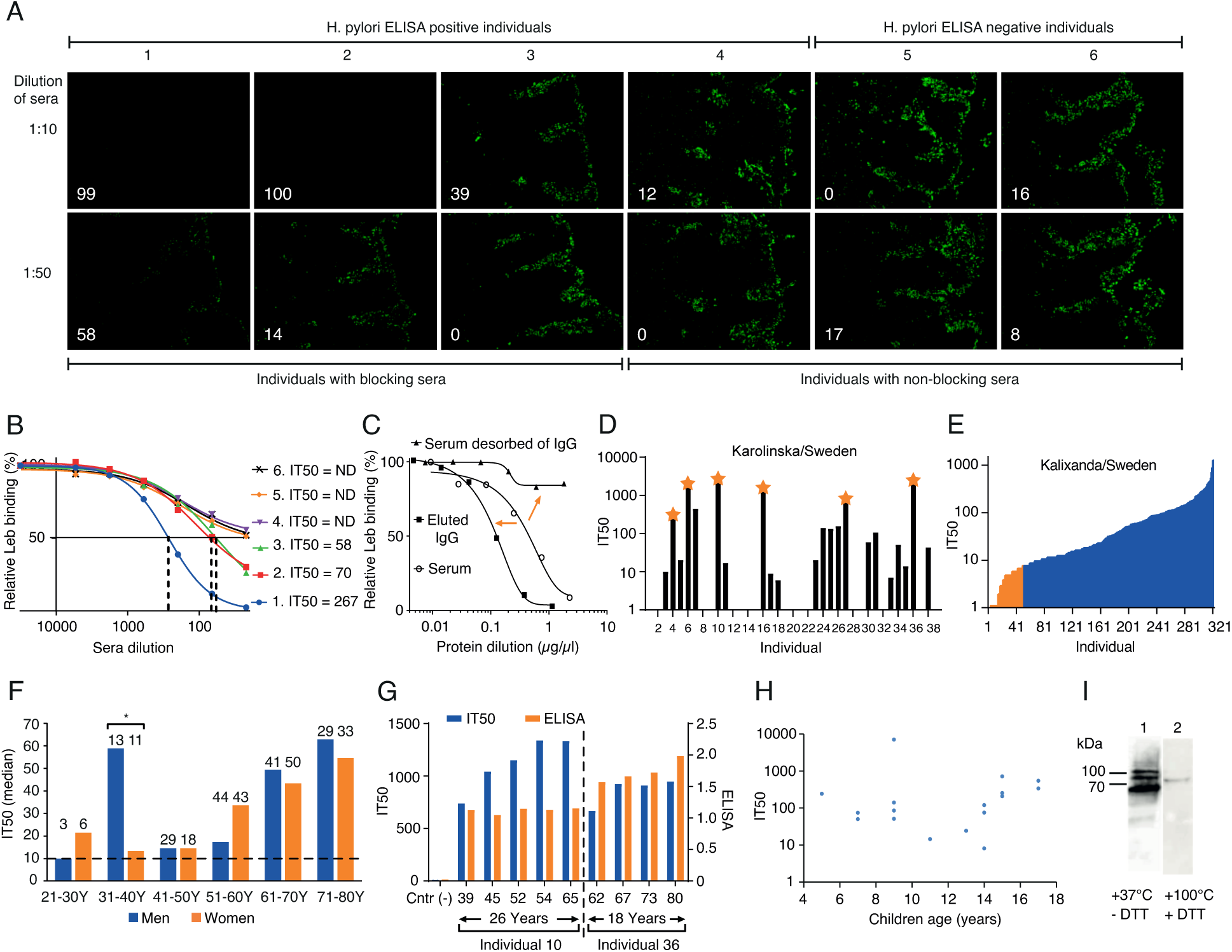
High global prevalence of serum inhibition of Leb binding. (A) Inhibition (% in white digits) of *in vitro* binding to human gastric mucosa by *H. pylori* (**Figure S1A**) from six individual serum samples (quantified in **Figure S1B**). (B) The individual sera IT50 was determined by incubation of *H. pylori* J166 with ^125^I-Leb-conjugate in a dilution series of serum sample. The dashed vertical lines show the serum dilution when Leb binding was reduced by 50%, i.e., the serum dilution that equals the IT50. Serum from individuals 4–6 did not produce a 50% reduction (Non-Detected (ND) IT50)). (C) The serum sample from individual 1 lost its blocking activity after IgG affinity desorption (right arrow) (**Figure S1C**). The IT50/blocking activity was recovered in the IgG fraction from the Protein G affinity column (left arrow). (D) The sera IT50s of 38 individuals from KI, Sweden (**Table S1A**). The stars indicate serum samples analyzed in **Figure 2A**. (E) The IT50s of 322 individuals from the Swedish Kalixanda study tested by strain 17875/Leb with a median IT50 = 41 and a mean IT50 = 89. The J166 strain was more sensitive because of its lower Leb binding affinity and identified 84% of the sera as IT50 positive (**Table S1B** and **Figure S2B**). (F) The median IT50s of the 322 Kalixanda individuals according to gender and age. The median IT50 was identical between all the men and the women but increased with age (**Figures S1E*i*** and **S1E*iv***). (G) The IT50s and *H. pylori* ELISAs of individuals 10 and 36 from **Figure 1D** were followed over 26 and 18 years, respectively. The IT50s increased with age during the ∼50-year observation period in contrast to the almost constant ELISA titers. (H) The children’s IT50s tested by strain 17875/Leb varied similarly to adults by 100– 1000-fold (**Table S1F**). (I) Immunoblot detection of purified native BabA protein and its high-molecular-weight multimers ^14^ in the serum sample of individual 10 from KI, Sweden (**Figures 1D** and **1G**) and in the serum samples from world-wide populations (**Figure S1H**) separated under semi-native conditions (1) or denaturing and reducing conditions (2).

### Worldwide high prevalence of serum IT50s

The results from **Figure 1A** suggest that many individuals infected by *H. pylori* might be positive for Leb IT50s. Thus, in a pilot study we tested a group of 38 asymptomatic carriers of *H. pylori* infection from the Karolinska University Hospital (KI), Sweden, and found that 22 (58%) of the sera reduced Leb binding with IT50s that ranged 100-fold, from ∼10 to >2000 (**Figure 1D**). To determine if most individuals who are ELISA-positive for *H. pylori* raise antibodies that inhibit Leb binding, we tested 742 individual serum samples from both healthy individuals and from patients with gastric disease. For the IT50 tests, we used two *H. pylori* strains, 17875/Leb and J166, described in **Tables S1B-E**), with high ^11^ *vs*. low (**Figure S1D** and **Figure S2B**) Leb-binding affinity, respectively. First, we tested a group of 322 volunteers, all ELISA positive, i.e., *H. pylori* carriers, from the Kalixanda Study of the Swedish general adult population, an essentially “gastric healthy” cohort ^20^. We found that the Kalixanda serum samples similarly ranged 100-fold in IT50s and identified 270 (84%) IT50-positive sera whereas the remaining 16% did not demonstrate IT50s above background and were defined as negative (**Figure 1E**). Notably, the IT50s demonstrated a strong correlation with the presence of antibodies against the onco-marker CagA (**Figure S1E*ii***), whereas the IT50-positive sera did not correlate with the *H. pylori* ELISA titers (**Figure S1E*iii***). Second, for comparison, 420 patients with *H. pylori* infection and gastric disease from Ukraine, the US, and Mexico were tested, and 56%, 59%, and 72% were IT50-positive, respectively (**Figure S1F**). Leb binding tests showed that 81% (26/32) of the Mexican *H. pylori* clinical isolates bound to Leb (**Table S2B**), and the corresponding sera IT50s correlated with Leb binding (Wilcoxon p = 0.036) (**Figure S1G**). Thus, the worldwide high prevalence of IT50s against Leb binding closely mirrors the 60%–80% world-wide prevalence of Leb binding among *H. pylori* isolates ^11^.

### Serum IT50s increase with age

The IT50s in the Kalixanda population increased from mid-age to high-age in both women and men, with two exceptions – in the 31–40-year age group men had 4-fold higher IT50s (Wilcoxon rank, p = 0.028) (**Figure 1F**). It is noteworthy that the IT50 augmentations coincided with men’s age of onset of DU ^20, 21^, where the age-matched increased IT50s might constitute protective antibody responses, as only 5% of the Kalixanda individuals had macroscopic gastric lesions ^20^. To test if the IT50 increases with age also at the individual level, we tested serum samples collected from two individuals in the timespans from 39 to 65 years of age and from 62 to 80 years of age, respectively. Similar to the Kalixanda cohort, both individuals’ IT50s increased by age and reached a high plateau at ∼55–65 years of age. In contrast, the ELISA levels did not change, which agreed with the lack of correlation between ELISA and IT50 **(Figure 1G** and **Figure S1E*iii*)**. This suggests that the IT50s do not level out or decrease at higher age either on the population level or on the individual level. To understand how early IT50s develop, we tested children from 3 to 17 years of age (all ELISA-positive *H. pylori* carriers). We found that 47% (17/36) of the sera demonstrated IT50s from 5 years of age (**Figure 1H**). The combined results showed that antibodies that inhibit Leb binding are formed already at an early age in response to *H. pylori* infections and are present and active throughout the lifespan.

### The serum IT50 antibodies bind to structural BabA epitopes

To characterize the epitope preferences for the antibodies that block Leb binding, we tested serum samples from each of the KI/Sweden, Ukraine, US, Mexico, and Kalixanda populations. The sera samples from all six global populations detected the Blood group Antigen Binding (BabA) protein, but only under semi-native conditions, suggesting that the antibodies do not recognize the denatured, linear epitopes but preferentially bind to structural epitopes in BabA (**Figure 1I** and **Figure S1H)**.

Thus, testing of the 780 global individual *H. pylori* carriers suggested that (1) blocking antibodies (IT50s) are present in a vast majority of individuals, (2) IT50 levels are highly individual and do not correlate with the ELISA level, (3) the IT50 responses are more prominent against the oncogenic CagA-positive *H. pylori* infections, (4) the IT50s are active over the lifetime of the individual, and (5) the IT50s are raised against world-wide conserved structural BabA epitopes instead of antigenic variable linear epitopes.

### The serum IT50 is derived from the activity of broadly blocking antibodies

To determine if the individual IT50 can also block Leb-binding of global, phylo-distant *H. pylori* strains, six Swedish serum samples (stars in **Figure 1D**) were tested against 12 *H. pylori* isolates from Europe, Asia, and the Americas, including the IT50 reference strains 17875/Leb and J166 (**Figures 2A** and **2B**). All six serum samples inhibited Leb binding of all 12 strains, i.e., all individual sera exhibited broadly blocking antibody activity (**Figure 2A**). The IT50s against strain 17875/Leb reflected the general inhibition strength of the six serum samples. Notably, the two LOW inhibitory sera preferentially inhibited the *H. pylori* strains J166 and I9, which were lower in Leb binding affinity (**Figure S1D** and ^14^). Reciprocally, the four HIGH inhibitory sera were less discriminating in inhibiting the 12 strains (**Figure S2A**). The results on the sera broadly blocking antibody activities help explain why the series of 780 global serum samples were so efficient in inhibiting strains 17875/Leb and J166 (**Figures 1** and **S1**) that both originate from Europe (**Figure 2B** and^22^.

**Figure 2.**
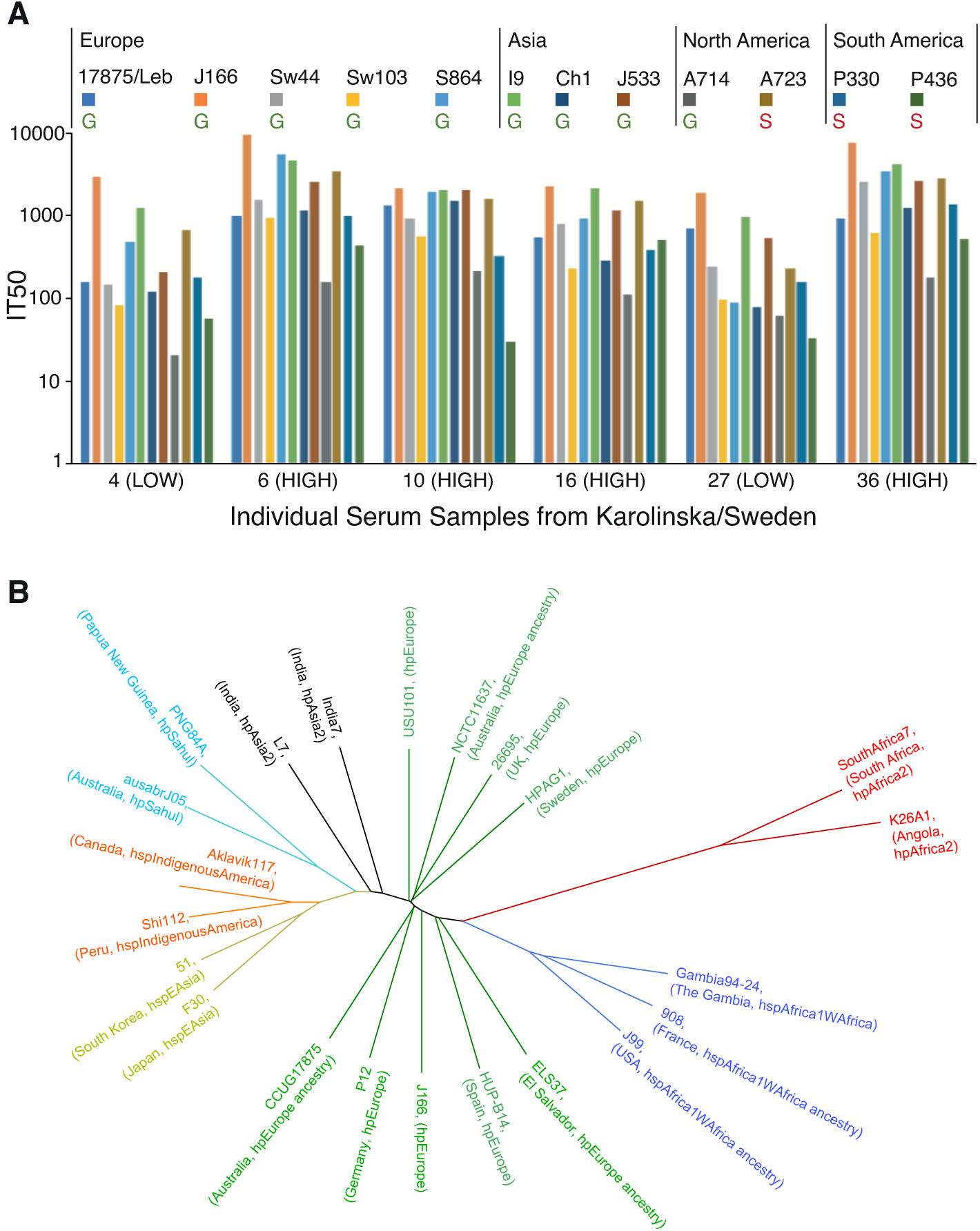
Broadly blocking serum Abs. (A) Serum samples from 6 Swedish individuals were tested against 12 world-wide strains, including 9 ABO antigen-binding Generalist (G) and 3 Indigenous American Specialists (S) ^11^. The BabA alignments and heterogenicities are shown in **Figure S2C**, and the strain binding characteristics are shown in **Figure S2B**. (B) Phylogenetic tree with the *H. pylori* strains CCUG 17875 (17875/Leb, GenBank CP090367) and J166 ^22^ both from central/southern Europe, TIGR26695, NCTC11637, P12, HPAGA1, HUP-B14, USU101 https://www.ncbi.nlm.nih.gov/nuccore/NZ_CP032818.1, and ELS37 in the *hp*Europe domain (green) and their distance from the African (red and blue, including J99 of African phylogeny ^40^), South Asian (black), Oceanian (turquoise), East Asian (yellow), and Indigenous American (orange) *H. pylori* strains.

### *H. pylori* with higher-affinity Leb-binding is a risk factor for overt gastric disease

To understand how Leb binding affinity affects the serum inhibition of *H. pylori* attachment to the gastric mucosa, we exposed strains 17875/Leb and J166, with high *vs.* low Leb binding affinity, respectively, to serum from individual 1 from **Figures 1A and 1B**. A 25-fold dilution of the serum reduced the attachment of strain J166 10-fold compared to strain 17875/Leb (p < 0.0001) (**Figure 3A**). Thus, the low-affinity Leb-binding strain J166 was more efficiently inhibited by serum antibodies in terms of both Leb binding (**Figure 2A**) and attachment to the gastric mucosa (**Figure 3A**). Thus, we re-analyzed the series of Swedish *H. pylori* isolates from patients with GA or DU for their Leb affinity ^11^ because the CagA, VacA, BabA “triple positive” virulence markers are present in almost all DU and gastric cancer strains ^7, 8^. Indeed, the DU strains demonstrated higher binding affinity (Wilcoxon p ≤ 0.05) (**Figure 3B** and **Table S2A**). Similarly, the Mexican Leb-binding DU isolates demonstrated higher binding strength compared to the GA isolates (Wilcoxon signed-rank test, p = 0.0015) (**Figure S3A**).

**Figure 3.**
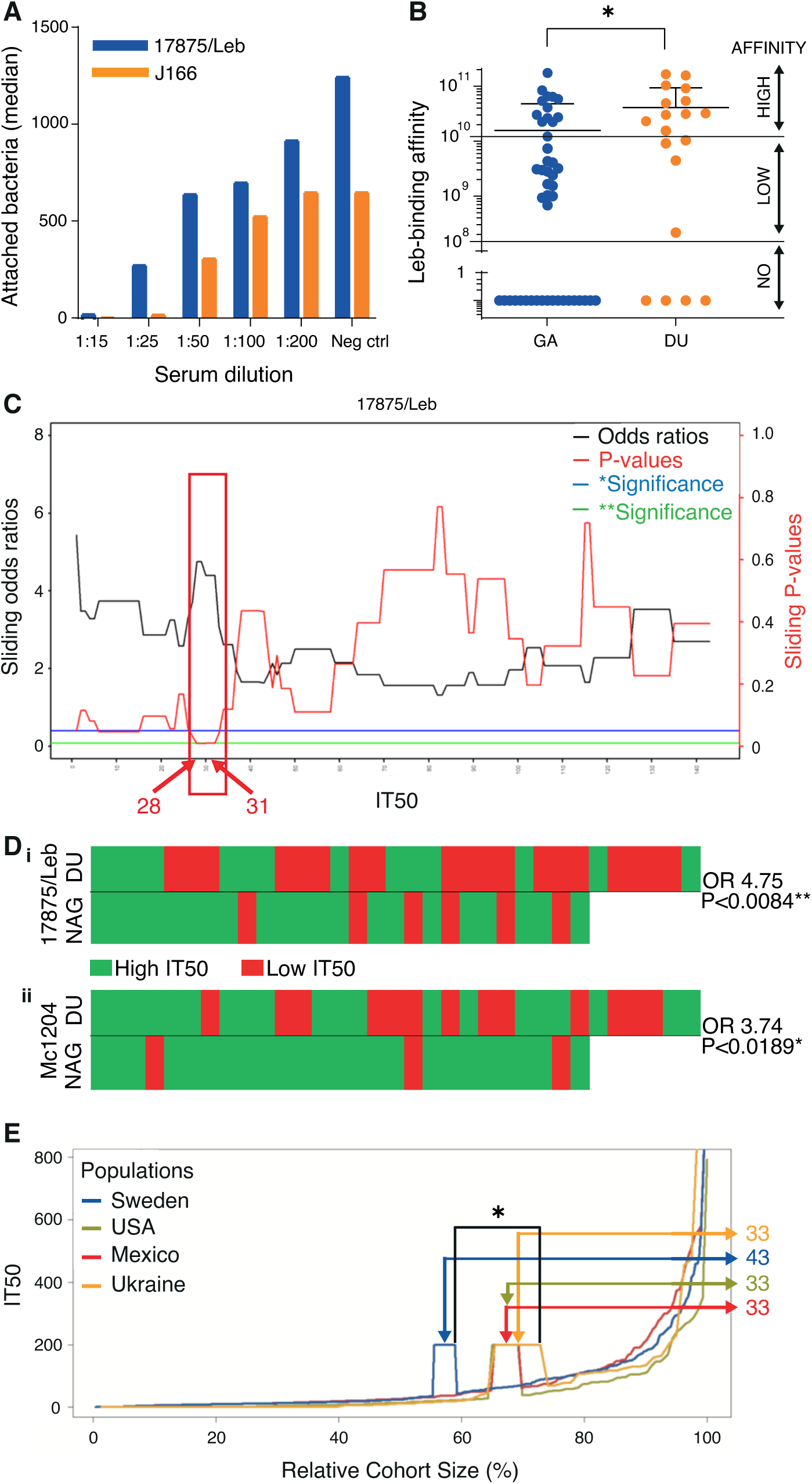
Broadly blocking serum Abs protect against overt gastric disease. (A) The serum sample from individual 1 in **Figure 1A** blocked the in vitro binding, i.e., attachment, of *H. pylori* to human gastric mucosa. In the 1:25 dilution, the 17875/Leb and J166 strains demonstrated 28% and 3% residual attachment, respectively (**Figure S3A**). (B) Among Leb-binding Swedish DU and gastritis (GA) isolates ^11^, ∼80% (12/15) of DU isolates were found to exhibit higher Leb-binding affinity compared to ∼50% (13/27) of the GA isolates (**Table S2A**). (C) The sliding window X-axis and ROC test (**Figure S3E*i***) show the IT50s for strain 17875/Leb that were used to calculate the series of ORs (black line) and their corresponding p-values (red line) (IT50s from **Table S3**). The IT50 = 28–31 interval with a significant OR is indicated by the red box. (**D***i*) Individual sera from the NAG *vs.* DU group, where the critical IT50 ∼30 for 17875/Leb was identified as an Rf for DU (IT50 > 30 in green, IT50 < 30 in red) with OR 4.75; 95%CI: 1.58–15.86, and p < 0.0084**. (**D***ii*) The *H. pylori* Mc1204 isolate (**Table S2B**) identified the Rf IT50 ∼15 with OR 3.74; 95%Cl: 1.12–14.98, and p < 0.0189* (**Figures S3D*i*** and **S3E*ii***) **(IT50s** from **Table S3)**, whereas the low affinity Leb-binding strains J166 and Sw44 (**Figure S2B**) did not produce significant ORs (**Figures S3D*ii*** and **D*iii*)**. (**E**) The serum samples from Kalixanda (Sweden), the US, Mexico, and Ukraine were ordered according to their IT50 and relative sample size. The three cohorts with gastric disease (the US, Mexico, and Ukraine) exhibited only 33% of individuals above the Rf IT50 = 30, compared to 43% in the general adult “gastric healthy” Kalixanda (Swedish) cohort (**Table S4**); p = 0.014 and p = 0.056 for the higher *vs.* the lower samples of the intervals.

### Low IT50 is a risk factor for overt gastric disease

Because more blocking antibodies, i.e., higher IT50s, are required to prevent the HIGH-affinity Leb-binding *H. pylori* strains from adhering to the gastric mucosa, and because the “triple-positive strains” with BabA of HIGH Leb-binding strength constitute a risk factor for overt gastric disease, we next tested if a low individual IT50 can be regarded a risk factor for overt gastric disease. Importantly, during gastric cancer (GC) progression and loss of gastric acidity, the *H. pylori* infections often fade and disappear, with a parallel decrease or loss of antibodies against *H. pylori* ^23^. Thus, the IT50s from patients with GC could be erroneously low. Instead, DU patients were chosen for this investigation because the chronic *H. pylori* infection can readily adapt to the acidified environment. Hence, we selected a series of 79 serum samples from age-matched patients with DU or Non-Atrophic “healthier” Gastritis (NAG) from Mexico. First, the IT50s ranged several 100-fold, but the NAG sera displayed generally higher IT50s (**Figure S3B**), whereas the general ELISA titers were similar (**Figure S3C**). Second, we calculated the full range of odds ratios (ORs) and their p-values for sera samples with high *vs.* low IT50 for the NAG *vs.* DU group. By using sliding windows and ROC diagrams, we identified IT50 ∼30 as a risk factor (Rf) for DU, by strain 17875/Leb, where the highest OR of 4.75 coincided with the highest significance (**Figures 3C** and **3D*i***). We also tested the high-affinity Leb-binding Mc1204 isolate from the cognate Mexican group and identified IT50 ∼15 as an Rf where the highest OR of 3.74 coincided with the highest significance (**Figures 3D*ii*** and **S3D*i***), i.e., the two strains produced similar low IT50s as Rfs. The critical ORs were confirmed by ROC diagrams (**Figure S3E**) and by permutation tests (**Figure S3F**). This series of results suggest that individuals with low IT50 serum levels are at a higher risk for DU. Notably, the 1:15 serum dilution required to completely block attachment to the gastric mucosa of the high affinity binding strain 17875 (**Figure 3A**) corresponded to IT50∼10 when tested using 17875. Thus, the IT50 serum titers must exceed the critical Rf IT50 ∼30 in order to provide for the IT50∼10 level of adherence blocking efficacy in the gastric mucosal lining. To understand how the Rf IT50 ∼30 relates to gastric disease in populations worldwide, we next aligned the IT50s of all individuals from Kalixanda, Ukraine, USA, and Mexico. This identified a ∼30% higher prevalence of individuals with protective IT50s >30 in the healthier Kalixanda general adult population compared to the global gastric disease patients and hence a predisposition for lower IT50 among individuals with gastric disease (**Figure 3E** and **Table S4**). Our results suggest that DU patients often carry the high-affinity Leb-binding *H. pylori*, i.e., the strains that require higher IT50 sera for blocking of Leb binding and hence reduction of *H. pylori* attachment; however, the DU group of patients often presented with lower and thus insufficient IT50 titers of protective bbAbs.

### Cloning and identification of the bbAbs

For a better understanding of the mAbs that correspond to the human sera IT50s, we next cloned and expressed the natural blocking mAbs from their original source, i.e., from individuals with high IT50s. First, peripheral blood mononuclear cells were collected from the three volunteer donors with the highest IT50s – namely, individuals 6, 10, and 16 in **Figure 1D**. Their antibody repertoires were then cloned into a phage-display library, and 14 single-chain variable fragment (scFv)-phages were bio-panned and isolated using the 17875/Leb BabA protein as the affinity matrix (denoted as ABbAs (<u>A</u>nti-<u>B</u>a<u>bAs</u>). Six of the clones (43%) had identical sequences, so this mAb was denoted ABbA(1) (**Figure S4A**). To determine if ABbA(1) functionally corresponds to the inhibitory antibodies in serum, two tests were performed with an expressed ABbA(1)-scFv ^24^. First, the ABbA(1)-scFv was found to strongly inhibit Leb binding with an inhibitory concentration 50% (IC50) of 27 ng/mL (**Figure S4B**). Second, ABbA(1)-scFv was found to bind specifically to 17875/Leb bacteria but not to the isogenic *babA*-minus mutants (**Figure S4C**). Thus, ABbA(1) exhibited both the necessary affinity and specificity properties. Then, for improved stability and avidity, ABbA(1)-scFv was expressed as a human IgG1 antibody, denoted ABbA(1)-IgG mAb (and named ABbA). Three tests were performed to determine if ABbA itself demonstrated broadly blocking Ab activity. ABbA was tested against the series of 12 global *H. pylori* strains (from **Figure 2A**). Notably, ABbA reduced both the Leb binding (**Figure 4A**) and the attachment to human gastric mucosa (**Figure 4B**) of the majority (9/12) of world-wide *H. pylori* strains from **Figure 2A** and hence exhibited a broadly blocking activity. Thus, ABbA can protect the mucosa from *H. pylori* attachment. Third, to understand the epitope preferences that might explain the broadly blocking ABbA properties, we applied the semi-native-immunoblot method and found that ABbA bound only to folded structural BabA epitopes and not to denatured linear epitopes. Thus, the ABbA mAb binds BabA similarly to the human (polyclonal) serum antibodies from infected individuals (**Figure 4C *vs*. Figure 1I** and **Figure S1H**). Finally, the efficacy, specificity, and affinity of the IgG mAb was assessed by five different tests. First, the ABbA-IgG efficacy increased ∼15-fold in IC50 concentration compared to ABbA-scFv (**Figure S4B**). Second, ABbA bound specifically to BabA but did not recognize the non-Leb-binding BabB paralog (**Figure S4D**). Third, electron microscopy showed that ABbA bound specifically to BabA on the surface of *H. pylori* bacteria (**Figure S4E**). Fourth, ITC tests demonstrated an affinity (Kd) to BabA of 10 nM (**Figures S4F*i*** and **S4F*ii***). Fifth, SPR tests showed the notably ∼5-fold higher affinity to BabA of 2 nM (**Figure S4G**), which is a valid affinity for mAbs with potential for translational applications. Thus, the ABbA mAb exhibited similar high-affinity binding and specificity for BabA as the polyclonal patient sera with a remarkably efficient broadly blocking activity for worldwide *H. pylori* strains. How then does IT50 relate to the pool of bbAbs similar to ABbA in the blood circulation? Titration of a dilution series of ABbA mixed with sera from an IT50-negative individual showed that an IT50 = 70 equals ∼1 µg/mL of ABbA (**Figure 4D**). Because human sera contains ∼10 mg/mL IgG, the ABbA pool constitutes 1/10,000 part of the total IgG content or ∼5mg mAbs, similar to ABbA out of the ∼50g of total serum IgG in circulation. This series of results suggest that ABbA exhibits similar high-affinity binding for BabA as the polyclonal patient sera and that ABbA shows a remarkably broad binding-blocking and attachment-blocking activity for worldwide *H. pylori* strains.

**Figure 4.**
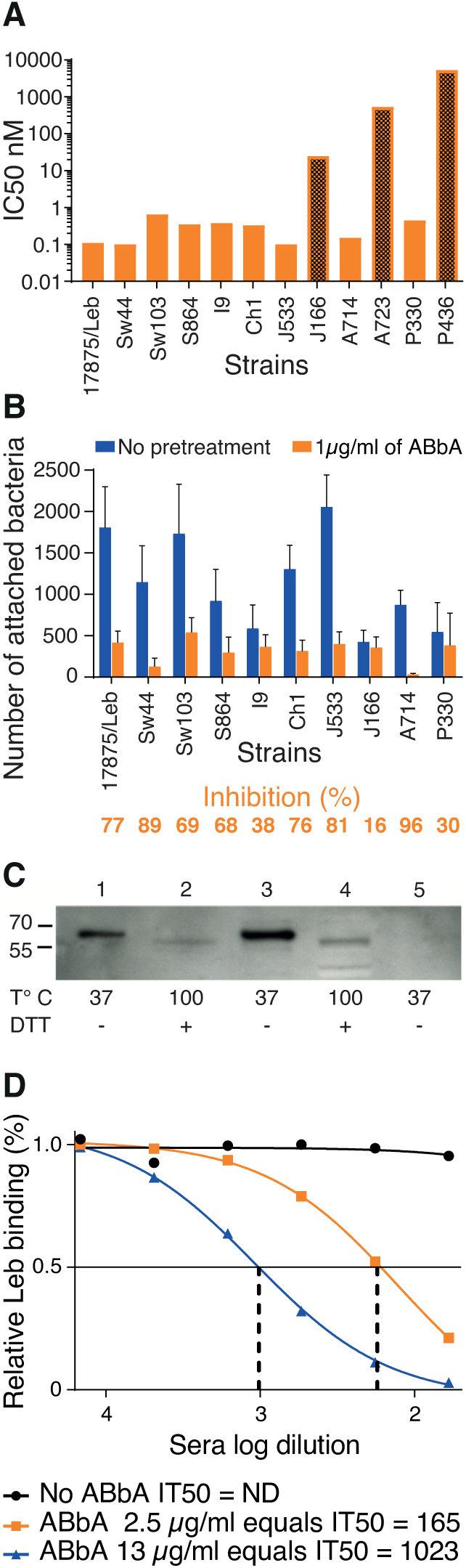
Cloning and characterization of ABbA, the broadly blocking human mAb. (A) Tests of the ABbA IC50 for the 12 world-wide *H. pylori* strains from **Figure 2A**. ABbA blocked Leb binding of 9 strains but was less efficient with J166 (which has low Leb-binding affinity) (**Figure S1D**) (darker bar 1) and did not block binding of the two Indigenous blood group O-binding American Specialist strains A723 and P436 (darker bars 2 and 3) that both exhibit adaptive substitutions in the CBD that is critical for binding to ABO/Leb (**Figure S2C**) ^13, 14, 25^. (B) Test of the blocking of *H. pylori* attachment by ABbA to human gastric mucosa *in vitro* by the series of *H. pylori* strains from (A), except for A723 and P436. The reduction in attachment by ABbA inhibition (in orange) and Inhibition (%) closely reflected the ABbA IC50 for the corresponding strains, where the J166 strain with the higher IC50 was similarly modestly blocked by ABbA in terms of attachment (16%). (C) ABbA-scFv recognized both purified BabA and BabA in size-separated bacterial whole-cell protein extracts (WCPE) under semi-native conditions (Lanes 1 and 3, respectively) but not under denaturing conditions (Lanes 2 and 4, respectively). As a negative control, WCPE of the 17875*babA1A2*-mutant (with no BabA) was applied under semi-native conditions (Lane 5). (D) Tests of ABbA concentration in terms of inhibition activity as defined by IT50 were performed by two dilution series, which showed that 1 µg/mL of ABbA equals IT50 ∼70.

### ABbA performs glycan mimicry in its broadly blocking binding to structural epitopes

The ABbA binding epitope in BabA was identified in five consecutive steps. First, a BabA library was panned for ABbA binding, which identified a minimal 260aa fragment that included the full CBD (**Figures 5A** and **S5A**). Second, for detailed binding epitope mapping, a series of rabbit sera were raised against short peptides that sequentially overlapped with the CBD domain of strain 17875/Leb (**Figure 5B*i***). Only a single serum sample, denoted TB4, successfully outcompeted both the ABbA and the Leb binding (**Figures 5B*ii*** and **5B*iii***, respectively). The dual inhibition ability of TB4 serum suggested that the Leb-binding epitope architecture in the CBD tightly overlaps with the ABbA-binding epitopes because the critical Leb-binding DSS-Triad residues in the CBD Entrance loop are all present in the TB4 peptide and thus constitute binding epitopes for the TB4 serum (**Figure 5B*iv***). However, in contrast to the bbAb activity of ABbA, the TB4 serum only blocked Leb binding of the cognate strain 17875/Leb and poorly inhibited worldwide isolates (**Figure 5A** *vs.* **Figure S5C**). This fundamental difference relates to the high degree of antigenic variation in the Entrance loop (**Figure S5D**). These results argue that human antibody responses against linear epitopes are not efficient in protecting against adaptive microbes that exhibit high mutation and amino acid substitution rates, in contrast to ABbA, which is less sensitive to antigenic variation due to its preference for structural binding epitopes. Third, for identification of the ABbA binding epitope in BabA, the ABbA co-complex crystal structure with BabA was determined to 2.76 Å resolution (Materials & Methods and **Table S5**). The BabA body domain is formed by a 4+3-helix bundle with a four-stranded β-plate insertion that forms the CBD located at the top of the body domain (**Figure 5C*i***). In the CBD, the CL2 cysteine loop and the Key-loop (DL1) and Entrance-loop (DL2) all contribute with binding sites for Leb (**Figures 5C*ii*** and **5D** and **S5E**). The principal determinant of ABO/Leb antigens, the α1.2 secretor fucose, is bound by a pocket formed by the conserved CL2 loop, whereas Leb’s reducing end is bound by the DSS-Triad in the Entrance-loop ^13, 25^ (**Figures 5C*ii*** and **S5F**). In the co-crystal structural model, ABbA binding targets these same three areas where the variable heavy (VH) residue W102 perfectly inserts into the α1.2 secretor fucose loci whereas VH residue L31 resides in a pocket formed by V231 in a position equivalent to the α1.4 Lewis fucose. In addition, VH binds to both D206 and N208 in the Key-loop and to the DSS triad (**Figures 5C*ii*** and **5D**). Thus, ABbA competes with Leb in binding to BabA through functional glycan mimicry and binds the majority of amino acid residues in the CBD that are critical for Leb binding. Notably, ABbA W102 and L31 provide surrogates for the secretor and Lewis fucose by filling the two hydrophobic pockets, glycan mimicry 1 (GM1) and 2 (GM2), respectively, where the hydrophobic determinants of the L-enantiomeric fucose residues reside. Furthermore, in the third glycan mimicry domain (GM3), ABbA binds the DSS-Triad. Fourth, the glycan mimicry property was tested by mutagenesis, where BabA with the DSS-Triad replaced with AAA was expressed in *H. pylori.* The AAA mutant demonstrated complete loss of both ABbA and Leb binding. A single DSS *vs.* D**A**S mutant similarly lost all ABbA and Leb binding (**Figures S5G** and **S5H**).

**Figure 5.**
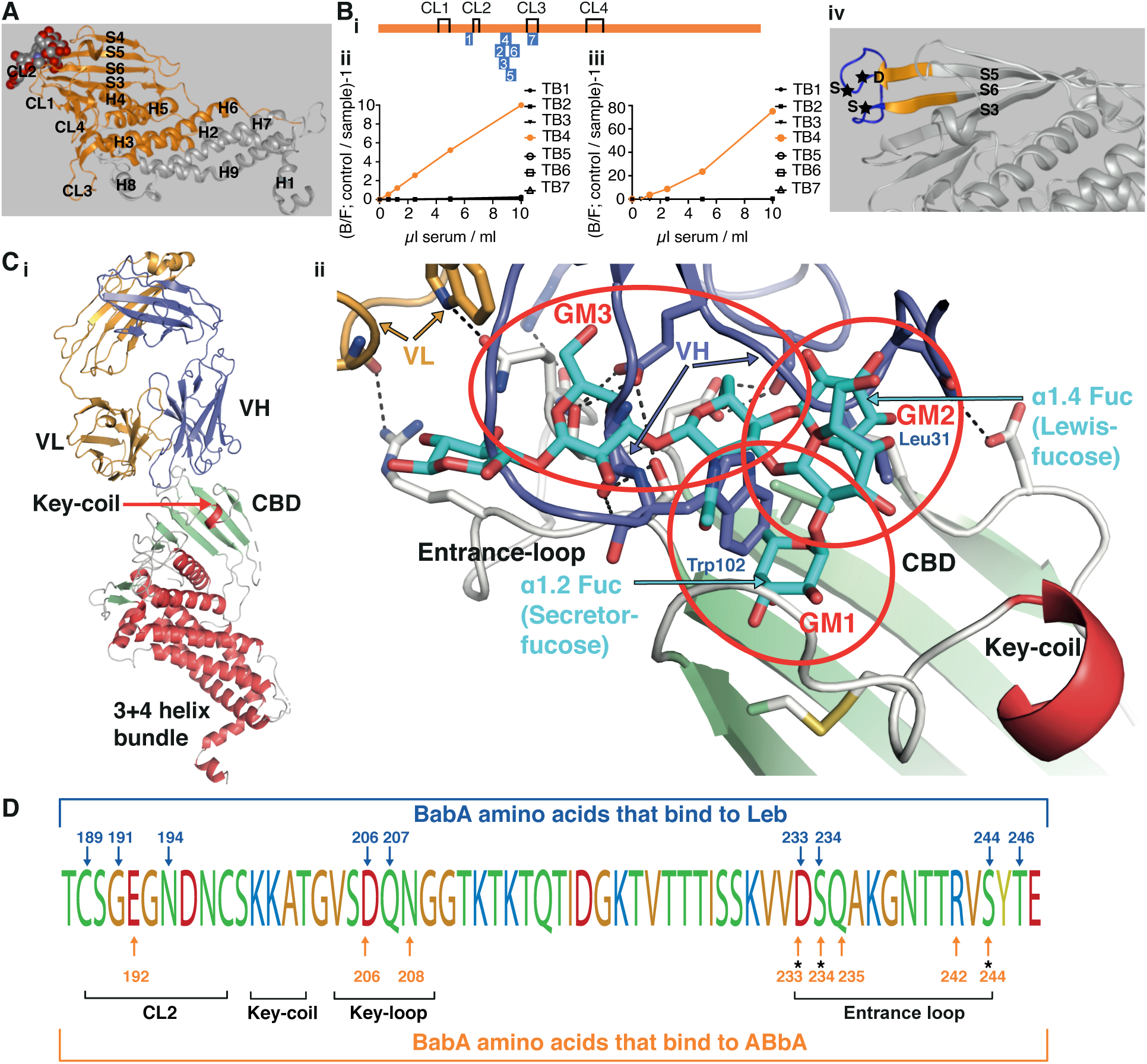
Identification of the structural binding epitope in BabA for the broadly blocking ABbA. (A) The aa76-335 BabA fragment in orange was superimposed on the 460 aa BabA structure (grey) constituting the minimal 260 aa BabA domain required for ABbA binding. The Leb glycan in red/grey spheres indicates the CBD location. (**B***i*) The schematic locations of the seven BabA peptides (TB1-7) derived from strain 17875/Leb used to raise rabbit anti-sera. The four disulfide loops (CL) are indicated for orientation (**Figure S5B**). (**B***ii*) Inhibition of ABbA binding and (**B*iii***) inhibition of Leb binding to 17875/Leb bacterial cells was tested with the seven rabbit sera in the dilution series. (**B***iv*) The TB4 (BabA_230-247_) peptide includes the Entrance loop (blue) to the CBD, where the critical D233-S234-S244 (DSS) residues are indicated by stars, and the proximal parts of the connecting β-strands S5 and S6 are shown in orange. (**C***i*) Co-crystal of ABbA-Fab and BabA with the locations of the ABbA Light (VL) (orange) and Heavy (VH) (lilac) chains. The CBD (green) and the Key-coil (bright red) are located in the extra-cellular domain i.e., the BabA body domain (red). (**C***ii*) The ABbA/BabA co-crystal structure with Leb (turquoise) superimposed in the CBD region, with the Key-coil (in red from C*i*) for orientation. The ABbA-CBD interactions demonstrate three sites of Leb glycan mimicry, including GM1, where ABbA VH (lilac) replaces the α1-2 “Secretor” fucose in the CL2 pocket; GM2, where ABbA VH replaces the α1-4 “Lewis” fucose; and GM3, where ABbA VH binds to the DSS triad residues. (**D**) The 17875/Leb BabA CBD domain with the amino acids positions that bind to Leb and/or to ABbA are indicated by blue vs. orange arrows, respectively.

Thus, the triple-locus glycan mimicry performance of ABbA provides a universal fit to the CBD despite its polymorphic landscape, and this works “hand-in-glove” to protect the ABbA mAb from immune escape and to help explain its broadly blocking properties.

### Functional identification of the world-wide extended ABbA binding epitope

To better understand the global broadly blocking repertoire of BabA epitopes for ABbA, we tested the ABbA efficiency in inhibition of Leb binding with 146 *H. pylori* world-wide clinical isolates. First, the IC50 of ABbA was found to range >100-fold, from efficient inhibition in 48% of strains (IC50s <1 nM) to no inhibition in 18% of the strains (IC50s >100 nM) (**Table S6**). Second, the ABbA binding strength was tested by ^125^I-labeled ABbA and was also found to range 100-fold (**Table S6**). Third, the IC50 and ABbA binding strength were found to strongly correlate (**Figure S6A**). Thus, less ABbA is needed for inhibition of Leb binding of the clinical isolates that exhibit stronger ABbA binding. Fourth, the ABbA and Leb binding strengths were found to correlate among worldwide strains (**Figure S6B**). However, the ABbA binding strength was higher for European, medium for Asian, and lower for the Indigenous American/Latin American *H. pylori* isolates (p < 0.001) (**Figures 6A** and **S6C**). This difference could be a consequence of the phylogenetic distance between the Old and the New World and hence be related to BabA antigenic variation, or it may be a consequence of functional adaptation to the difference in predominant ABO blood group phenotypes. This idea was supported by the finding that the Indigenous American/Latin American blood group O or A specialist binding strains were 7-fold more prevalent in the ABbA low binding strength group (p < 0.001) (**Figure 6B**). Furthermore, 81% of the Indigenous American/Latin American specialist strains were low, whereas 38% were high in ABbA binding strength (**Figure 6C**), i.e., *H. pylori* strains with adaptation to the specialist preference in binding were more prevalent in the ABbA low binding strength group (p < 0.0017). To better understand the global diversity in ABbA glycan mimicry, we aligned the BabA sequences according to their ABbA binding strength (**Table S7**). The alignment showed that low ABbA binding strength was associated with amino acid substitutions located in BabA’s Leb-binding landscape, where charged residues are replaced with small neutral or large hydrophobic amino acids (**Figure 6D** *vs.* **Figure 5D**). The high substitution level in these epitope positions corresponds to their high positive selection activity, i.e., positions where nucleotide mutations preferentially result in amino acid substitutions ^11^. These findings suggest that the lower ABbA binding strength and hence reduced epitope fit in the New World/Americas is a consequence of functional adaptation of the BabA binding preferences to the ABO blood group phenotypes in the local populations. Thus, rapid adaptation in BabA binding preference with critical amino acid residue replacements can lower the binding strength of the bbAbs with subsequent reduced immuno-protection, which is in line with the pandemic GC prevalence in South America ^22^.

**Fig 6.**
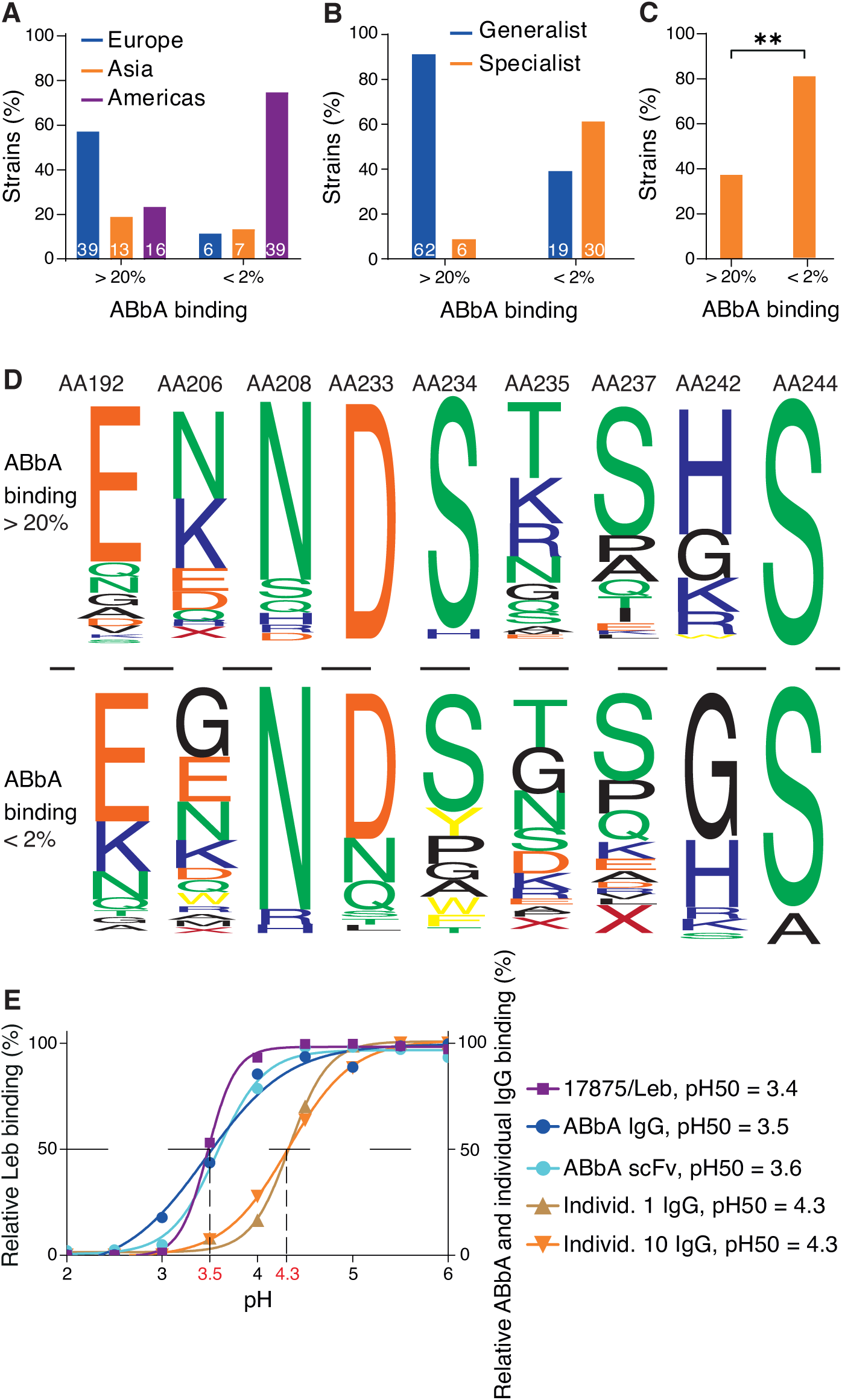
The ABbA global binding epitopes and their acid sensitivity in binding. (B) ABbA binding strength is higher (>20% binding of ^125^I-ABbA) among European isolates, whereas it is lower (<2% binding) among the majority of Indigenous American/Latin American isolates (**Table S6)**, and the Asian isolates are distributed over both groups. (C) Strains with the specialist binding preference are 5-fold more prevalent among strains with low ABbA-binding strength (<2% binding) (**Table S6)**. (D) The great majority of the Indigenous American/Latin American specialist strains have low ABbA binding strength (**Table 6B)**, p<0.0017**. (E) The BabA web logo is based on the alignment of the series of amino acid positions that demonstrate replacements between strains that bind ABbA with high strength (>20%, 62 strains) *vs*. low strength (<2%, 43 strains) (**Table S6**) with a focus on positions E192K, K/N206G, D233N/X, S234G/A/W/Y/X, K/R/Q235D/G, S237E/D/K/R/Q, and K/R/H242G. (F) The acid sensitivity profiles, denoted as pHgrams, were determined by incubation of *H. pylori* with ^125^I-Leb in the pH 2–6 ^14^. The dashed line shows when half of the Leb binding remained, i.e., 17875/Leb pH50 = 3.4. A similar procedure was applied for profiling ABbA-scFv, ABbA-IgG, and Fc-purified IgG from individuals 1 (**Figure 1B**) and 10 (**Figure 1D**) (**Figure S5I**).

### Binding of the bbAbs to *H. pylori* is acid sensitive

In the gastric mucosa, *H. pylori* adherence needs to be synchronized with the desquamation and rapid turnover of the gastric epithelium. Close to the cell surfaces the pH is buffered by the bicarbonate system, but the pH quickly drops in the mucus layer towards the stomach lumen and its extremely acidic gastric juice. In the mucus layer, *H. pylori* adheres to the Leb glycosylated desquamated epithelial cells and large mucins ^26^ until the bacteria reach the critical acidic zone where BabA binding is inactivated and *H. pylori* breaks free from the Leb entanglement. Unusually well adapted to its environment, BabA binding is fully reversible after acid exposure, which allows for the detached *H. pylori* bacteria to return to the buffered environment where they reattach to the epithelium. In other words, *H. pylori* use the pH gradient for recycling of infection and thus are constantly functionally bio-panning for the next generation of fittest *H. pylori* ^14^. Two tests were performed to determine if BabA has adapted acid sensitivity in binding to also evade bbAbs. First, ABbA-IgG and ABbA-scFv were pH titrated for binding, and both showed a pH50 = 3.5 (the pH at which half the maximal ABbA binding was lost). Thus, ABbA binding is most similar in acid sensitivity to Leb binding by 17875/Leb (which also has pH50 = 3.5) (**Figure 6E**) and hence similar to the median pH50 = 3.7 for clinical *H. pylori* isolates ^14^. Second, to characterize the general acid sensitivity in serum antibodies that bind to *H. pylori* bacterial cells, the purified serum IgGs from individuals 1 (**Figures 1A** and **1B**) and 10 (**Figure 1D**) were similarly pH50 titrated. The IgGs from both individual 1 and 10 were almost a full pH unit more acid sensitive in *H. pylori* binding (pH50 = 4.3/4.4) compared to ABbA (**Figure 6E**). The glycan mimicry performed by ABbA constitutes the most plausible explanation for the identical acid resistance in Leb and ABbA binding, with the “hand-in-glove-fit” to critical Leb binding residues such as the DSS-triad in the CBD with the aspartic acid (D) responding quickly to pH changes due to its acidic pKa ∼3.5.

## DISCUSSION

Human responses to *H. pylori* infection are well recognized to raise high levels of humoral and cell-mediated immunity, and although these antibodies form the basis for the clinical diagnostics, the immune responses fail to eradicate the infection and also fail to protect against reinfection after antibiotic treatment. *H. pylori* is inherited over the generations from our parents, i.e., through vertical dissemination in contrast to horizontal pandemics, and thus is persistent over the lifespan. The chronic outcome is understandable considering that *H. pylori*, over the millennia of human infections, has established multiple mechanisms to evade our immune responses through fast and confluent adaptation by high mutation rates and antigenic variation ^27^ combined with dysregulation of T-cell responses (reviewed in ^28^). The many escape routes are reflected in both the shortcomings of conventional *H. pylori* vaccine development ^29^ and the high re-infection rate in many parts of the world after eradication of infection with antibiotics ^30, 31^. Fortunately, our results demonstrate that in the vast majority of *H. pylori-*infected individuals the humoral immune system responds with broadly blocking antibodies (bbAbs) (**Figures 1** and **S1**) that target conserved structural epitopes in the Leb-binding domain in the otherwise highly polymorphic BabA adhesin. By doing so, the bbAbs perform functional glycan mimicry and can reduce *H. pylori* mucosal adherence by competing with and weakening the BabA-mediated Leb binding (**Figure 5C**). Glycan mimicry has been suggested for mAbs that compete with the sialic acid mono-saccharide residue of the influenza haemagglutinin ligand ^32^. In comparison, ABbA takes full advantage of the glycan mimicry strategy and performs mimicry of the two critical Leb-fucose residues located in the bottom and the wall of the CBD and, in addition, mimics the lactose in the Leb-core chain in the upper part of the CBD. Thus, ABbA performs “wall-to-wall” triple-locus glycan mimicry, and this strategy allows ABbA to function despite numerous amino acid substitutions in BabA over the lifetime of the individual. As a proof of concept in this work, the triple glycan mimicry enabled ABbA to block most of the world-wide *H. pylori* strains despite the unprecedented high level of polymorphism in BabA. Notably, this glycan mimicry mechanism that blocks mucosal adherence does not constitute an Achilles heel for *H. pylori* infections *per se* because our immune system nevertheless does not manage to eradicate the *H. pylori* infection. Similarly, the ∼30% of *H. pylori* infections in Western-Europe and the United States among individuals with predominant non-atrophic gastritis, where *H. pylori* does not express BabA and does not bind Leb, are still persistent over the lifespan ^7, 10, 33^. However, the glycan mimicry of bbAbs can suppress the adherent-inflammatory lifestyle of *H. pylori* among those individuals who carry the triple-positive *H. pylori* infection, which express BabA/VacA/CagA and are highly associated with DU and GC. In this way, and without being able to eradicate the infectious burden, the humoral immune system can constrain the mucosal inflammatory stress of the *H. pylori* infection, resulting in a more balanced and “commensal” level of bacteria, and importantly, with a resultant lowered risk for severe gastric disease. But how about the GI-protective IgA? Perhaps surprisingly, there is no increased prevalence of severe gastric disease/cancer in IgA-deficient individuals (which is a common affliction affecting about 1/800 individuals), and this argues that IgA and/or secretory IgA are not major intestinal immune protectors against chronic inflammatory *H. pylori* infection ^34, 35^. Humoral bbAb mediated protection is further illustrated by the elevated IT50 at older age (**Figure 1F**), which might be a compensation for the reduced T-cell reactivity seen in elderly people ^36, 37^. In addition, we show that individuals with low IT50 are at higher risk for DU, which suggests that even though the etiology of DU is the *H. pylori* infection, the cause of DU is a consequence of suboptimal humoral responses, i.e., DU is an immunodeficiency disease with insufficiently induced levels of protective bbAbs. The risk for DU is also increased in this group because they often harbor *H. pylori* with greater binding strength with consequential unbalanced and augmented inflammation pressure. It is noteworthy that the higher binding affinities among DU strains might be a consequence of the low and hence less protective IT50 levels among individuals who develop DU disease.

We have previously shown that Leb-binding at buffered pH provides for tight adherence to the gastric epithelium ^14^, which allows for access to essential nutrients and iron ^38, 39^. The many decades of unsuccessful development of *H. pylori* vaccines based on conventional antigen compositions can be better understood by our new results on how *H. pylori* makes use of the pH gradient in the mucus layer for a daily “acid wash cleaning” in order to rid itself of antibodies and acid-sensitive complement factors. After which, *H. pylori* can return, fully “immuno-recycled”, to its protected niche in the epithelial lining, as illustrated in Video S1 (https://play.umu.se/media/t/0_bf1lh9gt). Our results for the protective bbAbs suggest translational applications for non-antibiotic treatment regimens based on therapeutic bbAbs such as ABbA and predictive diagnostics such as IT50 for individuals who are at risk for severe gastric disease.

## ACKNOWLEDGEMENTS

We thank T. Ny for valuable discussions; The Biochemical Imaging Centre Umeå (BICU) within the National Microscopy Infrastructure (NMI); The Protein Expertise Platform (PEP) at Umeå University; S. Dübel for providing plasmids. We thank Ö. Furberg (<u>NoPolo.se</u>) for the digital movie and illustration and M. Borén and E. Morrow for the figure work. This research work was supported by the Swedish Medical Research Council (VR) (2017-02183), Cancerfonden (CF) (CAN 2018/807, and 21/1875), the Umeå University Biotechnology Fund, the J.C. Kempe and Seth M. Kempe Memorial Foundation to TB, and the Erling-Persson Family Foundation to TB, LH, and RMo, and was in part performed within The Ukrainian-Swedish Research Center SUMEYA. The authors gratefully acknowledge the financial support of the ERASMUS program (EU funding) for PhD student exchange within the collaboration between Umeå University and Sumy State University (grant 2017-1-SE01-KA107-034386). The work was also supported by the VR (2019-01598) and the CF (20 0964 PjF) and Region Västerbotten (ALF) to ML; the VR, the Swedish Society of Medicine, the Norrbotten County Council, Astra Zeneca R&D, the Orion Research Foundation, and the Finnish Medical Foundation to JR; the US NIH (R21AI140038) to DSM; and the Science Foundation Ireland Awards 16/RC/3889 and 13/IA/1959 to SO. Structural studies were supported by the Flanders Research Foundation (FWO; grants G033717N and G1521517N). We acknowledge access to the ESRF and are grateful to the staff at the ID29 beamline.

## AUTHOR CONTRIBUTIONS

All authors contributed substantially to the work and interpretations of the data. J.A.B., K.M., A.P., A.Sc., J.O.E., G.L., K.B., Y.A.C., K.T., I.T., O.S., I.V., A.B., P.S., R.C., D.X.J., H.M., R.S., A.Sh., G.B., L.R., J.G., R. Ma., F.G., C.A.R., A. Lu., R.H., H.R., R. Mo. performed the research and together with L.H. and T.B. analyzed the data. M.M.D’E., L.A., J.R., P.A., L.E., D.Y.G., V.Kac., A.M., S.C., B.C.K., S.P., O.K. M.C., J.T., J.M.W., D.S.M., Q.P.H., A. Lo., I.V., and O.S. provided serum samples and *H. pylori* strains. SO performed the synthesis of Leb. KH performed the EM. O.B., V.Kas., A. Lo., V.I., M.A.A., and M.L. supervised the research. J.A.B., K.M., A.P., A.Sc., J.O.E., K.T., H.M., A.Lu., H.R., D.E.B., R.Mo., L.H., and T.B. wrote the paper. L.H. and T.B. designed the research. All authors reviewed the manuscript.

## Abbreviations

scFv: single chain fragment variable
mAb: monoclonal antibody
bbAb: broadly blocking antibody
Rf: Risk factor
CBD: carbohydrate binding domain
GM: glycan mimicry
DU: duodenal ulcer disease
GA: gastritis.

## STAR * METHODS

### KEY RESOURCES TABLE

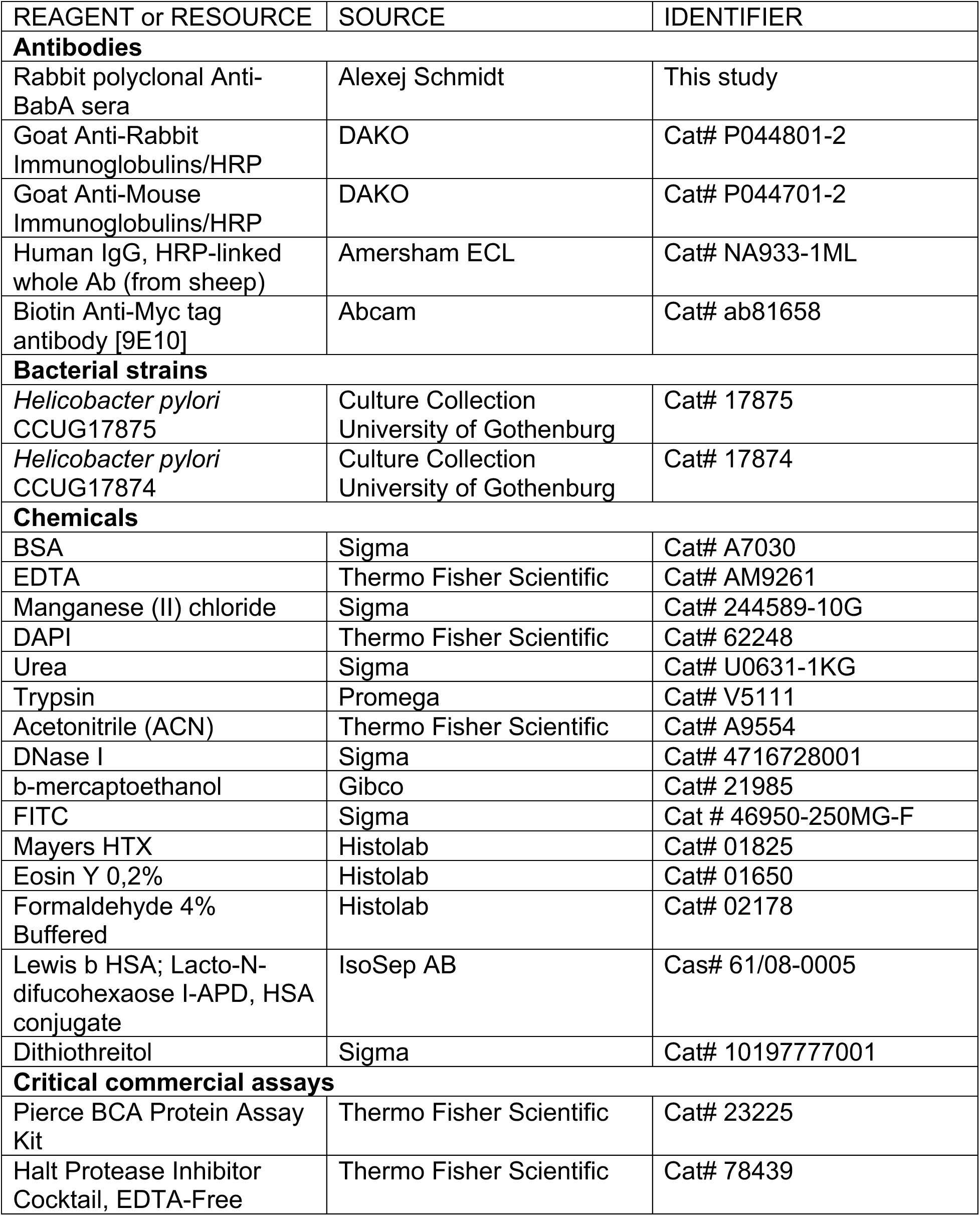
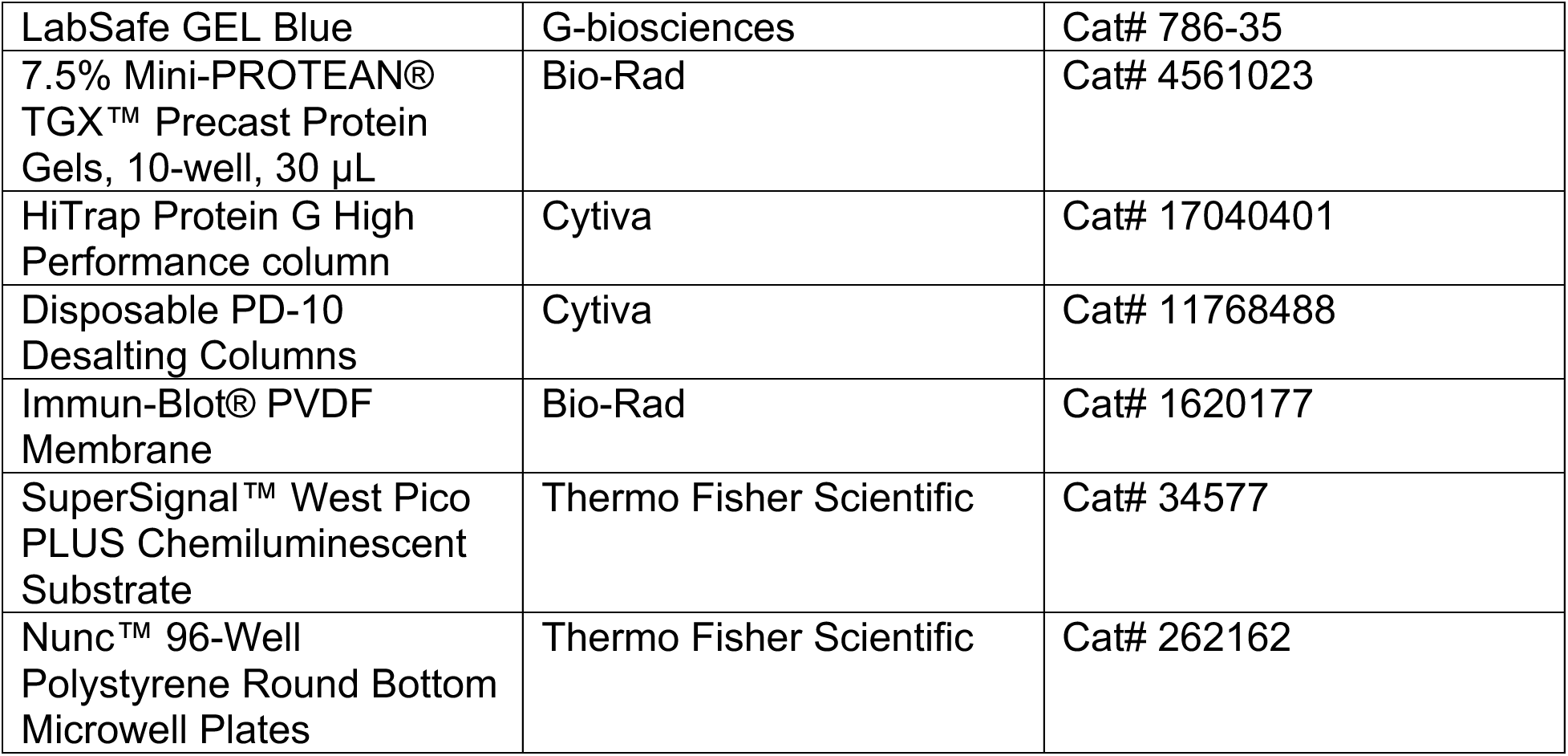

### RESOURCE AVAILABILITY

#### Lead contact

Further information and requests for resources and reagents should be directed to and will be fulfilled by the lead contact, Thomas Borén.

#### Materials availability

Strains and plasmids generated in this study are available upon request to the lead contact, Thomas Borén.

#### Data and code availability

Genome sequencing data for *H. pylori* CCUG 17875 and its plasmid are available from GenBank with the accession number CP090367 and CP0903678, respectively. The ABbA-BabA co-crystal coordinates are available by the Accession code: PDB ID 7ZQT. All other data are available in the main text or the supplemental information. Any additional information required to reanalyze the data reported in this paper is available from the lead contact upon request.

### EXPERIMENTAL MODEL AND SUBJECT DETAILS

The Supplementary Materials and Methods includes the following topics:

▪ *H. pylori* Strains and Media
▪ Blood Group Antigens and Conjugates
▪ Serum Samples: Human sera collection, poly clonal rabbit sera, mice sera of infected and vaccinated FVB/n mice, and histological tests of Leb mouse gastric mucosa.
▪ Monoclonal ABbA-IgG
▪ *In situ* binding by *H. pylori In situ* binding of *H. pylori* to human gastric mucosa histo-tissue sections, and inhibition of *in situ H. pylori* binding by human sera and by ABbA-IgG.
▪ Radio-immuno Analysis (RIA): BabA binding properties, inhibition titers (IT50s), ABbA-Ig inhibition capacity (IC50), and analysis of ^125^I-ABbA and Leb competitive RIA.
▪ SDS-PAGE and Immunoblot detection.
▪ Isolation, purification, and expression of ABbA-IgG: cDNA synthesis and PCR of human variable regions, ScFv-library construction, selection of BabA-binding scFvs by phage display, and construction of a human IgG; ABbA-IgG binding properties by isothermal titration calorimetry (ITC) and surface plasmon resonance (SPR); immuno-EM and detection of BabA by ABbA; construction of a BabA phage library; and selection of ABbA-IgG-binding BabA fragments.
▪ The BabA protein co-crystallized with ABbA-IgG (Fab).
▪ Genetic complementation by shuttle vector-expressed recombinant BabA and DSS mutants in *H. pylori*.
▪ Amplification and analysis of BabA sequences.
▪ Amino acid positions in the BabA CBD that modulate ABbA binding strength.
▪ Acid sensitivity in Leb-binding and ABbA binding.

#### 1. *H. pylori* strains

##### Laboratory strains

*H. pylori* 17875/Leb is an isolated single clone of *H. pylori* CCUG17875, and it binds to ABO/Leb antigens with high Leb-binding affinity, but not to sialylated antigens ^41^. The 17875*babA1A2* strain is a null mutant with the two *babA1* and *babA2* genes deleted from *H. pylori* CCUG17875, referred to as the *babA1A2*-mutant ^10^. The *H. pylori* CCUG17874 binds sialylated antigens but not the ABO/Leb antigens ^42^.

##### Clinical *H. pylori* isolates

The *H. pylori* isolates from Sweden (29 strains), Germany (4 strains), Spain (13 strains), Japan (13 strains), Alaska (11 strains), and Peru (38 strains) have been previously described ^11^. The 13 Indian strains

The H. pylori isolates Sw44, Sw103, S864, J533, A714, A723, P330, P436 have been previously described ^11^, and the 13 Indian strains including isolate I9 and the J166 strain have been described in ^14^ and ^43^, respectively. The 32 (**Table S2B**) Mexican isolates from the UMAE Pediatria IMSS in Mexico City were isolated from patients with GA (n = 13) or DU (n = 13). The *H. pylori* Ch1 strain was isolated from a Chinese individual with a family history of gastric cancer. *H. pylori* USU101 and a Rhesus macaque passage clone USU101Δ*babA* with a natural *babA* deletion were as described ^14, 44^.

#### 2. *H. pylori* culture media

Cultures were performed with blood agar plates, and *H. pylori* cultures were grown in a mixed-gas incubator under micro-aerophilic conditions as described ^42^.

#### 3. Blood group antigen conjugates

Two different types of fucosylated blood group antigen conjugates were used, namely 1) semi-synthetic Leb and ALeb-glycoconjugates (Leb-HSA and ALeb-HSA, respectively) with natural purified oligosaccharides covalently linked to human serum albumin (Isosep AB, Tullinge, Sweden) or 2) oligosaccharides synthesized by S. Oscarson (co-author) and conjugated to HSA. The conjugates were used for the radio-immuno assay (RIA) binding experiments.

#### 4. Serum samples

##### 4.1. Human subjects and sera

Procedures involving human subjects were approved by the following boards. The series of sera from the Karolinska Institutet Hospital was approved by ethical permits Dnr 185/93 and 2013/1255. The Kalixanda study was approved by the Umeå University ethics committee permit Dno. 98-99, §156/98. The series of adult sera from Sumy, Ukraine, was approved by the local ethics committee of Sumy Council “Sumy Regional Clinical Hospital” permit protocol 2/5, 15 February 2019. The series of pediatric sera from Sumy, Ukraine, were approved by ethical permit protocol №11/1 of the Institutional Bioethics Committee of the Medical Institute of Sumy State University from 26 November, 2020. The series of serum samples from Baylor College of Medicine were made anonymous and approved by the IRB at Baylor College of Medicine. The series of serum samples from Mexico City was approved by ethical permit IMSS, Mexico; 2008-785-001. Informed consent was obtained from all human participants or legal guardians of participating minors.

###### 4.1.1. Karolinska Institute (KI) series

Serum samples from volunteer donors were tested by *H. pylori* ELISA ^34^, and 36/72 (50%) were positive (**Table S1A**). Additional follow-up serum samples were collected four times for individual 16 and three times for individual patient 36 over ∼30 years **(Figure 1D).**

###### 4.1.2. Kalixanda series

A total of 322 sera samples were obtained from volunteers, and ELISA analysis demonstrated that all sera were *H. pylori* positive with individual signal intensity, and 74% of the samples were CagA positive ^20^. Of these, 161 samples were collected from men with a median age of 59 years and 161 samples were collected from women with a median age of 61 years (**Table S1B**).

###### 4.1.3. Ukraine, adult series

A total of 104 sera samples were collected from patients with gastric diseases at Sumy Regional Clinical Hospital. The sera were tested by ELISA, and 79 of them (76%) were defined as *H. pylori* positive. The 20 men and 59 women were from 20 to 79 years of age (**Table S1C**).

###### 4.1.4. Ukraine, children series

A total of 135 serum samples were obtained from a children’s sera collection at Regional Children’s Hospital and St. Zinaida City Children’s Hospital in Sumy, Ukraine. The patients were tested by ELISA, and 36 of them (27%) were defined as *H. pylori* positive. The 14 girls and 22 boys (39% girls and 61% boys) were from 3 to 17 years of age **(Table S1F**).

The Ukrainian sera samples were analyzed with an anti-*H. pylori* ELISA (IgG) test system (EUROIMMUN, a PerkinElmer company, Medizinische Labordiagnostika AG, Lübeck, Germany).

###### 4.1.5. Mexico series 1

A total of 200 ELISA-positive serum samples from patients with gastric disease were obtained from Mexico UMAE Pediatria. Of these, 200 *H. pylori* ELISA-positive serum samples were tested. The patient group included 82 men from 30 to 81 years of age and 118 women from 30 to 86 years of age (**Table S1E**).

###### 4.1.6. Mexico series 2

A total of 79 serum samples from patients with NAG or DU diagnosis were obtained from Mexico UMAE Pediatria from patients with defined gastric disease. Among these, 35 patients had NAG, including 19 men (33–82 years of age) and 16 women (30–79 years of age), and 44 patients had DU, including 26 men (38–86 years of age) and 18 women (27–79 years of age) (**Table S3**).

###### 4.1.7. The US series

A total of 141 *H. pylori* ELISA-positive serum samples from patients with gastric disease at Baylor College of Medicine (Houston, TX, USA) were subjected to further investigation (**Table S1D**).

#### Sera and antibodies

##### 4.2. Sera from rabbits immunized with synthesized peptides (**Figure 5**)

For epitope mapping, rabbit sera were raised against 7 peptides located in the BabA CBD (**Figure S5B**). Peptides were synthesized, and Cys residues were added to peptides 1–6 for keyhole limpet hemocyanin conjugation and used in four consecutive subcutaneous immunizations (Agrisera AB, Vännäs, Sweden). Each rabbit serum sample was quality tested and shown by immunoblot to bind BabA.

##### 4.3. Polyclonal rabbit sera for immunoblot detection of BabA and BabB (Figure S3G and Figure S4E*iii* and *iv* and Figure S5F)

BabA and BabB proteins were detected with the polyclonal anti-BabA VITE antibody and VIRA antibody, respectively, diluted 1:6000 and secondary HRP-goat antibody diluted 1:1000 (DakoCytomation, Denmark A/S) according to ^14^.

##### 4.4. Monoclonal ABbA-IgG (Figure 4, 5, and 6)

Monoclonal ABbA-IgG was initially expressed and produced in Schneider cells ^45^ and later by Absolute Antibody Ltd., Oxford Center for Innovation (New Road, Oxford, 0X1 1BY, United Kingdom) using transient expression in HEK293 cells.

#### 5 Blocking buffer and blocking solution

“Blocking buffer” contained 1% BSA (SCA Cohn fraction V, SWAB, Sweden) in PBS-Tween = PBS-T (PBS, pH 7.4, 0.05% (v/v) Tween-20). For preparation of “SIA-buffer”, 1% BSA was oxidized by 10 mM sodium periodate in 0.1 M acetic acid buffer (pH 4.5) for 1 h. The reaction was stopped by incubating for 30 min in 20 mM sodium bisulfite and 0.125 M potassium phosphate. The solution was then dialyzed against deionized water overnight at 4°C. For SIA blocking buffer, the periodate-treated BSA was mixed with PBS-T. For SIA blocking solution, the periodate-treated BSA was mixed with 150 mM sodium chloride/0.05% Tween. The two different preparations were then filtered through 0.22 µm filters, aliquoted, and kept at –20°C.

### THE FOLLOWING METHODS ARE PRESENTED IN EXPERIMENTAL ORDER AND ACCORDING TO THE SECTION HEADINGS

#### 1. *In situ* binding of *H. pylori* to human gastric mucosa histo-tissue sections (Figure S1A)

Bacteria were labelled with fluorescein isothiocyanate (FITC) (Sigma, St. Louis, MO), and *in vitro* bacterial adhesion was tested as described ^42^. Slides were mounted with DAKO fluorescent mounting media (DAKO North America, Inc., CA, USA) and imaged with a Zeiss Axio Imager Z1 microscope (Carl Zeiss AB, Stockholm, Sweden) or the Leica Thunder microscopes. FITC-labelled *H. pylori* were digitalized from at least 3 identical fields of view from serial sections to quantify bacterial attachment. All images were evaluated by ImageJ. Briefly, micro-picts were subjected to the threshold in order to separate objects of interest from the background and quantified by “Analyze particles”. Graphs and curves were created using GraphPad Prism 9 (San Diego, USA)

##### 1.1. Inhibition of *in situ H. pylori* binding by human sera (**Figure 1A** and S1B)

Inhibition tests were performed with the *in situ* binding methods as described above with the following modifications. FITC-labeled bacteria were first mixed with sera in SIA buffer at the dilutions shown in the figures on a slowly rocking table for 1 h at room temperature and then processed as described ^42^. The attachment of bacterial cells was digitalized and quantified as described above.

##### 1.2. Inhibition of *in situ H. pylori* binding by ABbA-IgG (**Figure 4B**)

Inhibition of bacterial attachment by ABbA-IgG was carried out as described above with slight modifications. FITC-labeled bacteria were first mixed with different dilutions of ABbA-IgG diluted in SIA buffer, incubated on a rocking table for 2 h at room temperature, and then processed, digitalized, and quantified as described above.

#### 2. Tests of inhibition titer (IT50) and of ABbA IC50 and binding affinity by RIA and ITC and SPR

##### 2.1. Labeling of Leb-HSA and ALeb-HSA by ^125^I

The Leb-HSA and ALeb-HSA conjugates (IsoSep AB, Tullinge, Sweden) were I^125^-labeled (I^125^-Leb-conjugate) using the chloramine-T method ^42^.

##### 2.2. Labeling of human sera IgG and ABbA-IgG by ^125^I

Labeling of human sera IgG and monoclonal ABbA-IgG was done according to the chloramine-T method with slight modifications. Briefly, 1 µl of ^125^I (PerkinElmer, Waltham, MA, USA) was added to 2 µg of sera/ABbA-IgG in 50 µl K_2_HPO_4_/KH_2_PO_4_ buffer (pH 7.4). Labeling was started by the addition of 20 μL 0.6 mg/mL chloramine-T and stopped by the addition of 100 μL 1 mg/mL Na_2_S_2_O_5_ and 200 μL KI (9.6 mg/mL) after 20 and 40 seconds, respectively. ^125^I-labeled sera/ABbA-IgG was purified with a PD-10 column (GE Healthcare, Sweden), and the fractions with the highest radioactivity were pooled.

The stability of the labeled IgGs was determined by radioimmune binding assay of bacterial suspensions with the target proteins present in excess. Typical binding for the radiolabeled monoclonal IgG was greater than 90% towards the 17875/Leb strain. The procedure was repeated on multiple days post labeling in order to monitor the rate of degradation due to the harsh conditions of the labeling protocol. All experiments with radiolabeled IgGs were performed within 5 days after labeling.

##### 2.3. Analysis of BabA binding properties by RIA

^125^I-labeled Leb-HSA conjugate (“hot conjugate”) or I^125^-labeled Leb-HSA diluted with unlabeled conjugate (“cocktail”) ^42^ was mixed with 1 mL of bacterial suspension (OD_600_ = 0.1) in blocking buffer. Following incubation, the bacteria were pelleted by centrifugation at 13,000 × *g*, and the ^125^I in the pellet and in the supernatant was measured using the 2470 Wizard^2^ Automatic Gamma counter (PerkinElmer, Waltham, MA, USA) giving a measure of binding activity (% binding).

##### 2.4. Analysis of sera inhibition titers (IT_50_) by RIA

Serum samples from 960 *H. pylori*- infected patients were tested for their ability to inhibit the binding of radiolabeled Leb-HSA conjugate (I^125^-Leb-conjugate) to selected *H. pylori* strains. In order to compare IT50s fairly between strains with varying maximum binding properties, all strains were calibrated in a pilot experiment to find the dilution corresponding to 10% I^125^-Leb-conjugate binding. To ensure consistency of bacterial numbers and to aid pellet recovery, strains were diluted with *H. pylori* 17874, which does not bind Leb. For example, 17875/Leb at an OD_600nm_ = 1.0 (2.5 x 10^9^ CFU/mL) was diluted 1:900 with *H. pylori* 17874 to reach 10% binding. Serial dilutions of the serum were made in blocking buffer, and 50 µL of I^125^-Leb conjugate (0.01 ng/µL) was added to a final volume of 500 µL. After addition of 500 µL of the 17875/Leb and 17874 mixture, the tubes were rotated for 17 h at room temperature. Samples were centrifuged (13,000 x *g* for 13 min), and the I^125^-Leb-conjugates in the pellet and supernatant were measured to ascertain the bound and free conjugate amounts, respectively. The relative titer of the tested serum was defined as the dilution titer sufficient to reduce Leb binding to half the maximum value as determined by binding of the Leb conjugate in the absence of serum (IT50).

A micro-scale assay was established in a conical polystyrene 96-well plate (Thermo Scientific, Germany) that allowed the determination of the inhibitory titer with only 3 µL of serum. For each serum sample, a 1:3 serial dilution series (starting from 1:33) was established in 6 wells of the 96-well plate with a final volume of 60 µL. Then, 60 µL was added to each well of a mixture consisting of 1 ng of I^125^-Leb conjugate and a suspension of *H. pylori* 17875/Leb (OD_600_ = 0.1) as well as *H. pylori* 17874 (OD_600_ = 0.1). All samples were diluted in blocking buffer. The 96-well plate was rotated for 17 h at room temperature and then centrifuged (4,000 x *g* for 5 min), and the I^125^-Leb-conjugate was measured separately in the pellet and supernatant to determine the IT50.

This micro-scale assay was additionally adapted to be performed like the original assay in 1.5 mL Eppendorf tubes. The procedure was identical to the 96-well assay with the difference that the centrifugation conditions post incubation were done according to the original assay at 13,000 x *g* for 13 min. IT50s were calculated identically as for the previous assay variations.

##### 2.5. Analysis of ABbA-IgG inhibition capacity (IC50)

The ABbA-IgG and ABbA-scFv inhibition capacities were measured as described in Section 2.2.4 with slight modification. The known concentration of ABbA-IgG was used to calculate the IC50, i.e., the concentration of antibody that provides 50% inhibition of *H. pylori* binding.

##### 2.6. Analysis of ABbA-IgG binding properties by ITC and SPR (**Figures S4E** and S4F)

**ITC** was performed on an Auto-ITC200 from Malvern Panalytical (Malvern, UK). Recombinant BabA from *H. pylori* 17875/Leb was stabilized by adding hexa-lysine and myc-tag to the BabA protein C-terminus as described previously ^25^. Experiments were performed at 25°C in PBS with 10 µM of BabA and injection of 50 µM ABbA-IgG. The titrations were repeated three times with high feedback and a filter period of 5 s. For each experiment, 19 automated injections of 2 μL each were performed (duration 0.8 s) with 300 s intervals between each injection with a stirring speed of 1000 rpm. Calorimetric data were plotted and fitted using a single-site binding model with MicroCal PEAQ-ITC Analysis from Malvern Panalytical (Malvern, UK).

**SPR** experiments were performed on a Biacore 3000 system (GE Healthcare, Uppsala, Sweden). ABbA-IgG was immobilized on a CM5 chip at around 50 RU levels by standard amine coupling at pH 5. Recombinant BabA from *H. pylori* 17875/Leb was injected for 2 min over the surface at concentrations of 66 nM or 40 nM at two- or three-fold dilutions in PBS + 0.005% Tween-20 at 25°C. Regeneration was done by injection of 10 mM glycine (pH 1.7) for 18 s. Dissociation and rate constants were determined by global fitting with Scrubber2 software (Biological Software, Australia). The experiment was repeated three times and presented as the average value.

##### 2.7. Analysis of ^125^I-ABbA and Leb competitive RIA (**Figures 5B*ii*** and **5B*iii***)

For competitive RIA assays, the ABbA-IgG was labeled as described in Section 2.2.2. To measure whether ABbA-IgG could compete with the anti-peptide rabbit serum for BabA- binding, I^125^-ABbA-IgG1 (0.13 nM, ∼20.000 cpm) was incubated with titrated amounts of rabbit serum (1:100 to 1:1600) with constant amounts of *H. pylori* 17875/Leb (OD_600nm_ = 0.002) and *H. pylori* 17874 (OD_600nm_ = 0.1). After overnight incubation at 4°C, the bacteria were pelleted, and γ-emission was measured in the supernatant (free fraction of I^125^- ABbA-IgG1) and pellet (bound fraction). For measuring Leb competition, 1 μg/mL radiolabeled Leb (∼25.000 cpm) was added instead of I^125^-ABbA-IgG1. Dilutions were made in blocking buffer (1% BSA in T-PBS).

##### 2.8. Analysis of ABbA-IgG inhibition capacity corresponding to IT50 (**Fig 4D**)

Given that the antibodies in the patient serums contain antibodies with a binding efficiency on the same level as the ABbA IgG mAb, we devised the term “ABbA IgG-like antibodies” in order to assign more quantitative values and to more understandably talk about the relationship between IT50 (unknown concentration) and IC50 (known concentration). For example, “What level/concentration of “ABbA like IgG” would a patient have if their IT50 corresponds to 100?” Given that the IC50 of ABbA is 12.86 ng/mL, and that we know that this concentration corresponds to the same level of binding as the IT50, we calculate in reverse to find the stock concentration of “ABbA like IgGs” in the serum:

100× dilution = 12.86 ng/mL

1× dilution = 12.86 ng/mL * 100

= 1286 ng/mL OR 1.286 µg/mL

This is the likely concentration of “ABbA like IgGs” in a patient’s serum that has an IT50 of 100.

##### 2.9 Analysis of odds ratios (ORs) by sliding window calculation (Figures 3C and S3E)

We started out by creating a database containing 4 entries derived from the milder symptoms of the infection (gastritis, G) and the more severe symptom of the infection (duodenal ulcer, DU). Both of these are further divided into High or Low depending on the selected cut-off value for the OR calculations.

The measurable parameter is the patient serum IT50 level, i.e. “How many times we can dilute the serum and inhibit 50% of the binding”. Depending on the test-strain, the IT50 for a given serum will change depending on strain’s Leb-binding affinity (see IT50 method). Because there is currently no definition of what a High or Low IT50 is in terms of antibodies towards the *H. pylori* binding adhesin, we want nature to tell us “is there a numerical value of these IT50 for any given strain that indicates whether an individual patient is at risk for more severe disease progression?”

We had 60 patients evenly/randomly divided between the DU and G diagnoses. These patients had serum IT50s tested against multiple strains of different affinities for the Leb conjugate. For example, with the CCUG 17875/Leb strain (high affinity) we had IT50s ranging from 1 (serums that inhibits, but does not reach the 50% binding mark) and up to a few hundred. The mean and median IT50 was around 75. So, we started testing the cut-off values at 2, and then slid our way up towards 150.

For every step we created an entry in the database. The first step was the temporarily defined “cut-off” (in this first entry it is equal to 2). The script then counted all observations of patient with the G diagnoses that had an IT50 below the current cut-off into the “Low G group”, everything with a G diagnose and above the cut-off was sorted into the “High G group”, and the same procedure was applied for the sera samples of DU diagnoses. For each tested cut-off we were then left with 4 values that told us how the patients were distributed among the 4 groups, and a row-name was created corresponding to the cut-off it was calculated for.

The slider increased 1 more step and it iterated itself and created another entry for the cut-off equal to 3, and then continued to do so until we ran across all of the different cut-offs we needed assessed by that particular test-strain. Because a sample could only move from the Low group to the High group, once a sample had moved into the High group it could never end up in another group again. So, after a certain time there were very few samples left that could “move” into another group. This means that we always reached a point where there was no point in checking above a certain value as things would change with very low frequencies.

The final Table was then comprised of rows containing the cut-off value and the patient distribution among the 4 groups.

The OR was then calculated by dividing the ratio of “low DU/high DU” by the ratio of “low G/High G”. If the OR was equal to or close 1 this meant that there was no difference between the 2 different diagnoses and the patients’; IT50s. If the OR was greater than 1 this indicated that a high IT50 was connected to a milder diagnosis (G), and if the OR was less than 1 this indicated that a high IT50 was connected to a worse diagnosis (DU).

For each of the different cut-offs, we ran a Fisher’s exact t-test to test for significance, because low and high ORs did not necessarily correlate with significant observations.

The ORs across the whole tested range for each strain was illustrated in a window containing all the ORs and how the OR changed as the cut-off level increased. On the second (right) Y-axis, the corresponding significance from the ORs was plotted. The classical 0.05 and 0.01 p-values were plotted as horizontal lines to visually show where (if anywhere) the ORs were presented as significant.

At these points we suggest that these cut-offs potentially could be used to indicate whether a patient should seek eradication (or future vaccine) treatment of their *H. pylori* infection or whether their own immune system will most likely be able to keep the infection in balance i.e., without disease progression.

### 3. SDS-PAGE and immunoblot detection of denatured or semi-native BabA

SDS-PAGE was performed with 7.5% or 10% Mini-PROTEAN TGX Gels (Bio-Rad Laboratories, Hercules, CA, USA). Bacterial extracts оr the BabA protein solution were applied in Laemmli buffer under non-reducing, semi-native conditions and mild heating at 37°C for 30 min or under denaturing conditions (5% β-mercaptoethanol or 10 mM DTT and 10 min boiling) prior to loading onto the gels. For immunoblots, proteins were transferred to a polyvinylidene difluoride membrane (Bio-Rad Laboratories, USA) and blocked with 5% skim milk. Detection of native/recombinant BabA with human sera or ABbA-scFv on immunoblots was as described above, with the human sera being diluted 1:1000 and incubated overnight at 4°C. Secondary antibody HRP-conjugated goat anti-rabbit antibody (DAKO, Glostrup, Denmark) was diluted 1:1000 or ECL sheep anti-human HRP-conjugated (GE-Healthcare, USA) was diluted 1:5000 and incubated with the membrane for 1 h at room temperature. Bound ABbA-scFv was detected by incubation of the membrane with biotinylated murine anti-myc mAb 9E10 (Abcam, Cambridge, UK) followed by a streptavidin-HRP conjugate. Signals were developed with ECL chemiluminescence (SuperSignal West Pico, Thermo Scientific/Pierce, IL, Rockford, USA). For visualization of the signal, the Bio-Rad ChemiDoc^TM^ Touch Imaging System (Bio-Rad Laboratories, Inc) was used.

### 4. Purification of human IgG

A volume of 200 µl of human serum (total protein concentration 5 mg/mL) was diluted with 2.8 mL of binding buffer (20 mM sodium phosphate, pH 7.0). The sample was applied by a syringe onto a 1 mL HiTrap Protein G High Performance column (Cytiva, USA). The flow-through fraction with a total protein concentration of 4 mg/mL was saved for RIA analysis. The column was washed with 10 column volumes of binding buffer, and IgG was eluted as a 1 mL fraction with 10 mM glycine (pH 2.7) into tubes containing 100 µl 1M Tris (pH 9.0) and ≈2.5 mg/mL of protein. The protein concentration in the plasma and the flow through was determined by Pierce BCA Protein Assay Kit (Thermo Scientific, Rockford, IL, USA) according to the manufacturer’s instructions. The purified IgG fraction concentrations were determined by absorbance at 280 nm with a Nanodrop ND-1000 Spectrophotometer (NanoDrop Technologies, In., USA). Human IgG was identified by immunoblotting with sheep anti-human HRP-conjugated antibody (GE Healthcare, USA) diluted 1:5000. The human IgG was detected as the 55 kDa heavy chain (HC) and the 30 kDa light chain (LC).

### 5. Isolation, purification, and expression of ABbA-IgG (Figure 4)

#### 5.1. cDNA synthesis and PCR of human variable regions

Peripheral blood mononuclear cells from 10 mL of patient blood were isolated on a Ficoll gradient, and total RNA was extracted using standard protocols (RNeasy-Mini, Qiagen, Hilden, Germany). First-strand cDNA was synthesized with an oligo-d(T) primer (Amersham Biosciences, Buckingham, UK), and human variable immunoglobulin genes were amplified via PCR in 50 µL reactions containing 1 µL cDNA, 200 µM dNTPs, 5 µL 10x reaction buffer, 1 U polymerase (BD-Advantage2, BD Biosciences Clontech, Palo Alto, CA, USA), and the appropriate family-based sense and antisense primers (500 nM) for 36 cycles (15 s denaturation at 94°C, 30 s annealing at 65°C, and 30 s extension at 72°C). The sense and antisense primers were described elsewhere ^46-48^. For VL-chain amplification, the sense primers were extended at the 5’ end by the sequence TAC AGGATCC<u>ACGCGT</u>A in order to introduce an MluI restriction site, and the antisense primer was extended with TGACAAGCTT<u>GCGGCCGCG</u> for introduction of a Not I site. For VH-chain amplification, the sense primers were extended with GAATAGG<u>CCATGG</u>CG (<u>Nco I</u> site) and the antisense primer with CAGTC<u>AAGCTT</u> (<u>Hind</u> <u>III</u> site). The antisense primers annealed at the 5’ ends of the CH1 and constant Kappa regions, respectively. All amplifications were performed independently for each of the family-specific sense primers. The PCR products were pooled, gel-extracted (QIAquick Gel Extraction Kit, Qiagen), and digested with Mlu I and Not I (New England Biolabs, USA) for cloning of VL chains and with Nco I and Hind III for cloning of VH chains. After digestion, the fragments were gel purified again and stored at –20°C.

#### 5.2. ScFv library construction

The phagemid vector pSEX81-phOx ^24^ was digested with Mlu I and Not I and separated from the phOx scFv cassette by extraction in a 0.7% agarose gel. One hundred nanograms of the digested vector were ligated with 10 ng of the purified VL chains in a final volume of 40 µL with 1 U ligase (Roche, Mannheim, Germany) at 16°C overnight. Plasmid DNA was ethanol precipitated and electroporated into the *E. coli* strain XL1-Blue (Stratagene, La Jolla, CA), and the bacteria were grown for 1 h in 1 mL SOC media (LB plus 0.1 M glucose) for recovery. The bacteria were subsequently plated on SOB_GAT_ plates (0.1 M glucose, 100 µg/mL ampicillin, 12.5 µg/mL tetracycline) and incubated overnight at 37°C. Colonies were scraped off, and vector DNA containing the VL chain repertoire was isolated using anion exchange chromatography columns (Macherey & Nagel, Germany).

For cloning of the VH chains, vector DNA was digested with Hind III and Nco I (New England Biolabs), ligated with the appropriately digested VH-encoding genes, transformed into XL1-Blue cells, and grown as described above. Independent colonies were scraped off and stored in 25% glycerol at –80°C as the final scFv library.

#### 5.3. Selection of BabA-binding scFvs by phage display (fig S4A)

Phage scFvs were retrieved from the libraries essentially as described ^49^ using helper phage M13KO7 (New England Biolabs, USA) for packaging. Panning was performed in Maxisorb Immunotubes (Nunc, Wiebaden, Germany) coated overnight with 5 µg of purified BabA (according to ^14^) at 4°C and blocked with PBS containing 2% (w/v) low-fat dried skimmed milk (2% M-PBS). Tubes coated with BSA were used as negative controls. For selection, phage-scFvs (10^12^ CFUs) were blocked by the addition of an equal volume of PBS containing 4% milk (w/v) to the tubes and incubation with constant rolling for 2 h at room temperature. The solution was subsequently discarded, and the tubes were washed 10 times with PBS in the first panning round. With subsequent panning rounds, washing stringency was increased by vortexing 10 times with 0.1% Tween 20 in PBS. Bound phages were eluted by addition of 1 mL 0.1 M triethylamine (pH 11.5) for 5 min with gentle agitation and neutralization with 0.5 mL 1 M Tris-HCl (pH 7.4). The neutralized mixture was used to infect 20 mL of exponential-phase *E. coli* XL1-Blue grown in 2YT media (12.5 µg/mL tetracycline) at 37°C. After incubation for 15 min at 37°C without shaking, the bacteria were shaken for 45 min, plated on SOB_GAT_ plates, and incubated overnight at 37°C. The bacteria were harvested and stored in aliquots at –80°C, representing a phagemid-packaged scFv-library. For the subsequent panning round, phages were produced by inoculation of 10 mL LB_GAT_ media with an aliquot of the library in order to reach an OD_600nm_ = 0.4. The titers of eluted phage containing either helper phage or the phagemid genomes were determined by CFU titration on LB plus kanamycin (70 µg/mL) or LB plus ampicillin (100 µg/mL) plates, respectively, essentially as described^50^. The enrichment of specific binders during the selection procedure was determined by dividing the number of phage that were eluted from BabA-coated immunotubes by the number of phage that were eluted from BSA-coated immunotubes (the enrichment factor was 55 in the second and 2381 in the third panning rounds).

#### 5.4. ELISA screening with scFv-gIII fusion proteins

BabA-specific scFvs were screened by taking advantage of the pIII protein encoded in the phagemid vector essentially as described ^51^. Briefly, production of scFv-gIII fusion proteins in logarithmically growing bacteria was induced with 100 µM isopropyl β-D-1-thiogalactopyranoside (IPTG) for 16 h at 30°C. Bacteria were centrifuged and the pellet was incubated in Spheroblast solution (50 mM Tris-HCl, pH 8.0, with 20% sucrose and 1 mM EDTA) for 20 min on ice, followed by centrifugation at 30,000 x *g* for 45 min at 4°C. The supernatant, representing the periplasmic extract, was diluted with an equal volume of 4% M-PBS and used for ELISA. Wells (Nunc Microtitre plates, Wiesbaden, Germany) were coated with 200 ng of BabA overnight at 4°C in 50 mM Na_2_CO_3_-NaHCO_3_ (pH 9.6). After blocking with 2% M-PBS, the periplasmic extract was added and incubated for 4 h at room temperature. Antigen-bound scFv-pIII fusion protein was detected by incubation with a mouse mAb specific for pIII (MobiTec, Göttingen, Germany ^24^ for 1 h at room temperature followed by an HRP-conjugated rabbit anti-mouse antibody (DAKO, Glostrup, Denmark) for 1 h at room temperature. Colorization was performed with 0.42 mM 3,3’,5,5’-tetramethylbenzidine (Merck-Germany) in substrate buffer (83 mM sodium acetate, 17 mM citric acid (pH 4.9), and 0.004% H_2_O_2_). After all incubation steps, the ELISA wells were washed three times with blocking buffer. As a negative binding control, a 2-phenyloxazolone (anti-phOx) scFv was expressed in the same phagemid vector and analyzed using ELISA-coated, phOx-conjugated BSA.

#### 5.5. Subcloning into the prokaryotic expression vector pOPE101 (Figure S4)

The entire scFv expression cassette from the phagemid vector pSEX81 was subcloned into the prokaryotic expression vector pOPE101 (Genbank #Y14585) via the Nco I and Not I sites. The C-terminal myc and (His)_6_ tags allowed detection and IMAC purification, respectively. Purification from the periplasmic space was performed as described ^24^.

#### 5.6. Production of a complete human antibody (Figures 4 and S4)

Variable regions were cloned into the insect cell expression vector pMThIgG1-V carrying the constant regions of the human IgG1 heavy chain and human kappa chain ^24^. PCR was performed using the VL chain primer pair *VL5’SfiI* (TTA CTC GCC TGG CCG TCG TGG CCT TTG TTG GCC TCT CGC TGG GCG ACA TCC AGA TGA CCC AGT C) and *VL3’BsiWI* (AGC GTA CGT ACG TTT GAT TTC CAC CTT GGT CC) and the VH chain primer pair *VH5’SnaBI* (GAT GTC TAC GTA GGC CTC TCG CTG GGC CAG GTG CAG CTG GTC CAG TC) and *VH3’ApaI* (ACC GAT GGG CCC TTG GTG GAG GCG GAG GAG ACG GCG ACC AGG G). PCR was performed with 100 ng of the phagemid vector as the template, 25 pmol each of the VL and VH primer pair, 2 µM MgCl_2_, 0.2 mM dNTPs, and 10 U of Taq polymerase (Promega). After an initial denaturation at 94°C for 2 min, 32 cycles were run with 15 s at 94°C, 30 s at 62°C, and 30 s at 72°C, with a final elongation step of 5 min at 72°C. PCR products were purified with the QIAquick PCR purification kit (Qiagen) and digested with the appropriate restriction enzymes prior to cloning. A stable antibody-secreting S2 Schneider cell line (Invitrogen) was established as previously described ^45^, and antibodies in the media were purified and enriched using protein G columns (GE Healthcare, Pittsburgh, PA). The purity and functionality of the purified antibodies were analyzed by Coomassie staining and ELISA, respectively.

#### 5.7. Immunoelectron microscopy and detection of BabA by ABbA (Figure S4G)

Bacterial cells from *H. pylori* 17875/Leb and the 17875*babA1A2* mutant were grown and adjusted in PBS to an OD_600_ of 1. An aliquot of each strain was resuspended in 2% BSA in 0.1 M sodium cacodylate buffer for 15 min, centrifuged, and resuspended with ABbA-IgG or an irrelevant anti-HCV antibody of the same human IgG1-isotype diluted 1:1 in 0.1 M sodium cacodylate plus 0.1% BSA and incubated for 60 min. After incubation, the bacteria were washed twice in 0.1 M sodium cacodylate plus 0.1% BSA and resuspended in the same buffer containing protein A conjugated to 10-nm gold particles (GE Healthcare) and incubated for 45 min. The incubation was terminated by adding glutaraldehyde to a final concentration of 1%, and the samples were fixed overnight at 4°C. Bacteria were subsequently pelleted and small aliquots were resuspended in distilled water. Small drops (3 µL) were placed on formvar-coated grids and allowed to attach for 5 min. Excess water was removed with filter paper and then the grids were air-dried for 5 min and then examined on a Tecnai 10 electron microscope (Fei, Eindhoven, Netherlands) at 100 kV, and the images were recorded on photographic film.

#### 5.8. Construction of a random BabA shotgun gene-fragment library (**Figure 5A**)

The phagemid vector pSEX81(phOx) was prepared for blunt end ligation by digestion with PvuII and EcoRV followed by agarose gel purification. The purified vector was treated with shrimp alkaline phosphatase (NEB, USA) to prevent self-ligation. The full-length *babA* gene from *H. pylori* strain 17875/Leb was excised from a midi-plasmid preparation by restriction enzyme digestion and purified on an 0.8% agarose gel. The purified *babA* gene was sonicated with a Covaris S sonicator to obtain DNA fragments with a size of 400, 800, or 1500 bp. DNA fragments were polished by treatment with T4 DNA polymerase, and 5’-ends were subsequently phosphorylated by T4 polynucleotide kinase (Next End Repair Module, NEB, USA). Fragments were purified on a Qiagen column (PCR purification kit) and ligated overnight with 200 ng of the prepared phagemid pSEX81 plasmid. Electroporation and library preparation were performed as described in ^49–51^, and aliquots were stored at –80°C.

#### 5.9. Selection of ABbA-IgG-binding BabA fragments by phage display (**Figure 5A**)

Phage display selection was performed essentially as described in ^49–51^, except that Hyperphage M13KO7ϕλpIII was used for phage packaging (Progen-Heidelberg, Germany) and immunotubes were coated with 10 μg ABbA-IgG or HSA-Leb.

### 6. The BabA protein co-crystallized with ABbA-IgG (Fab) (Figure 5C **and** Figure S5E, Table S5)

To generate the BabA-ABbA complex, purified BabA^AD^ (residues 25 to 460 of mature BabA) and ABbA-Fab were mixed at a 1:1 stoichiometric ratio. Crystal formation of the BabA-ABbA complex was enhanced by use of nanobody Nb-ER19 as described in ^13, 52^. Nb-ER19 is a camelid VHH single domain antibody that binds the 3+4 alpha domain of BabA^AD^, stabilizes BabA^AD^, and facilitates its crystallization. Binding of Nb-ER19 is distant from the CBD and does not interfere with the binding of glycan receptors or of ABbA to BabA. BabA^AD^ and Nb-ER19 were produced and purified as described previously ^13^. The ABbA-Fab was generated using the Pierce Fab Preparation Kit according to the manufactureŕs instructions (ThermoScientific, cat#44985). The BabA-ABbA-Nb-ER19 complex (40 mg/mL in 20 mM Tris-HCl (pH 8.0) and 10 mM NaCl) was crystallized in 15% PEG6000, 0.1 M NaAc (pH 5.5), and 0.2 M CaCl_2_ by sitting drop vapor diffusion. For data collection, crystals were cryo-protected by brief transfer into the crystallization buffer supplemented with 15% glycerol, after which crystals were loop mounted and flash cooled in liquid nitrogen. X-ray diffraction data were collected from a single crystal at beamline ID29 of the ESRF (date 31/08/2014) at a wavelength of 1.072 Å to a final resolution of 2.7 Å. Diffraction data were integrated and scaled using XDS and XSCALE ^53^, and the structure was solved by molecular replacement using Phaser ^54^ and BabA^AD^-Nb-ER19 (PDB: 5F7W) and an unrelated FAB (PDB: 1S5H) as search models. The final model was obtained after iterative cycles of manual building using Coot ^55^ and maximum likelihood refinement against the X-ray data using Phenix Refine ^56^ resulting in a model that contained two copies of the BabA^AD^-ABbA-Nb-ER19 complex per asymmetric unit, with an R and free R-factor of 22.9% and 28.3%, respectively, and 98.9% of the residues were in allowed regions of the Ramachandran plot. The BabA crystal structure was displayed with DNASTAR/Lasergene Protean 3D Software.

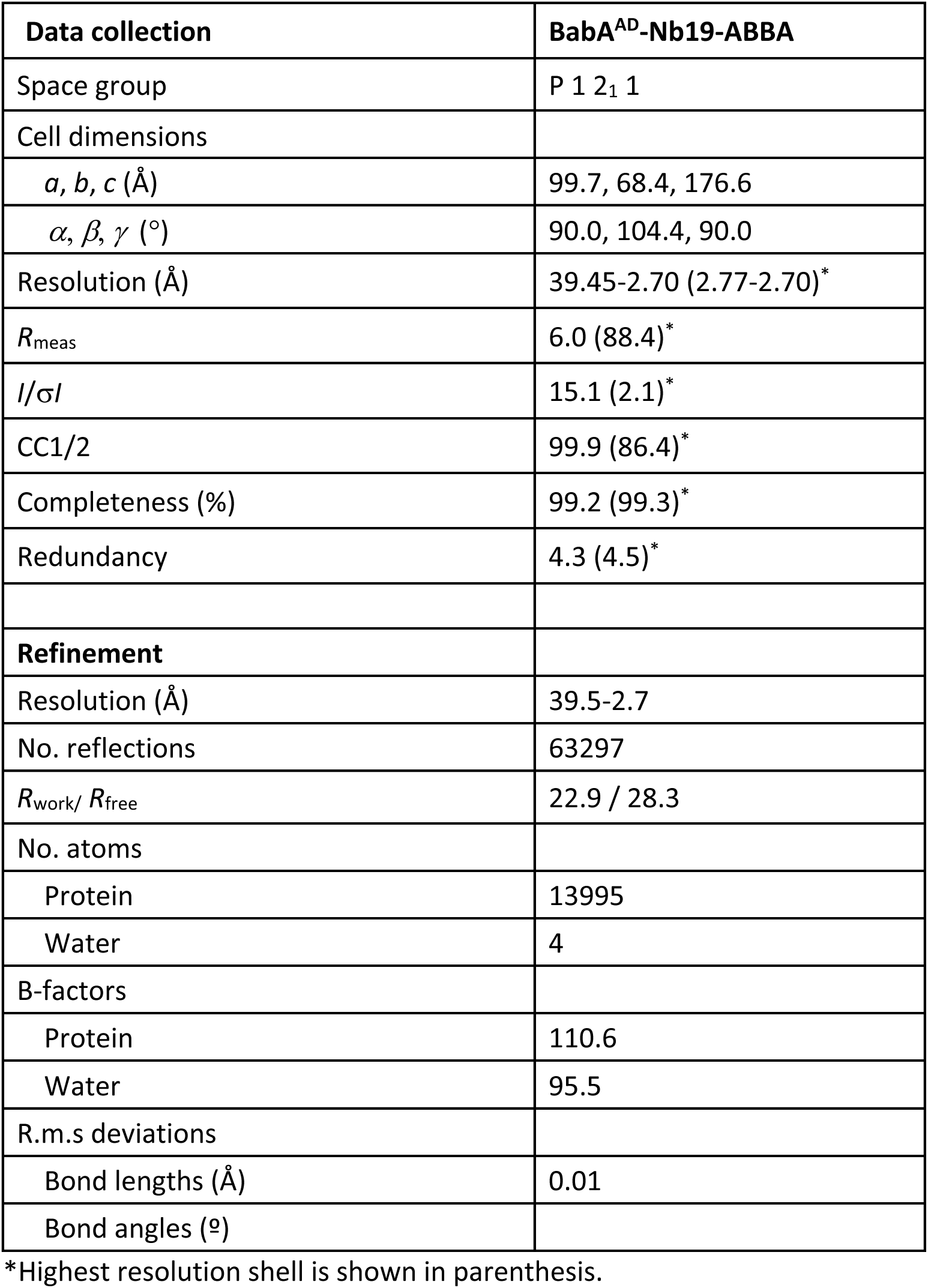

### 7. Shuttle vector design, genetic complementation by conjugation, and recombinant expression of the S234A and D233A-S234A-S235A (DSS mutants) BabA in *H. pylori* (Figure S5G)

The *H. pylori* shuttle vector pIB6 is based on the pHel3 plasmid ^57^ that has been further modified to contain the *alpA* promoter upstream of a multiple cloning site where the various *babA* alleles were introduced ^14, 58^. The *babA* gene from 17875/Leb was cloned into the shuttle vector pIB6 and transformed into the *H. pylori* P1Δ*babA* strain as described by ^14^. The pIB6-*babA*17875/Leb was used for the mutagenesis procedure where the S234A mutant and the DSS mutants were constructed by PCR-based site-directed mutagenesis using Genscript (Piscataway, NJ, USA). Fragments were re-cloned in pIB6 using NdeI and NotI restriction sites to make up the shuttle vector plasmids (SV) SVS234A and SVDSS/AAA that were transformed into the *dapA*-negative *E. coli* strain β2150. This “donor” strain is strictly dependent on exogenously supplied diaminopimelic acid. Hence, the removal of diaminopimelic acid provides an efficient counter-selection against this donor. All “donor” strains with the various *babA* shuttle vector plasmids were conjugated into the *H. pylori* strain P1Δ*babA*. Plasmids from transformed *H. pylori* clones were re-isolated, and the full *babA* sequence from each mutant type was sequenced at MWG Operon, Germany. The tested clones showed the correct sequence.

### 8. Amplification and analysis of BabA sequences

Bacterial cells of strains listed in **Table S7** were grown, and colonies were removed from the plates, washed, and re-suspended in 1 mL of PBS to OD_600nm_ = 1.0. This suspension was used to isolate genomic DNA according to the manufacturer’s instructions (DNeasy Blood and Tissue Kit, Qiagen). BabA fragments covering nucleotide 1 to approximately nucleotide 1200 were amplified by PCR with 4 µL of genomic DNA as the template and a combination of either one of the two forward primers babA2-271 ^11^ (5’-ATC CAA AAA GGA GAA AAA ACA TGA AA-3’) or babA2-Leader (5’-GCT TTT AGT TTC CAC TTT GAG-3’) and either one of the two reverse primers J11R (5’-TGT GTG CCA CTA GTG CCA GC-3’) or A26R (5’-TTG CTC CAC ATA GGC GCA C-3’). The PCR fragments were ligated into pGEM-T (Promega, Madison, CA, USA) and sequenced with T7 and SP6 promoter-specific primers. Nucleotide sequences were determined by the dideoxy chain-termination method of Sanger using the Big Dye Terminator Cycle Sequencing Kit (Applied Biosystems, Foster City, CA, USA) or using the services of Eurofins (Ebersberg, Germany). Assembly and sequence analysis was performed with Vector NTI version 10 (Invitrogen). Sequences that originated from Aspholm *et al.* ^11^ are indicated by stars. Sequences that originated from Bugaytsova et al. ^14^ are indicated by triangles. Mexican Mc1215, Mc1207, and Mc1201 were sequenced by the J. Torres lab (co-author). Sequence analysis including alignments using the ClustalW algorithm was performed using DNASTAR/Lasergene MegAlign Pro Software.

### 9. Amino acid positions in the BabA CBD that modulate ABbA binding affinity (Figure 6D and Table S7)

The amino acid residues were derived from BabA alignments of 123 strains from world-wide populations. The alignments were based on high-affinity binding strains that bind ABbA >20% *vs.* low-affinity binding strain that bind ABbA <2%. The amino acid configuration and composition at each position is expressed in terms of the relative size of the amino acid 1-letter code.

### 10. Genome sequencing of strain CCUG17875

DNA was extracted from bacterial biomass using a modified Marmur procedure ^59^. The short read library was prepared using the Illumina DNA PCR-Free protocol aiming for insert sizes of 550 bp and was subsequently sequenced using the Illumina MiSeq v3 chemistry generating 2*300 bp reads. Libraries for long read sequencing were barcoded using the SQK-RBK004 Rapid Barcoding Kit and sequenced on the Oxford Nanopore MinION platform, flow cell version FLO-MIN106. Raw Fast5 files were base called and demultiplexed using the Guppy software v. 4.2.2. Hybrid assemblies of Illumina and ONT reads were performed using the Unicycler software v0.4.8 ^60^. The de novo assembly produced a circular, complete chromosome of 1,635,834 bp and a plasmid of 8,108 bp. Genome sequencing data for *H. pylori* CCUG 17875 and its plasmid are available from GenBank under BioProject PRJNA793037 with accession number CP090367 and CP0903678, respectively.

### 11. Construction of the phylogenetic tree (Figure 2B)

The phylogenetic distance was calculated based on kmer sharing using mash v. 2.0. (REF: Mash: fast genome and metagenome distance estimation using MinHash ^61^). The tree was generated using RapidNJ v. 2.3.2. Genomic sequences used as worldwide references were:

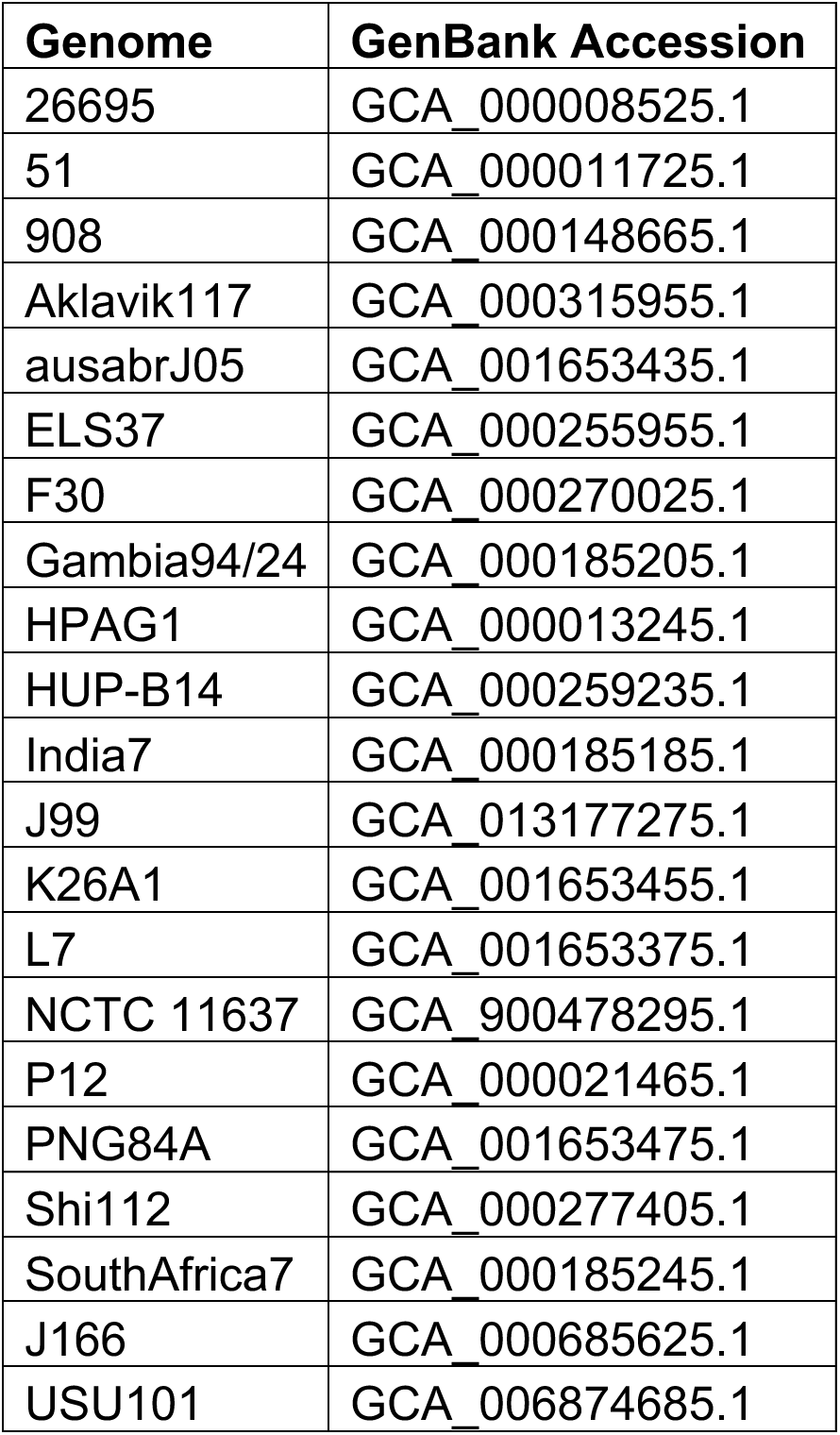

### 12. Acid sensitivity in *H. pylori* BabA-mediated Leb binding and Leb inhibition (Figure 6E)

Acid sensitivity of *H. pylori* binding was performed according to ^14^. Acid sensitivity of scFv and ABbA binding to the BabA of *H. pylori* 17875/Leb was assessed by incubation of the radiolabeled scFvs or ABbA-IgG in nine buffers with pH ranging from 2.0 to 6.0 with 0.5 pH unit increments (as described in ^14^). The assay was performed by mixing 890 µl of pure pH buffer, 10 µl of radiolabeled scFv or ABbA-IgG in the corresponding pH buffer, and 100 µl of bacterial suspensions (OD_600_ = 1.0) in blocking-buffer (pH 7.4). The obtained suspension was incubated on a rocking table at room temperature for 1 h, and the cells were pelleted by centrifugation. CPMs in the pellet and supernatant were measured using the 2470 Wizard^2^ Gamma Counter (PerkinElmer, Waltham, MA, USA), and binding at each pH was determined as described in **2.3. Analysis of BabA binding properties by RIA.**

### 13. QUANTIFICATION AND STATISTICAL ANALYSIS

Data were analyzed using Graphpad Prism 9.0 (Graph Pad software, La Jolla, CA, USA) or R, version 4.0.2 (**R Core Team, 2020**). Depending on the experimental design and type of variables under investigation, different methods were utilized. Statistical significance regarding group comparisons was investigated via either unpaired or paired Student’s t-test or the Wilcoxon signed rank test, while associations between variables were tested using Pearson and/or Spearman correlation or Odds Ratios. Further classification performance was assessed using Area Under the Curve (AUC) in a Receiver Operating Characteristic (ROC) curve and by permutation tests. Details on the tests used can be found in the respective figure legends. P-values < 0.05 were considered significant (*p < 0.05; **p < 0.01; ***p < 0.001), and p > 0.05 was non-significant (NS). Tests were two-tailed unless otherwise stated.

#### Reference for R; R Core Team (2020)

R: A language and environment for statistical computing. R Foundation for Statistical Computing, Vienna, Austria. URL https://www.R-project.org/.

## Supplemental figures

**Figure S1.**
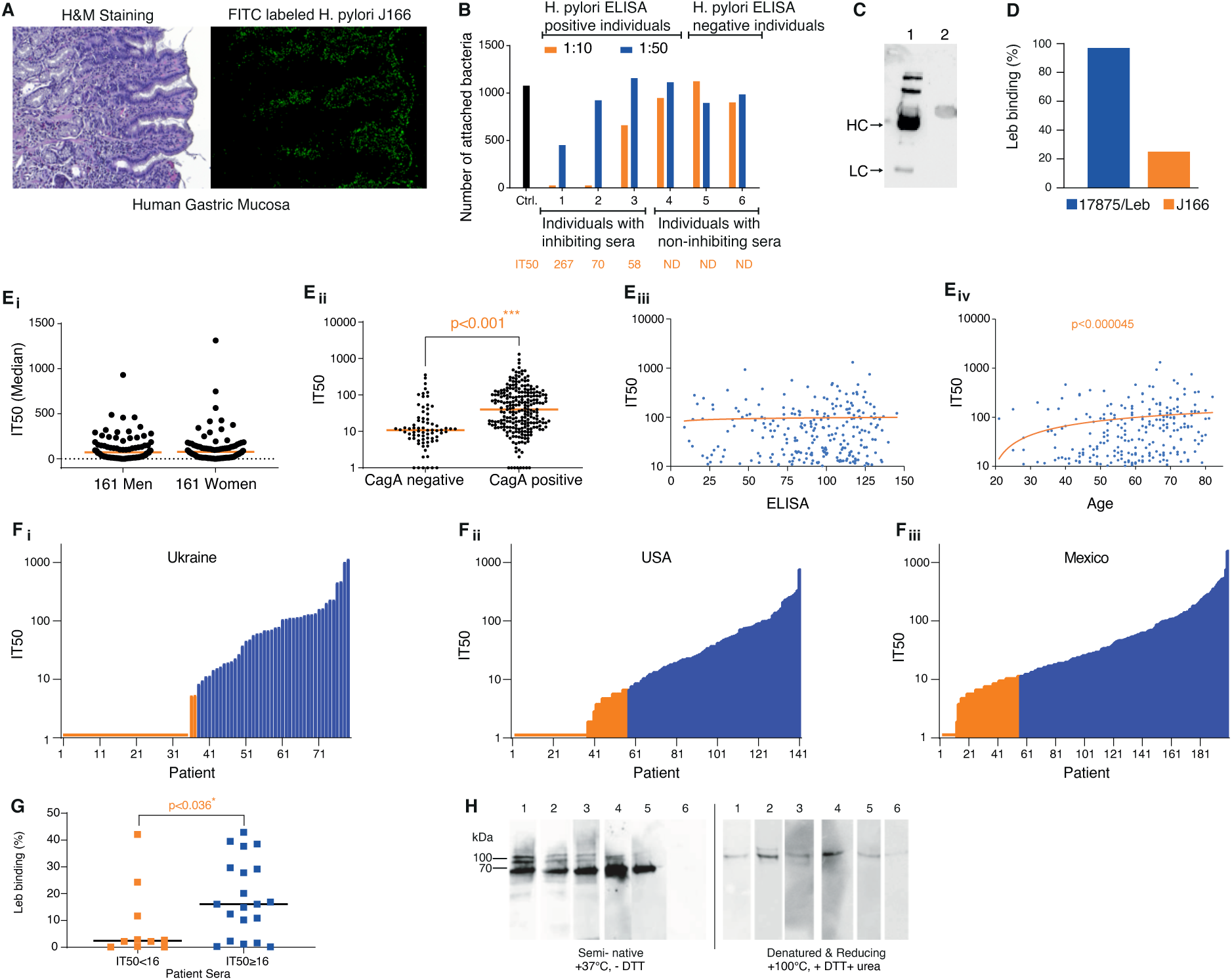
High global prevalence of serum inhibition of Leb binding. (**A***i*) H&E-stained adjacent sections of human gastric mucosa from Figure 1A, (A*ii*) and *in vitro* attachment by *H. pylori* J166 bacterial cells labeled with FITC-green fluorescence. (B) Quantification by ImageJ of attached J166 bacteria treated with 1:10 and 1:50 dilutions of individual serum samples from Figure 1A. Individuals 1-4 were ELISA positive for *H. pylori* infection, whereas individuals 5 and 6 were ELISA negative. Individuals 1–3 demonstrated IT50s of 267, 70, and 58, in contrast to individuals 4–6 with non-detectable (ND) titers (Figure 1B). (C) Immunoblot of the SDS-PAGE-separated serum proteins from individual 1 (from Figure 1A) detected with anti-human Ab. The heavy chain (HC) and light chain (LC), i.e., the IgG components, were fully removed by Protein G affinity desorption (sera before (1) and after (2) IgG desorption). (D) *H. pylori* J166 with only 25% “Hot-Leb binding” is a low-affinity Leb-binding strain, compared to 17875/Leb, with >95% “Hot-Leb binding” which is a high-affinity Leb binding strain. High and low-affinity Leb-binding *H. pylori* strains have been identified (**Figure S2B**). The 25% Hot-Leb binding by strain J166 corresponds to ∼50-fold lower affinity according to Scatchard tests, which is a radio immuno-assay (RIA) for affinity testing based on binding of Leb under equilibrium conditions ^42^. (**E***i*) Among the 322 Kalixanda Swedish *H. pylori* carriers (95% of whom were gastric healthy), the 161 men with median age 59 years and 161 women with median age of 61 years did not exhibit any difference in median IT50 (**Table S1B**). (**E***ii*) The CagA status by ELISA among 317 out of 322 Kalixanda individuals (5 individuals were not tested for CagA) exhibited strong correlation with IT50, with a median IT50 = 11 for CagA-negative individuals and a median IT50 = 40 for CagA-positive individuals. *** p < 0.001. (**E***iii*) The general immune response against *H. pylori* (ELISA) and IT50 among the 322 Kalixanda individuals showed no correlation. (**E***iv*) A rank correlation test showed that the IT50s increased with age among the 322 Kalixanda individuals, r = 0.28, p ≤ 0.000045. This was most similar to a regular correlation test, r = 0.18, p ≤ 0.001. (**F***i*) The IT50s of 79 ELISA-positive individuals from Sumy, Ukraine, tested with strain 17875/Leb with a median IT50 = 69 and a mean IT50 = 125, where 44 (56%) sera samples were positive for Leb inhibition by strain J166 (**Table S1C**). (**F***ii*) The IT50s of 141 ELISA-positive individuals from the US tested with strain 17875/Leb with a median IT50 = 39 and a mean IT50 = 79, where 83 sera samples (59%) were positive for Leb inhibition by strain J166 (**Table S1D**). (**F***iii*) The IT50s of 200 ELISA-positive individuals from Mexico City, Mexico, tested with strain 17875/Leb with a median IT50 = 50 and mean IT50 = 115, where 146 sera samples (72%) were positive for Leb inhibition by strain J166 (**Table S1E**). (G) The 30 Mexican sera and corresponding strains (**Table S2B**) were tested for correlation between IT50s and the Leb binding property. Low IT50 (<16) sera are in orange and high IT50 sera (≥16) are in blue with medians indicated (Wilcoxon p = 0.036). (H) Immunoblot detection by serum samples from (1) the Karolinska University Hospital (Sweden) (Figure 1I), (2) Ukraine, (3) the US, (4) Mexico, (5) the Kalixanda series (Sweden), and (6) the ELISA-negative individual 5 from Figure 1A. BabA protein was purified from strain 17875/Leb and separated by semi-native SDS-PAGE (samples were kept at 37°C for 10 min without reducing agent) or under full denaturing and reducing conditions (samples heated to 100°C in reducing SDS sample buffer with 8M urea and 10 mM DTT). BabA binding by sera antibodies was detected using HRP-conjugated anti-human antibody.

**Figure S2.**
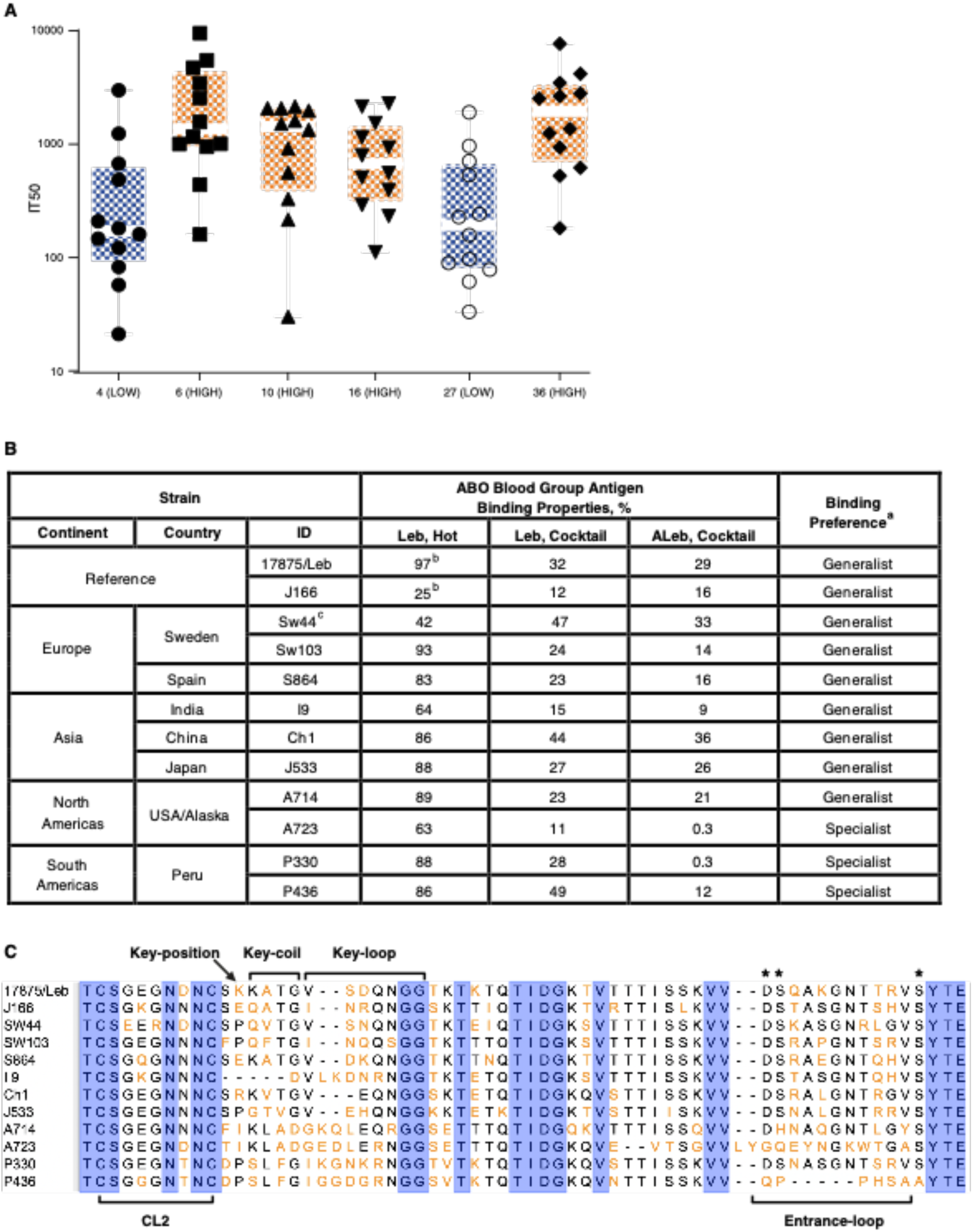
The broadly blocking serum Abs. (A) A box-plot presentation of the IT50 titers for the 12 strains in Figure 2A showing the relatively higher variation in IT50 titers for the two lower-inhibitory sera (LOW in blue, individuals 4 and 27) compared to the four sera with Higher inhibitory titers (HIGH in orange, individuals 6, 10, 16, and 36). In the log-scale presentation, the IT50 titers demonstrate increased spans between the quartiles for the LOW serum samples (Ansari-Bradley p = 0.07, i.e., close to significant and significant by removal of one outlier and between the “highest” from the LOW sera and the “lowest” from the HIGH sera (Ansari-Bradley p = 0.029). The results suggest that the LOW sera displayed a higher level of discrimination for different strains compared to the HIGH sera that efficiently blocked the Leb binding of the majority of *H. pylori* strains. (B) Binding properties and binding preferences of *H. pylori* strains tested for inhibition by human sera (Figure 2A) and by ABbA (Figure 4A). ^a^The Indigenous South American specialist isolates bind to Leb much better than to ALeb (from 2.5-fold to ∼100-fold), i.e., there is a Specialist preference for binding to blood group O antigen. In comparison, the common Generalist binding preference is defined as the Leb/ALeb-ratio interval from 1:1 to 1:2.5 ^11^. ^b^The Hot Leb binding is from **Figure S1D**. ^c^The ABO/Leb binding preference of *H. pylori* Sw44 is described in ^11^. (C) Alignment of the central part of the BabA CBD of the *H. pylori* isolates from Figure 2A. The strains were chosen with respect to differences in Leb-binding affinity, binding preference (ABO-Generalists *vs.* O-Specialists), and BabA phylogeny, illustrated in (B) ^11, 14^. The stars indicate the locations of the conserved Leb-binding DSS triad residues ^13^. However, exceptions do exist, and the DSS residues in the BabA Entrance loop that are critical for binding to ABO/Leb can be modified by a change of charge or hydrophobic substitutions or by deletions, e.g., in the Specialist strains that only bind to the blood group O antigen, which is a common adaptation in binding preference among Indigenous American/Latin American strains such as A723 and P436 ^11, 13^.

**Figure S3.**
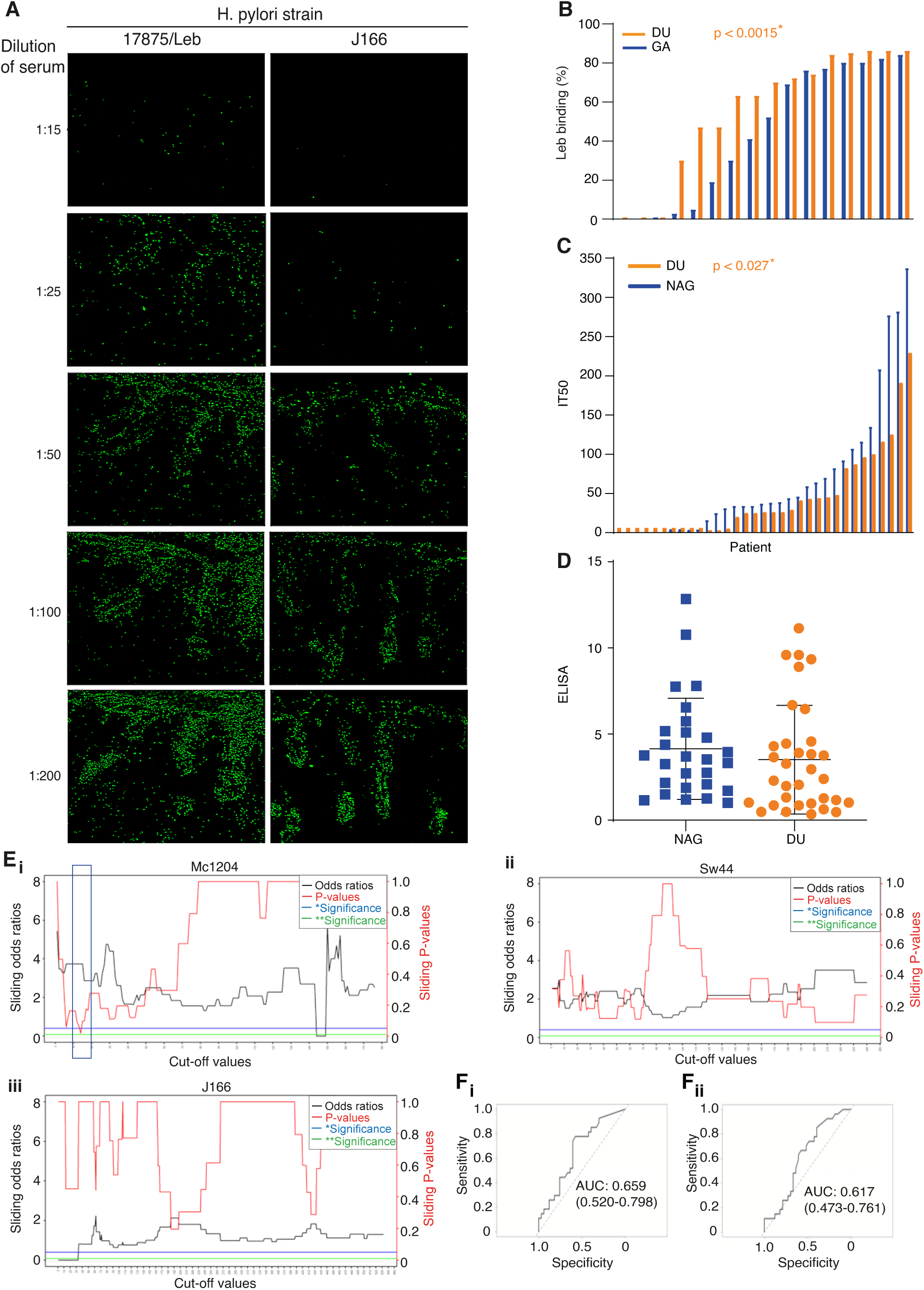
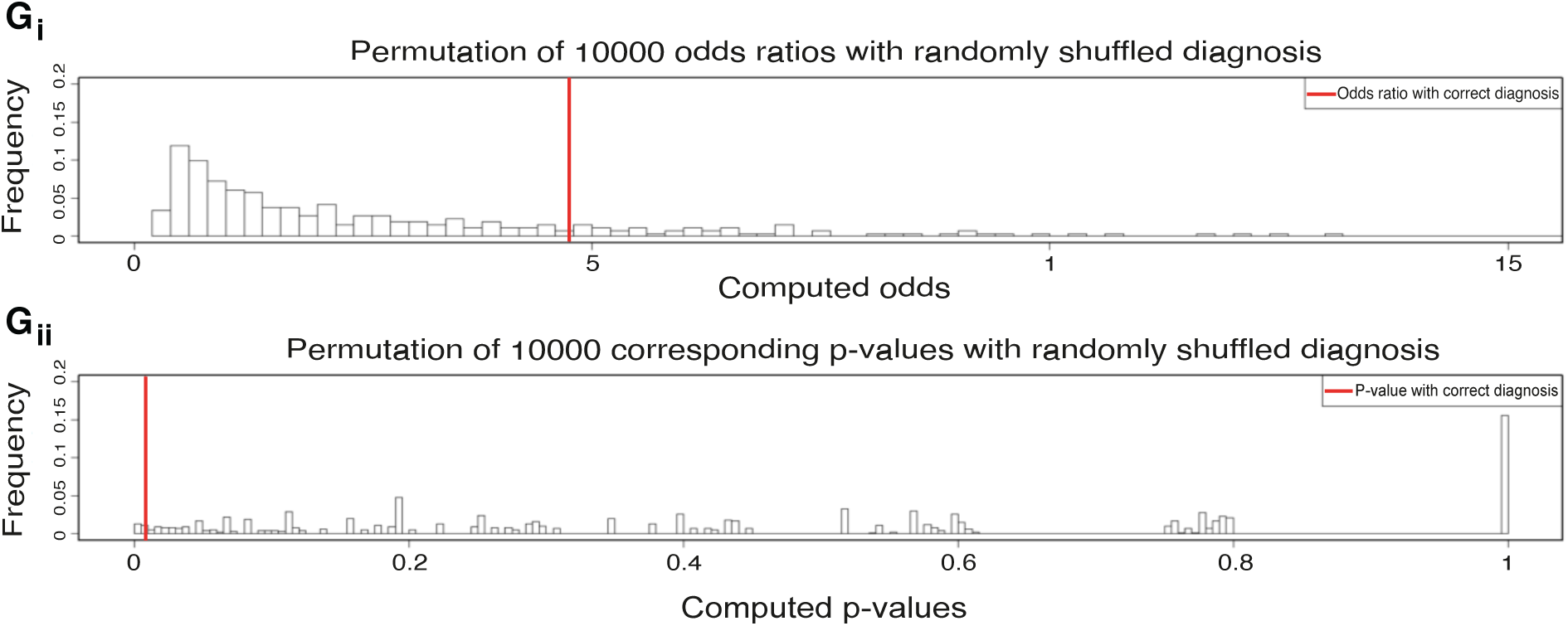
Broadly blocking serum Abs protect against overt gastric disease. (A) Strains 17875/Leb and J166 were tested for blocking of *in vitro* binding to human gastric mucosa (Figure 2B) by a dilution series of sera from individual 1 (from Figures 1A and 1B). Attached bacterial cells were quantified by ImageJ. (B) A series of 32 Mexican isolates from patients with GA or DU (16 isolates for both) were tested for Leb binding strength with 1 ng of ^125^I-labeled Leb-conjugate, i.e., a limiting concentration, defined as “Hot-Leb”. The DU isolates demonstrated generally higher binding strength compared to GA isolates (Wilcoxon signed-rank test, p = 0.0015) (**Table S2B**). (C) The 79 serum samples from NAG and DU patients demonstrated generally higher IT50 for patients with NAG (p < 0.027) (**Table S3**). (D) The 79 Mexican serum samples from patients with NAG or DU did not show significant differences in *H. pylori* ELISA, p < 0.4, i.e., the two patient groups exhibited similar general immune responses against chronic *H. pylori* infection (**Table S3**). (**E***i*) Sliding window for strain Mc1204; critical IT50 = 15 and p < 0.0189* (indicated by the box) (**Table S3**). (**E***ii-iii*) Sliding windows for strains Sw44 (ii) and J166 (*iii*), i.e., strains that are less discriminative for high *vs*. low-inhibition titers and thus do not provide the critical IT50 values with significant ORs (**Table S3**). Our approach using high *vs.* low-affinity binding strains is conceptually similar to tests with different mAbs for discrimination between sensitivity and specificity in epitope recognition. The high-affinity strains 17875/Leb (Figure 3C) and Mc2014 (**Figure S3E***i*) discriminate between high *vs.* low IT50 in contrast to the low-affinity binding strains J166 and Sw44 (**Figure S2B**), which do not produce significant ORs (***ii*** and ***iii***). However, the low-affinity binding strain J166 identified all IT50-positive serum samples with low IT50 *vs.* the true IT50-negative sera samples (**Table S3**). (**F***i*) The ROC diagram AUC 0.659 for strain 17875/Leb shows that the critical IT50 value that provides the highest significance is 29.5, i.e., the Youden index using the Area Under the Curve (AUC) where sensitivity and specificity are maximal. The described approach for finding optimal cut-offs using the OR as the discrimination measure can be seen as complementing a more common approach, namely that of using the Youden index while showing the overall classification performance across all possible cut-offs using an ROC curve and associated AUC. The optimal cut-offs identified using this approach are very similar, and we can thus view the “OR approach” as a way of illustrating the classification performance across the range of possible cut-offs. This complements the AUC, which gives a measure of overall classification performance. What we see using the OR approach is that the best classification performance (high ORs) is achieved within a narrow range of IT50 values around 30, i.e., where an optimal cut-off is located. This, together with the AUC value provided in the ROC, indicates that there is indeed useful information in the IT50 values. The ROC analyses were performed using the *pROC* package in R ^62^. (**F***ii*) The ROC diagram AUC was 0.617 for strain Mc1204, and the critical IT50 value was 15.5. (**G**) To verify the OR approach in an unbiased manner, we performed permutation tests where, in each iteration, we randomly permuted the outcome labels NAG/DU and then searched for an optimal Rf IT50 and its associated OR and its corresponding significance (p-value). We performed 10,000 iterations in each of these two permutation tests, and the F*i*-test showed that the number of ORs (i.e., classification performance) that exceeded OR 4.75 (red bar, from Figures S3B and 3C) were less than 2% of the permutation test iterations. In addition, a **G***ii*-test showed that the ORs with high corresponding significance (p-values) constituted non-random results (p = 0.027 and p = 0.0197, respectively).

**Figure S4.**
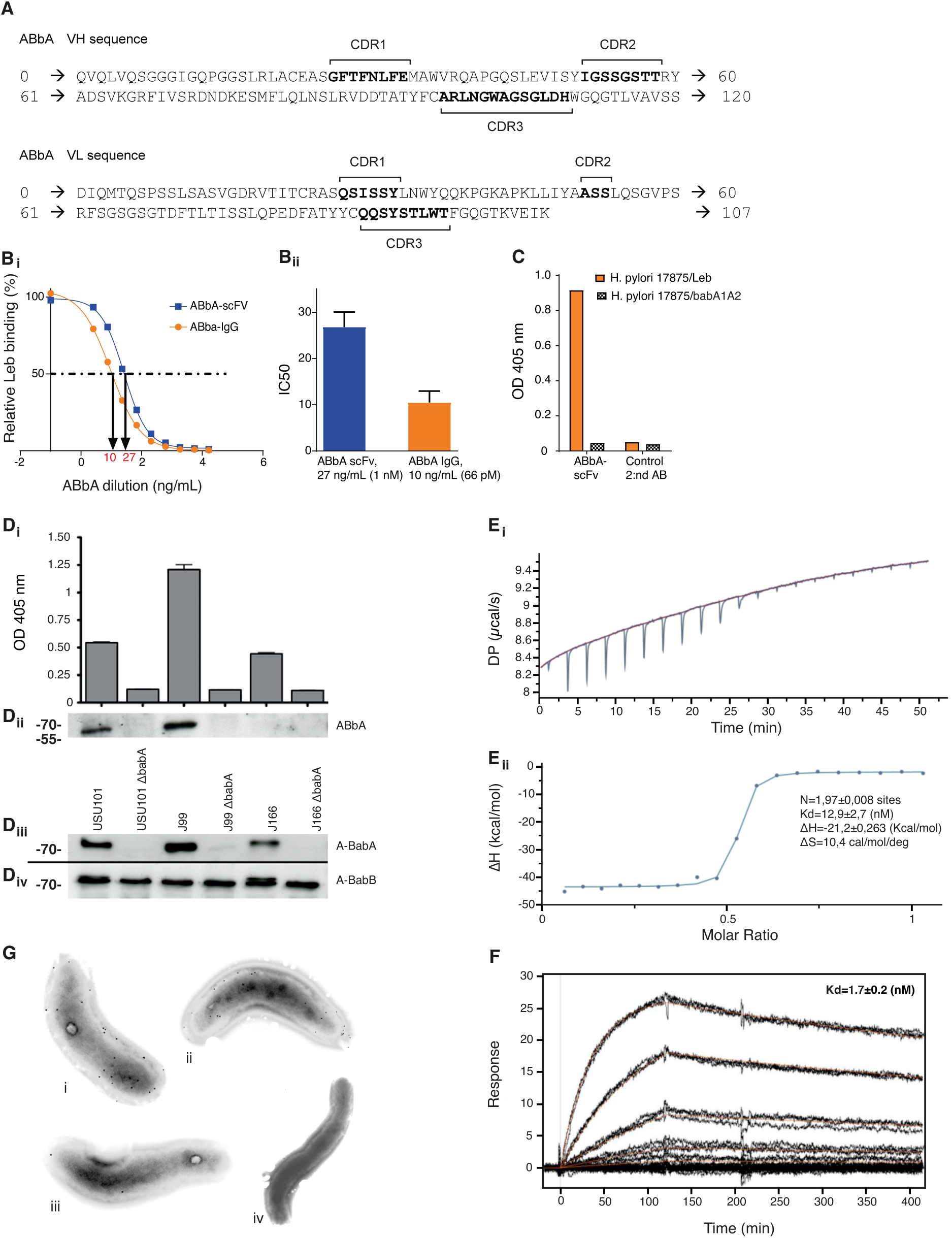
Cloning and identification of ABbA, the broadly blocking mAb. **(A)** Amino acid sequences of the ABbA VH and VL. The ABbA-VH region was generated by recombination of the germline gene IgHV3-48*3 with the D segment of IgHD2-15*01 and the J segment of IgHJ4*03. The high number of mutations in comparison to the germline gene indicates intense affinity maturation. The ABbA-VL chain was derived from the VL germline gene IgGKV1D-39*01 by recombination with the J segment of IgKJ1*01. Interestingly, the ABbA-VL chain was nearly identical to its most closely related germline gene. **(B*i* and B*ii*) Binding strength** of ABbA(1)-scFv (27 kDa) *vs.* ABbA-IgG (ABbA) (150 kDa) by strain 17875/Leb with a 16-fold difference in IC50 (the concentration of antibody that inhibits *H. pylori*-Leb binding by 50%) of 27 ng/mL (1 nM) and 10 ng/mL (66 pM), respectively. **(C) Specificity of ABbA binding.** ABbA(1)-scFv binds to *H. pylori* 17875/Leb but not to its isogenic 17875*babA1A2*-minus strain. ELISA wells coated with the two *H. pylori* strains were incubated with ABbA(1)-scFv. Binding was detected using an anti-myc mAb and HRP-conjugated anti-mouse Ab, and the control was secondary Ab only. **(D) Detailed specificity in ABbA binding.** To test for cross reactivity with the non-Leb-binding paralog BabB protein (with unknown function), three strains – USU101 (https://www.ncbi.nlm.nih.gov/nuccore/NZ_CP032818.1) and J166 from central/southern Europe ^22^ and J99 of African phylogeny ^40^ (Figure 2B) – and their *babA*-minus mutants – were analyzed. Two spontaneous *babA*-minus mutants, USU101Δ*babA* and J166*ΔbabA,* isolated after passage from Rhesus macaques ^44, 63^, and a J99Δ*babA* genetic deletion mutant ^41^ were compared with their cognate BabA-positive parent strains. (***i***) ELISA wells coated with the three *H. pylori* strains and the corresponding *babA*-minus mutants were incubated with ABbA-IgG. Binding was detected using an HRP-conjugated anti-human antibody. **(*ii***) Immuno-blot detection of BabA by the strains USU101, J99, and J166. ABbA-IgG recognizes and binds BabA but does not recognize any bands/proteins in the BabA-negative strains. The lower affinity of J166 for ABbA binding is reflected in the binding of ABbA-IgG to BabA on the bacterial surface in the ELISA, but not to semi-denatured BabA on immunoblots. The ELISA was repeated three times, and each bar represents the mean of three values with the standard error. (***iii***) BabA expression was verified in an immunoblot with BabA-specific rabbit serum under denaturing conditions, where BabA was expressed in the original strains but not in the corresponding isogenic BabA mutants. (***iv***) BabB expression was verified in an immunoblot with BabB-specific rabbit serum under denaturing conditions, where all tested strains expressed BabB. The BabA protein was detected with VITE rabbit antibody ^14^, and BabB was detected by VIRA rabbit antibody diluted 1:6000 and secondary HRP-goat (anti-rabbit) antibody diluted 1:1000 (DakoCytomation, Denmark A/S). **(E) Affinity of ABbA-IgG. (*i*)**The ABbA-IgG binding affinity to recombinant soluble BabA (527 aa) devoid of the membrane-spanning hydrophobic beta-barrel domain was measured using isothermal titration calorimetry (ITC). **(*ii*)** Five separate ITC tests demonstrated an affinity of 10.2 ± 1.5 nM with one representative experiment of Kd = 12.9 nM shown in the figure. **(F)** ABbA-IgG was immobilized on a chip and tested by Biacore Surface Plasmon Resonance (SPR) for binding to soluble BabA. Three separate tests demonstrated a Kd = 1.7 ± 0.2 nM. Most likely, the functional binding strength of ABbA-IgG will be log-folds increased by the avidity effects gained in binding to BabA multimers on the *H. pylori* bacterial surfaces ^14^. **(G)** Immuno-electron microscopy demonstrated specific BabA immune staining of *H. pylori* by ABbA. Bacterial cells with *H. pylori* 17875/Leb (***i*, *ii***) were incubated with ABbA, and bound IgG was visualized with 10 nm gold-labeled protein A. The 17875*babA*-minus mutant was used as a negative binding control (***iii***), and the anti-E2 HCV envelope IgG1 was used as a negative antibody control (***iv***). The 17875*babA* mutant displayed only non-specific background binding similar to the isotype control antibody.

**Figure S5.**
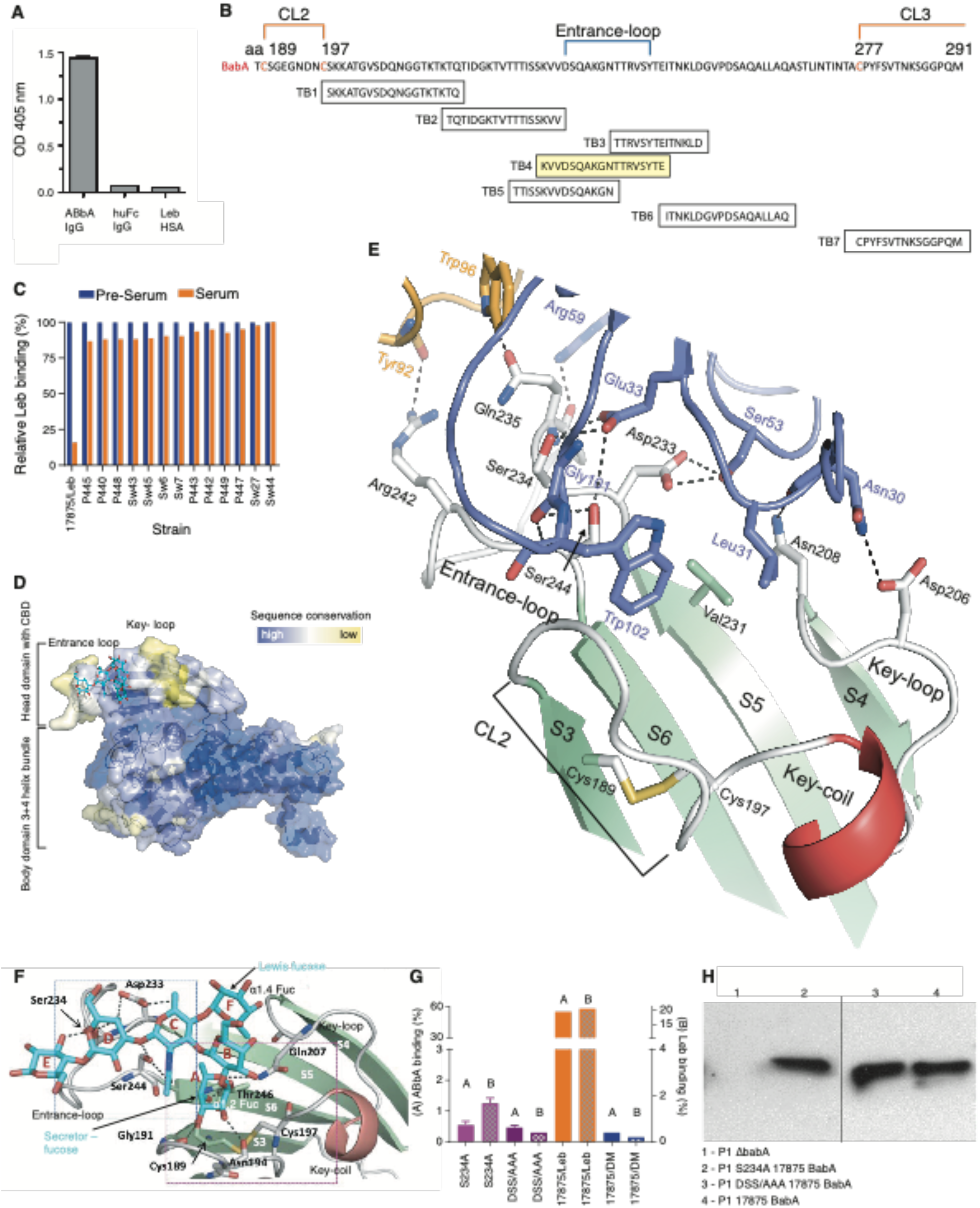
Identification of the structural binding epitope in BabA for the broadly blocking ABbA. (A) The 260 aa BabA_76-335_ polypeptide selected from the phage-display shotgun library was recombinantly expressed with its three disulfide loops. Binding was assessed by ELISA with ABbA-IgG and human Fc (IgG1) fusion protein (as the negative control) and Leb (as the receptor). The 260 aa BabA fragment comprises an adequate epitope for ABbA binding but does not provide sufficient structural stability and support for Leb binding. The 260 aa BabA fragment was identified as reviewed in ^64^. (B) Alignment of the seven synthetic peptides of the CBD domain from aa197 to aa291, where peptides TB2, TB3, TB5, and TB6 partially overlap peptide TB4. (C) Inhibition of Leb binding by the rabbit-sera TB4 of a series of *H. pylori* strains from Sweden (Sw) and Peru (P) and strain 17875/Leb. The TB4 serum, which was derived from the TB4 peptide of strain 17875/Leb, only efficiently inhibited Leb-binding by strain 17875/Leb and did not significantly inhibit Leb-binding of strains from geographically local Sweden or distant Peru. (D) The BabA adhesin exhibits extensive polymorphism with multiple amino acid substitutions (indicated in yellow) preferentially focused in the Entrance loop and the Key Coil/Loop in the CBD, which is located in the head domain ^14^. (E) For clarity, the co-crystal structure shows the ABbA-BabA interaction without Leb. Out of 15 bonds in total, the VL makes 3 bonds with the Entrance-loop and the VH makes the other 12 bonds, including the triple glycan mimicry (GM) domains. The BabA (grey loops and green β-strands) bound to ABbA-Fab (VH in lilac, VL in beige) with the amino acid residues that bind BabA indicated, including the hydrophobic W102 and L31 that substitute for the fucose residues. In addition, VH residues E33, S53, and G101 (in blue) form tight grips with the DSS triad. The VL also contributes with bonds to R242 and Q235. Q235 also binds to R59 in the VH and thus is the only BabA residue that binds to both the VH and VL. (F) Structure of the 17875 BabA-CBD bound to and co-crystallized with Leb (Leb shown in cyan; PDB: 5F7W). The six sugar rings of Leb are indicated by A–F, where A and F are the secretor fucose and Lewis fucose, respectively, making α1.2 and α1.4 bonds with the Gal (B) and GlcNAc (C), respectively. The D and E residues constitute the lactose core and together with GlcNAc (C) form H-bonds (dashed lines) with the DSS residues located in the BabA Entrance loop. The secretor fucose makes many bonds with the CL2 loop, whereas the Lewis fucose interacts with the hydrophobic V231 residue located in the immediate proximity of the DSS triad ^13^. (G) The triple DSS-to-AAA mutant and the critical single S234A DSS mutant of BabA were expressed in *H. pylori* strain P1Δ*babA,* where they demonstrated loss of both ABbA and Leb binding. Controls were the 17875/Leb BabA expressed in the P1Δ*babA* strain (positive control) and the 17875*babA1A2*-minus mutant (negative control). (H) BabA expression was verified by two (fused) immunoblots by BabA rabbit sera and showed that the expression level of BabA 17875/Leb in the *H. pylori* P1-DSS/AAA and P1-S234A mutant was similar to that of *H. pylori* P1 with the non-mutated BabA.

**Fig S6.**
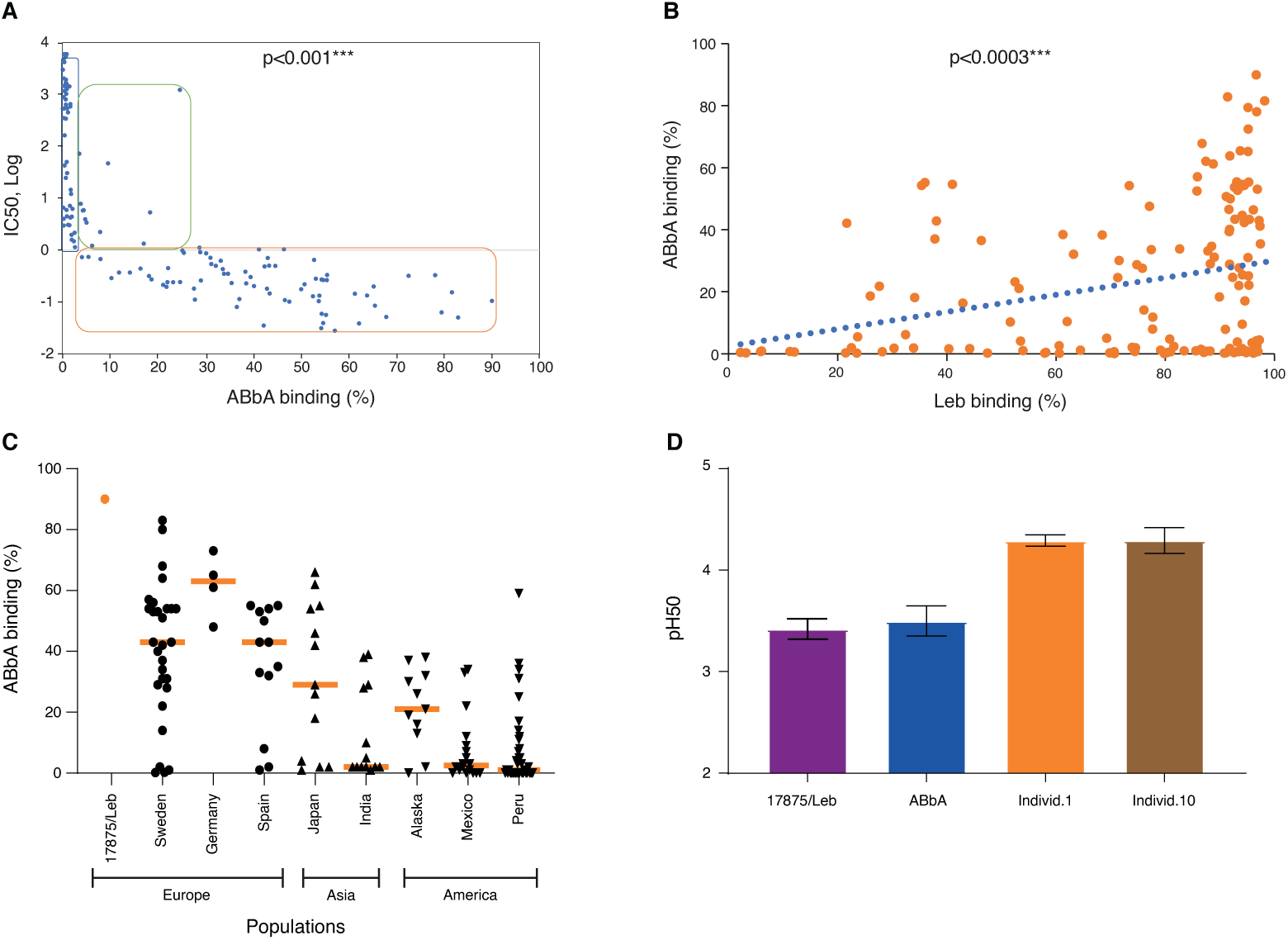
The ABbA global binding epitopes and their acid sensitivity in binding. (A) The ABbA-IgG binding strength showed a strong correlation with the ABbA IC50 (**Table S7**). Spearman’s rank correlation r = –0.881 (p < 0.001). (B) The ABbA and Leb binding strengths (**Table S7**) strongly correlate among global strains (r_s_ = 0.3, p < 0.0003). Thus, clinical isolates with higher binding strength for Leb most often bind ABbA-IgG with higher binding strength. (C) The ABbA-IgG binding strength is higher among European strains and also among strains from Japan, whereas strains from Southeast Asia (India) and North, South, and Latin America are lower in ABbA binding strength. (D) The acid sensitivity profiles, denoted as a pHgram, were determined by incubation of *H. pylori* with 125I-labled Leb and ABbA-IgG and Fc-purified IgG antibodies from individuals 1 and 10 in the pH 6–2 interval ^14^. The series of pH50s were repeated 3 times with SEMs. The test also showed that ∼1% of the purified serum IgG bound to *H. pylori*, i.e., 0.1 mg/mL out of 10 mg/mL IgG. With this new understanding that the bbAb/ABbA content is 1/10,000 part of the IgG pool (**Figure 4D**), we conclude that bbAb similarly to ABbA constitutes ∼1% of the serum IgG in circulation that binds to *H. pylori*.

**Video S1.**
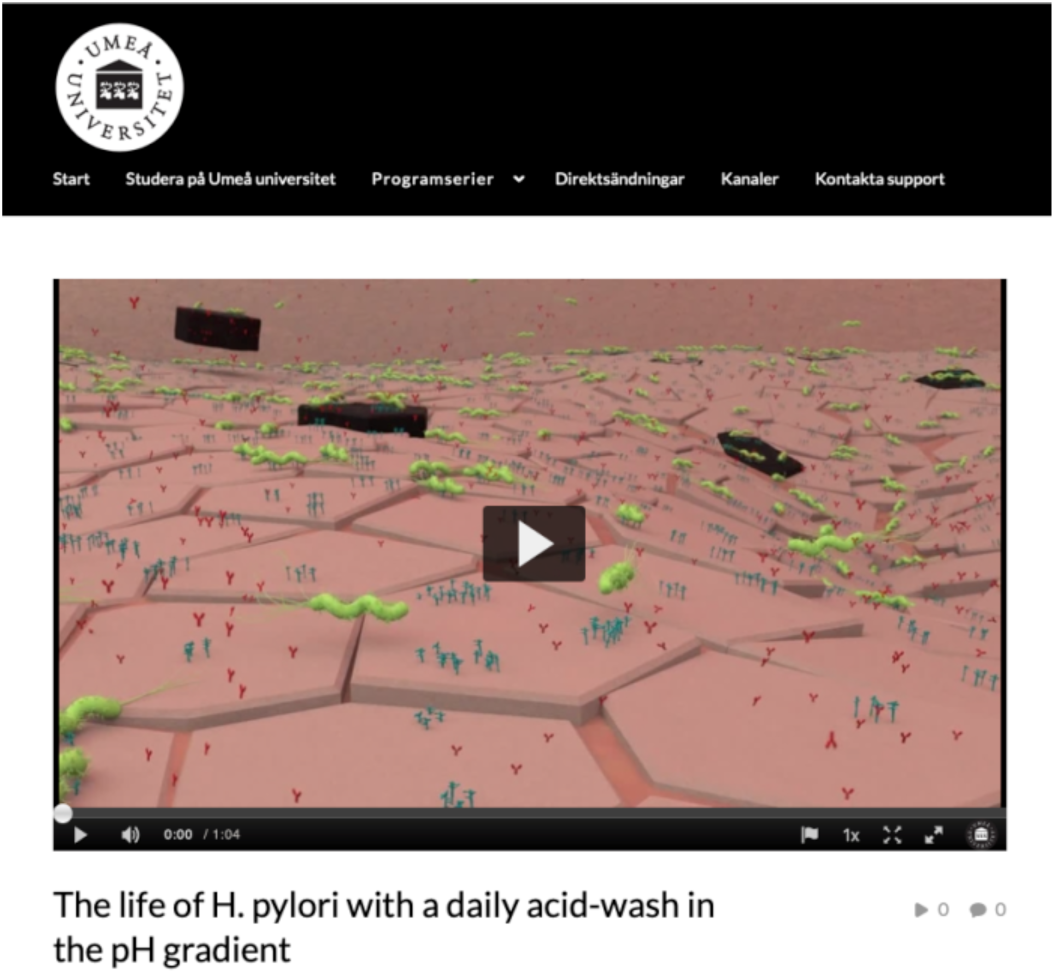
https://play.umu.se/media/t/0_bf1lh9gt The life of *H. pylori* with a daily refreshing acid wash.

The gastric onco-pathogen *H. pylori* lives in the stomach lining, referred to as the gastric epithelium. Close to the epithelial cells, the viscous mucus layer provides a bicarbonate buffer zone that protects the epithelial cell lining from the acidic gastric juice present in the stomach lumen just a third of a millimeter away. However, as the distance from the cell lining increases, the bicarbonate buffering fades to form a pH gradient that extends about half-way up the mucus layer. At this “altitude”, the impact of the very acidic gastric juice drops the pH back to pH1–2. The gastric juice is acidified by hydrochloric acid, which of course is too acidic for most microbes, *H. pylori* included. In this fascinating environment, *H. pylori* has adapted its life in the stomach to make life-long use of both the epithelial cell lining and the mucus layer pH gradient ^14, 26^. The more virulent *H. pylori* isolates bind the ABO blood group antigens (in blue) on the epithelial cell surfaces for firm bacterial attachment. In this prime location, *H. pylori* has full access to the necessary nutrients and iron leaching from the mucosal cells. However, pros come with cons, and the tight attachment exposes *H. pylori* to our immune system. The *H. pylori* infection is a major target of the humoral immune system, and the majority of carriers produce ELISA titers against *H. pylori* antigens. Hence, the *H. pylori* infections are exposed to high levels of antibodies (in red) that disseminate into the gastrointestinal tissues from the systemic circulation as well as from antibody-producing cells in both the stomach and gut lining. These antibodies bind to a plethora of *H. pylori* antigens such as the urease enzyme, LPS, heat-shock proteins, and the CagA effector protein. Although, the very best attempt possible by our immune system, the antibodies against *H. pylori* never manage protect against the *H. pylori* infection, which instead establish itself through adaptation and persists over the lifespan. The persistent infection is the cause of the chronic mucosal inflammation that in many millions of cases annually results in gastric disease, although of different degree of severity, ranging from mild and non-atrophic gastritis to “the silent killer” gastric cancer.

Our results suggest that the humoral immune system has found a way to balance the inflammation pressure by raising the bbAbs that prevent BabA adhesin from binding to the ABO/Leb glycans. Thus, the bbAbs block or reduce attachment of the *H. pylori* bacteria to the cell surfaces in the epithelial lining. The bbAbs bind to folded structures in the <u>c</u>arbohydrate <u>b</u>inding <u>d</u>omain (CBD) of the BabA attachment protein (the green hooks). This is illustrated by the *H. pylori* bacterial cell (located in the center) that is blocked by the bbAbs in binding the ABO glycans by the full set of BabA adhesins. Similarly, the bacterium to the left carries a multitude of bbAbs at its right pole and is barely hanging on in attachment to the cell surface ABO antigens.

To avoid detrimental consequences of the humoral defense and complement system activation, *H. pylori* can take advantage of the rapid gastric mucosal desquamation process, where epithelial cells are pushed into the mucus layer pH gradient. This is a natural, essential, and innate immune protective process and the epithelial cells undergo rapid turnover every 2–3 days and are continuously replaced with new cells proliferating from the stem cells located in the column-palisade just below. The protective mucus layer is renewed even faster, about twice per day. If these processes were irreversible and sufficient, the desquamation process would move and transport all attached *H. pylori* bacterial cells to the gastric juice where the bacteria would be destroyed by the acidic environment. If this were the case, the *H. pylori* infection would be naturally eradicated in a week or maybe two. But this does not take place and instead *H. pylori* adapts to the host and successfully persists for the lifetime of the individual.

Our new results suggest that *H. pylori* does not merely and passively adapt to the desquamation process but instead actively makes good use of it as a way to escape the humoral immune system. As the *H. pylori* cells that are attached to desquamating cells move upwards and through the mucus layer, they soon reach their first rejuvenation region at ∼pH 4.5, where the majority of acid-sensitive antibodies and complement molecules are both de-attached and inactivated. After about 2 h and further out in the mucus layer, at the slightly lower pH 3.5, the *H. pylori* cells will rid themselves also of the more acid-resistant bbAbs.

But how do the *H. pylori* manage to detach from the desquamating cells? The BabA adhesin binds to the ABO/Leb antigens on the cell surface similarly to the glycan mimicry of the bbAbs. During the millennia of adaptation to the human gastric mucosa, the main attachment protein BabA has evolved to be acid sensitive in its binding to ABO/Leb blood group glycans. At the low pH 3.5, the majority of BabA no longer bind, and the *H. pylori* will come loose from the desquamating cells. Once free, “acid-washed”, and again fully motile, *H. pylori* uses its chemotactic sensors and flagella (Goers Sweeney *et. al, Structure*, 2012) to propel itself along the pH gradient from the epithelial cells back to the buffered epithelial surface by its reported rapid motility in the mucus zone ^65^, *H. pylori* can in a short time return to the less acidic parts of the epithelial lining, where the bacteria cells can again reattach in a continuous recycling process. Thus, by taking advantage of the pH gradient, the *H. pylori* infection can escape the humoral effector molecules of the immune system and, in addition, can recycle the infection through natural bio-selection of those bacterial cells that have introduced mutations or induced or modified expression patterns that are the best suited for life in the local gastric environment. This also allows for adaptation to changes over the lifetime of the host. However, long-term infection by adherent *H. pylori* that cause chronic mucosal inflammation constitute the critical risk for gastric cancer development.

## Supplemental Tables

**Table S1A, related to Figure 1 and Figure S1, Figure 2 and Figure S2 and Figure 3.**
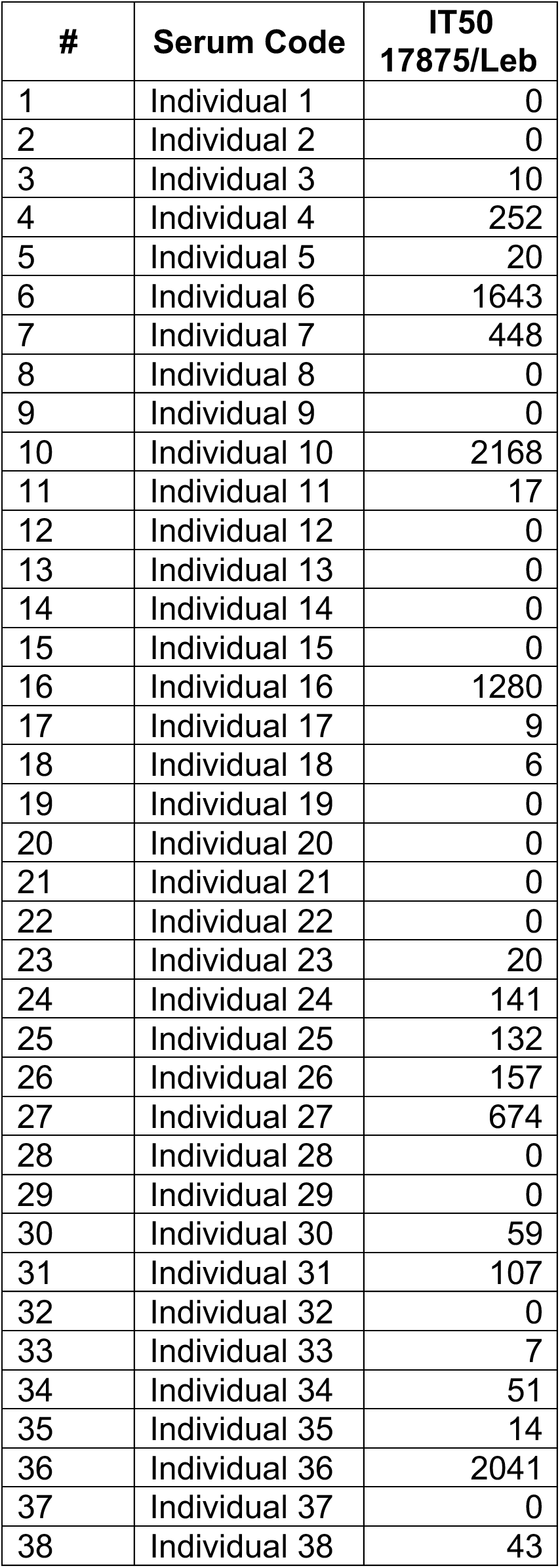
Characteristics of *H. pylori* ELISA-positive individuals from Karolinska University Hospital, Sweden.

**Table S1B related to Figure 1 and Figure S1 and Figure 3.**
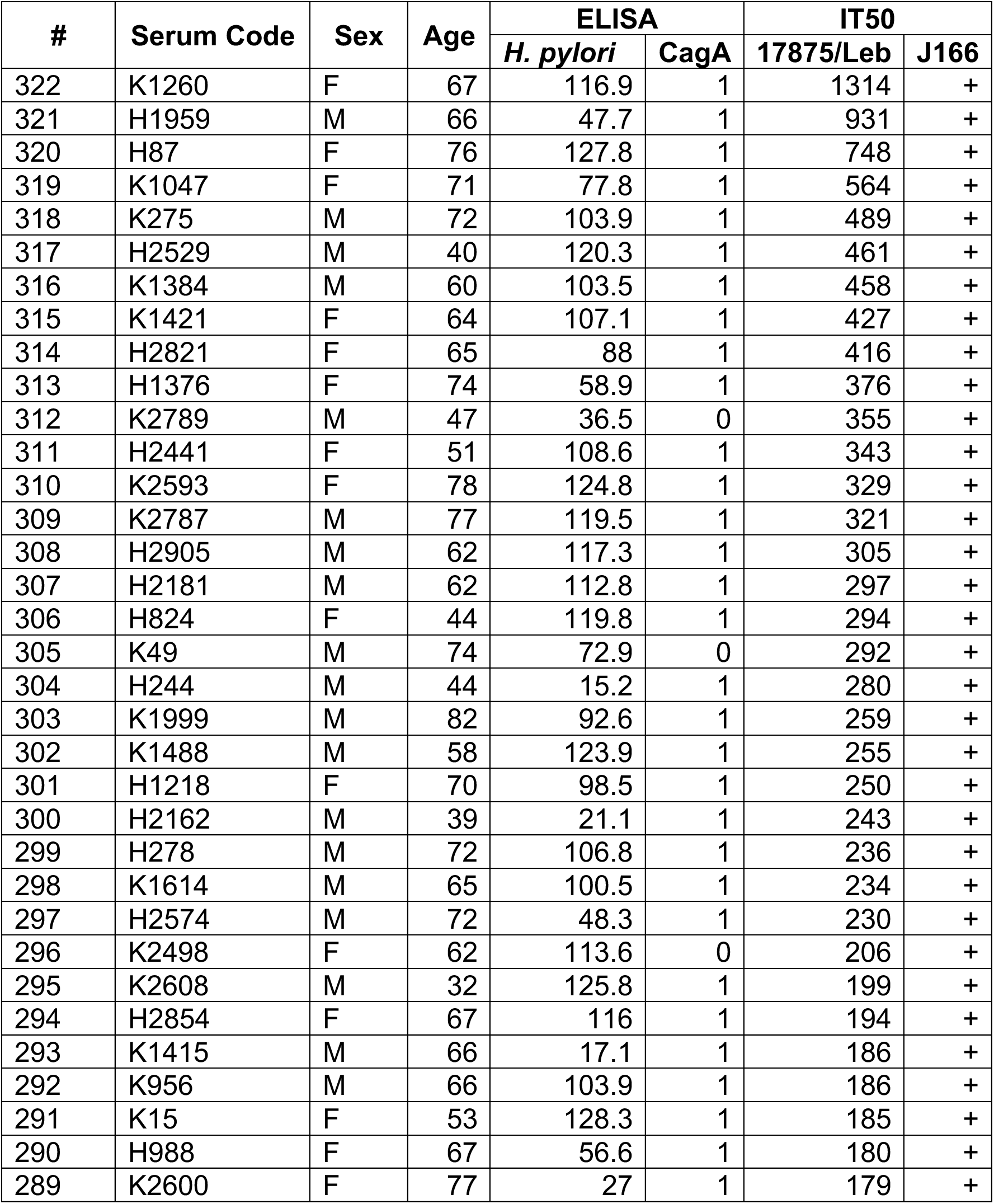
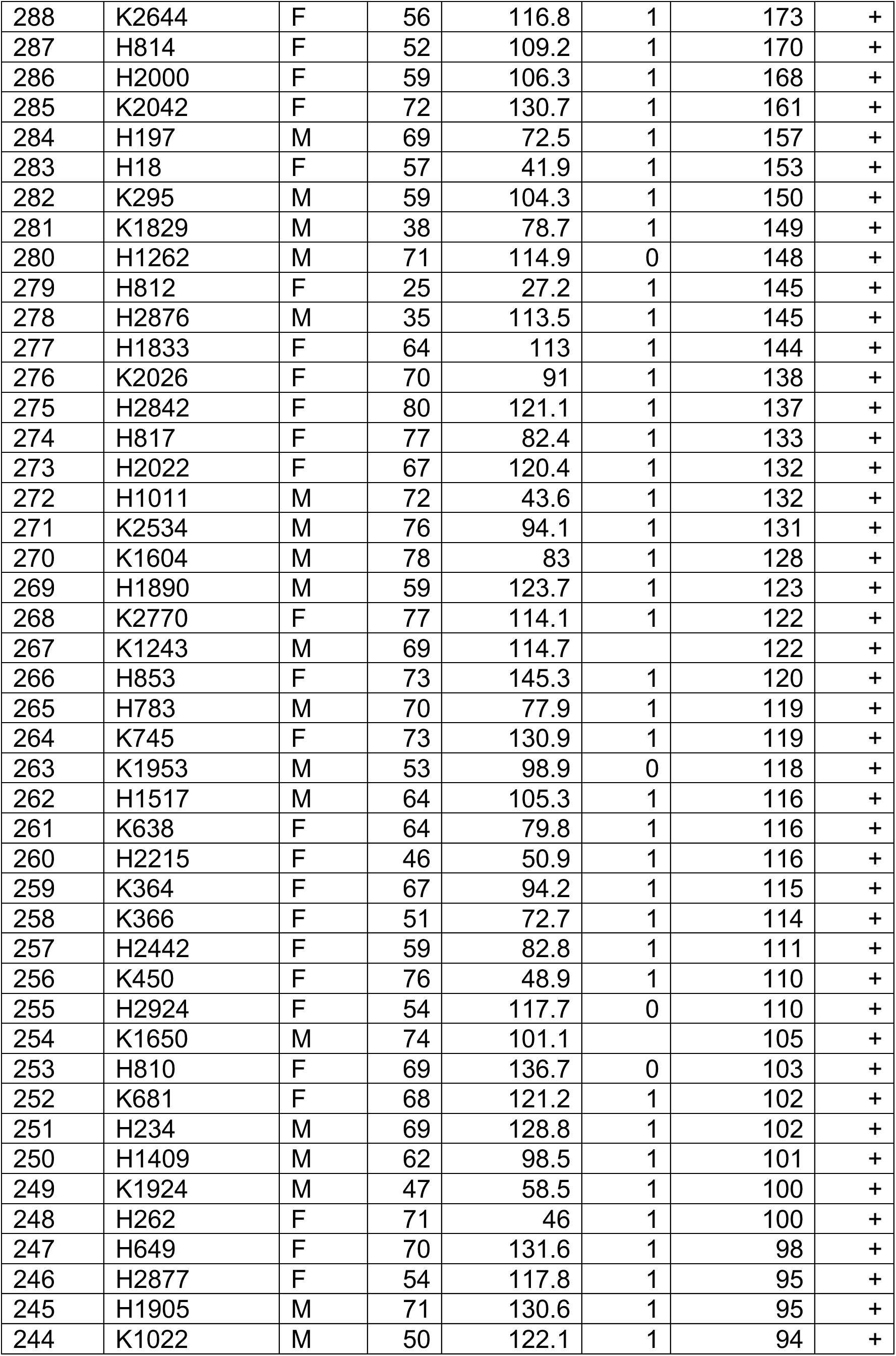
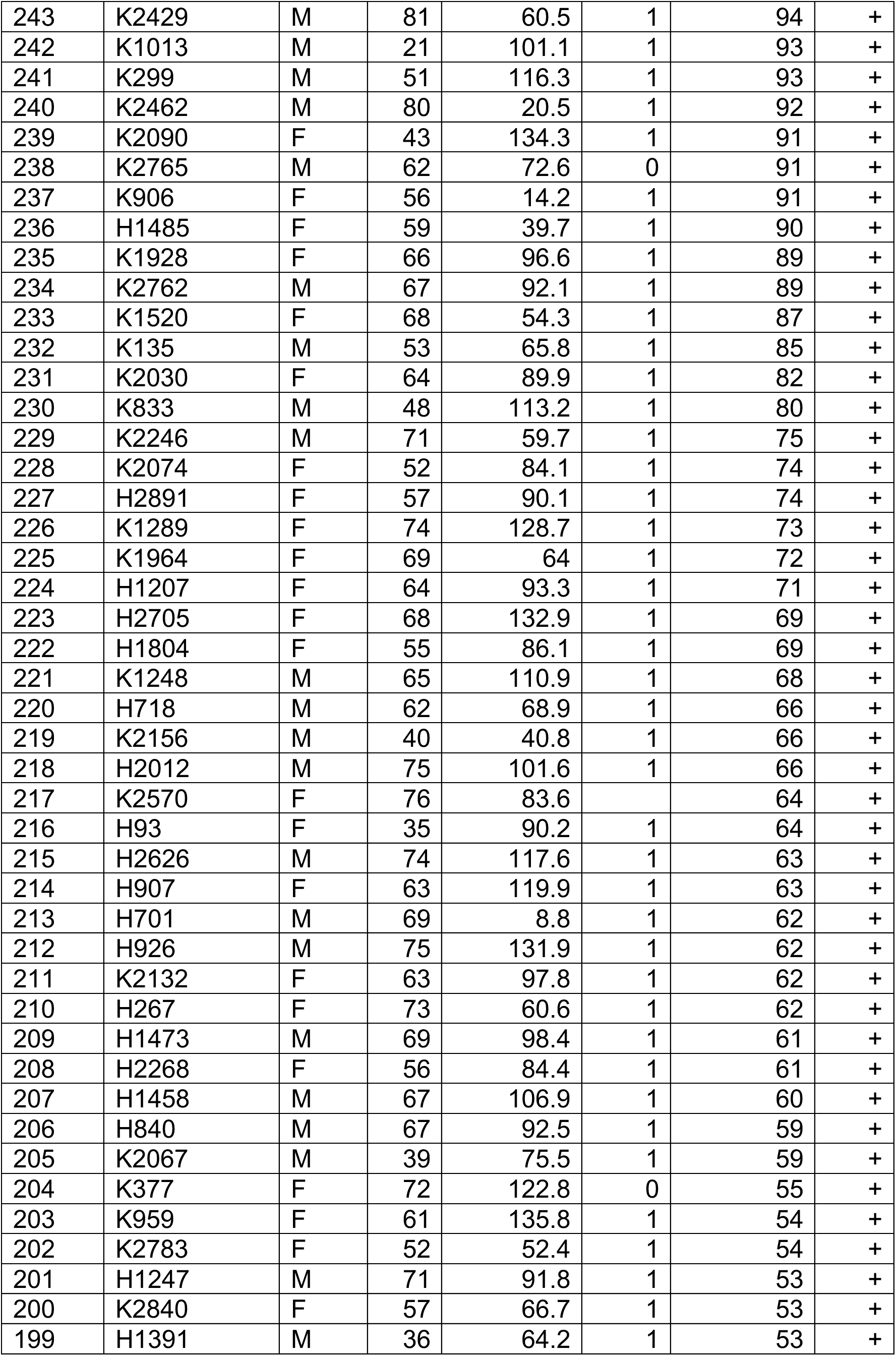
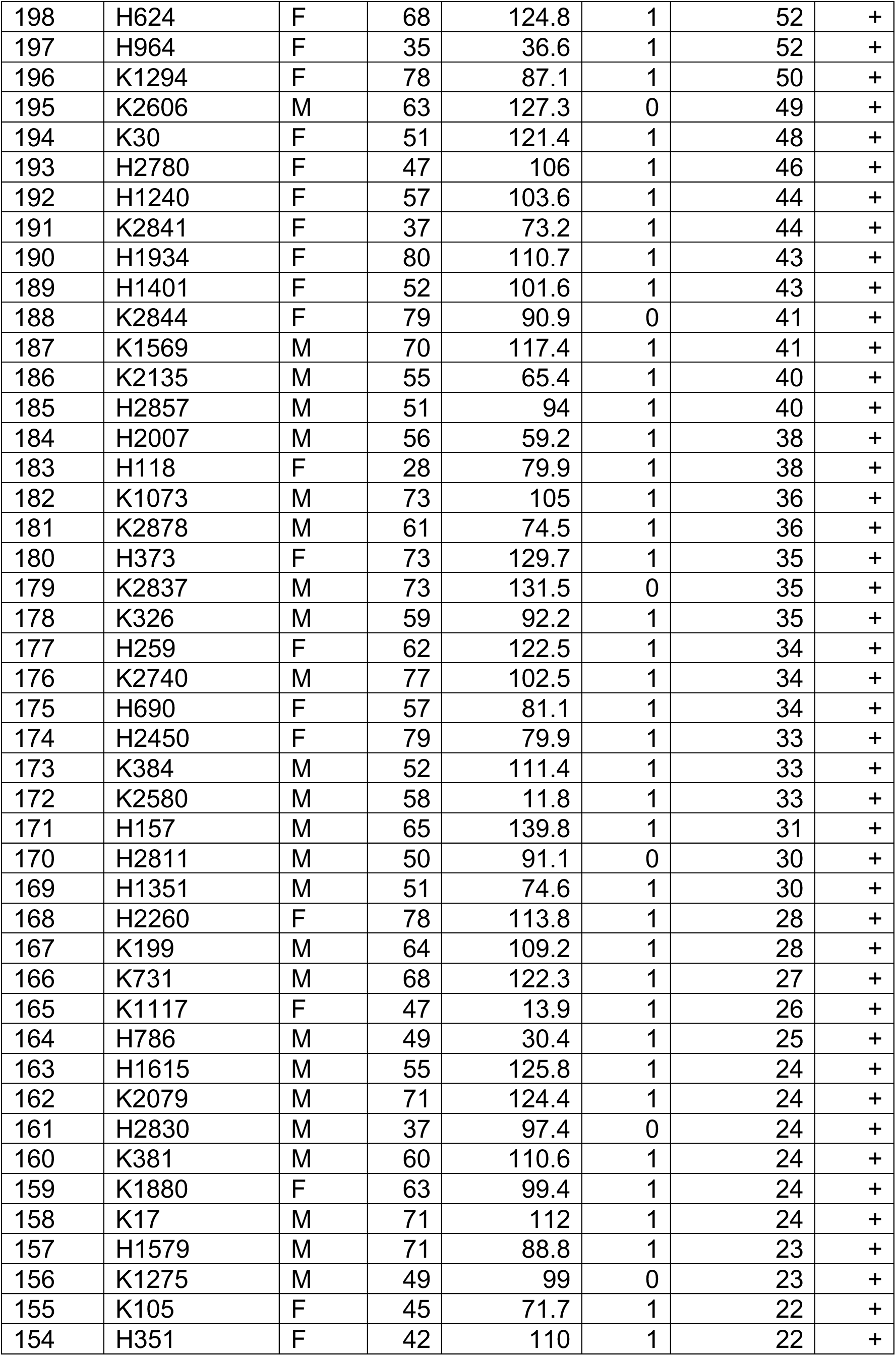
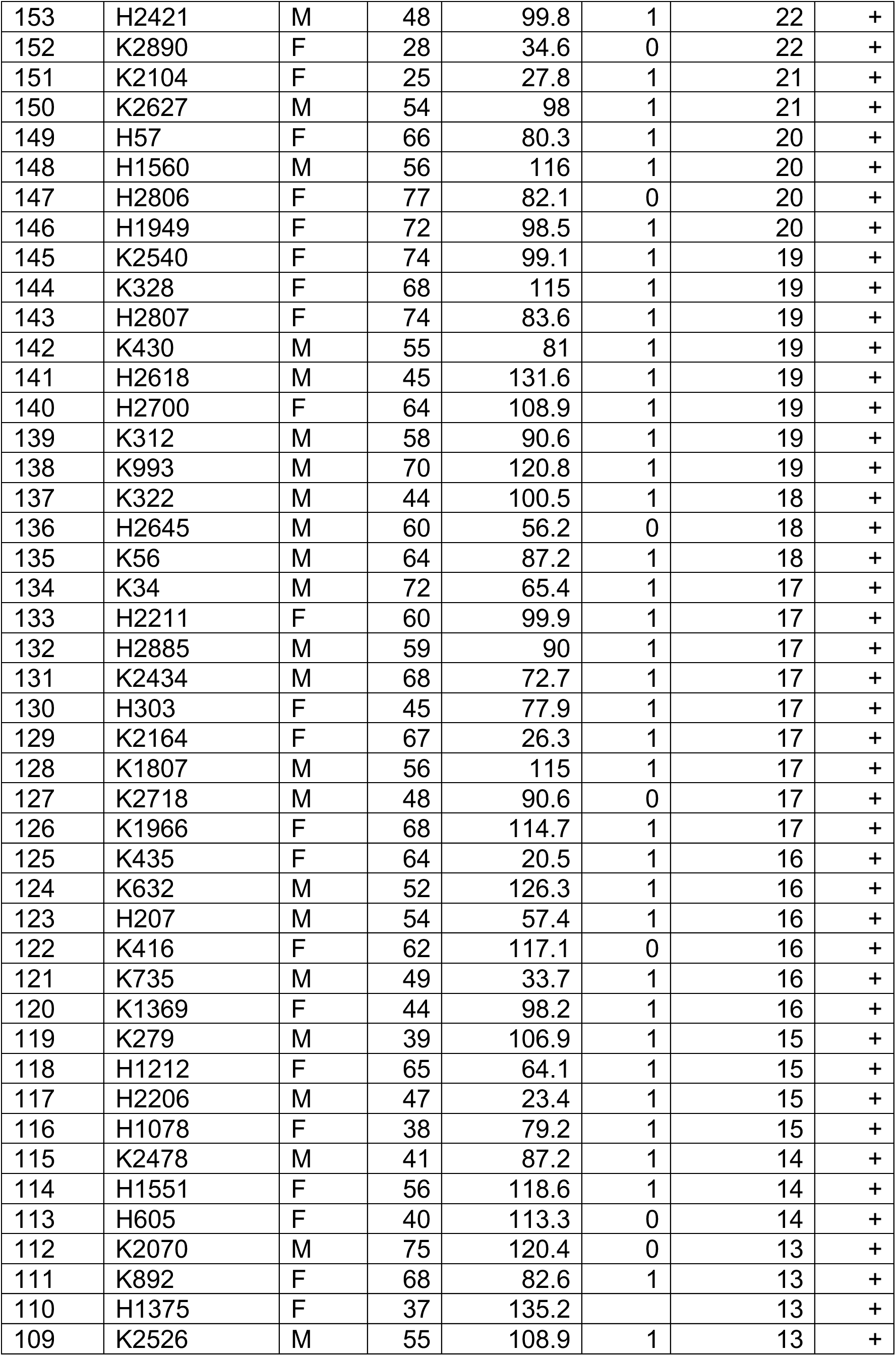
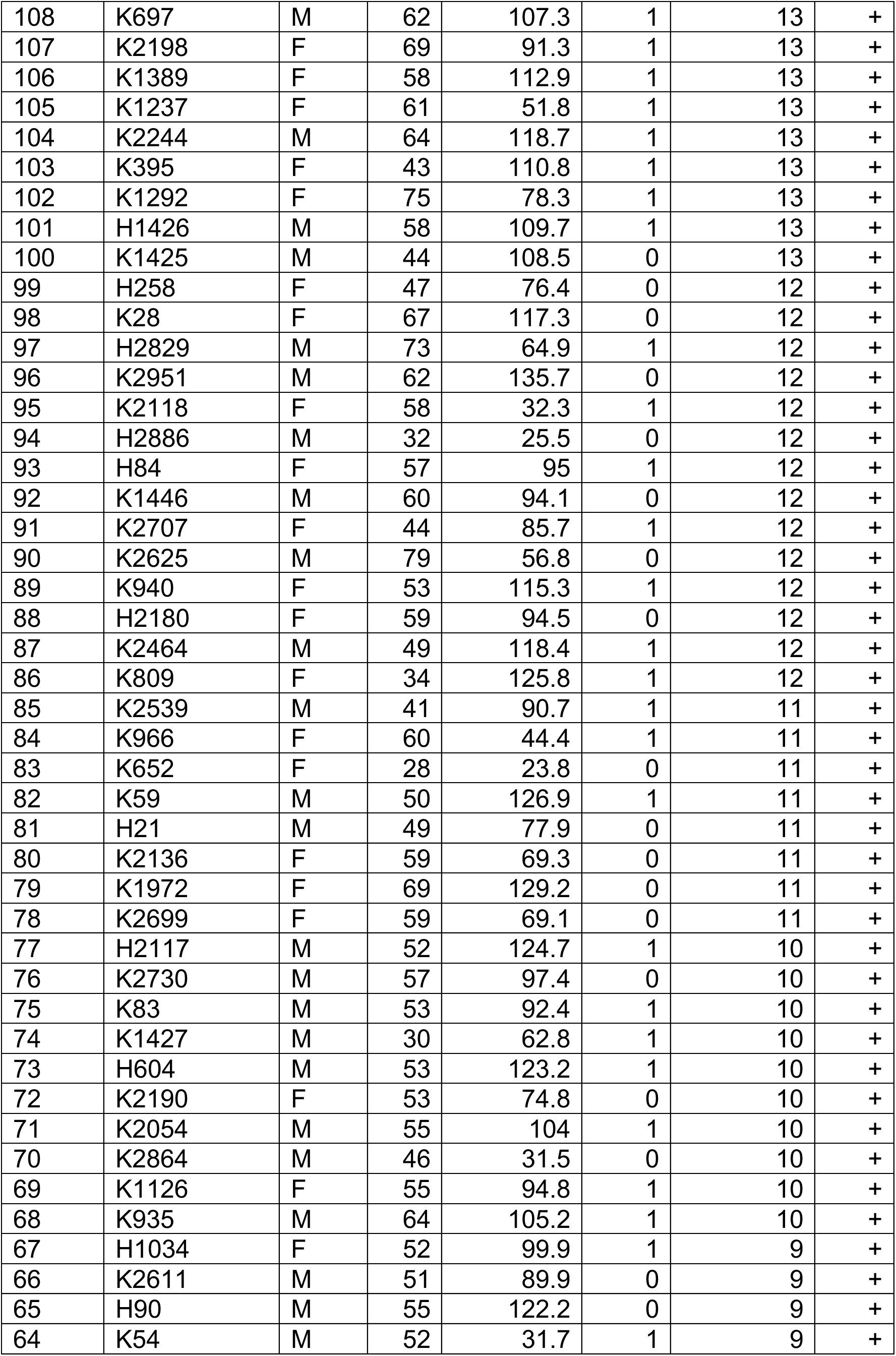
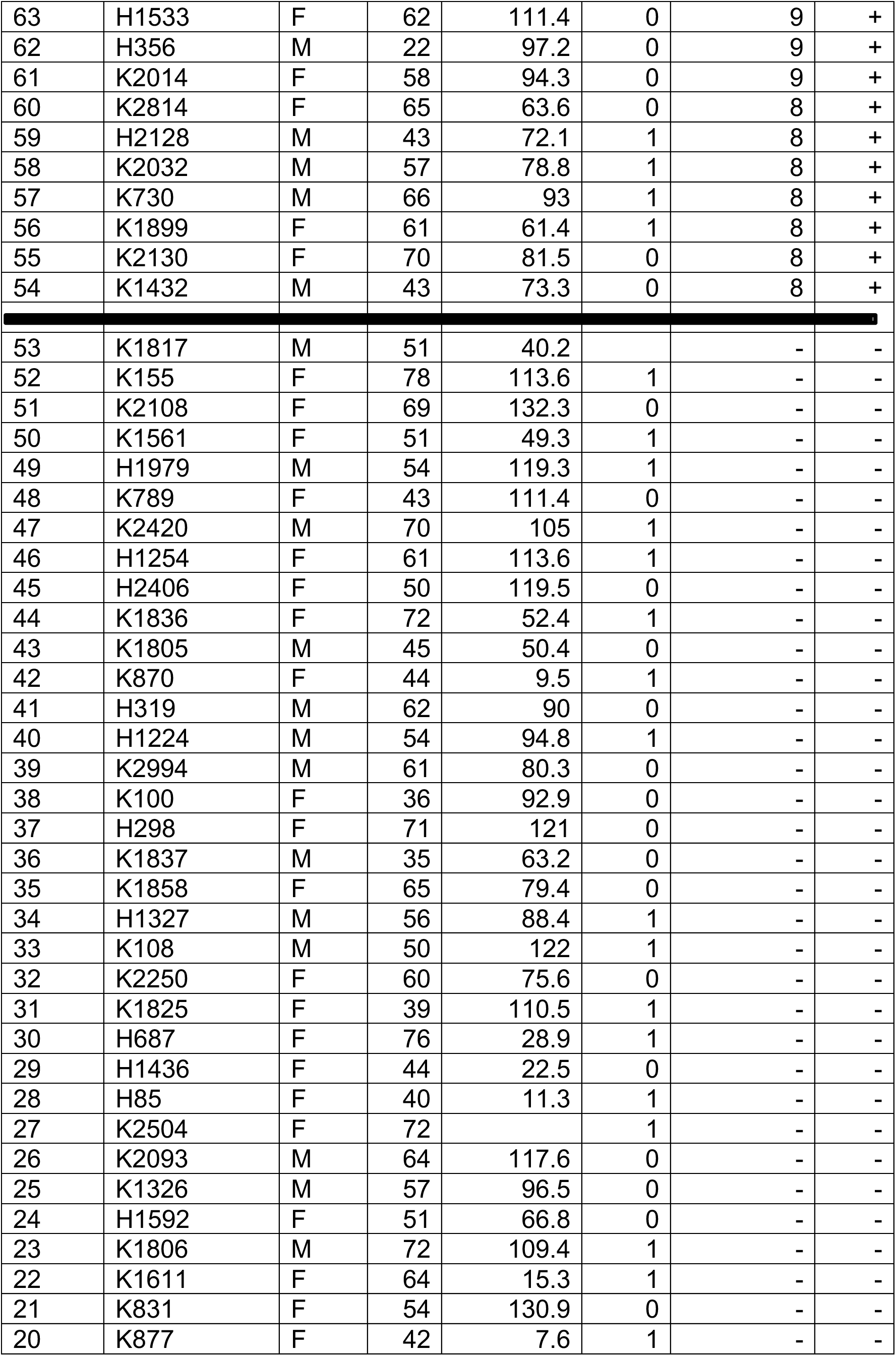
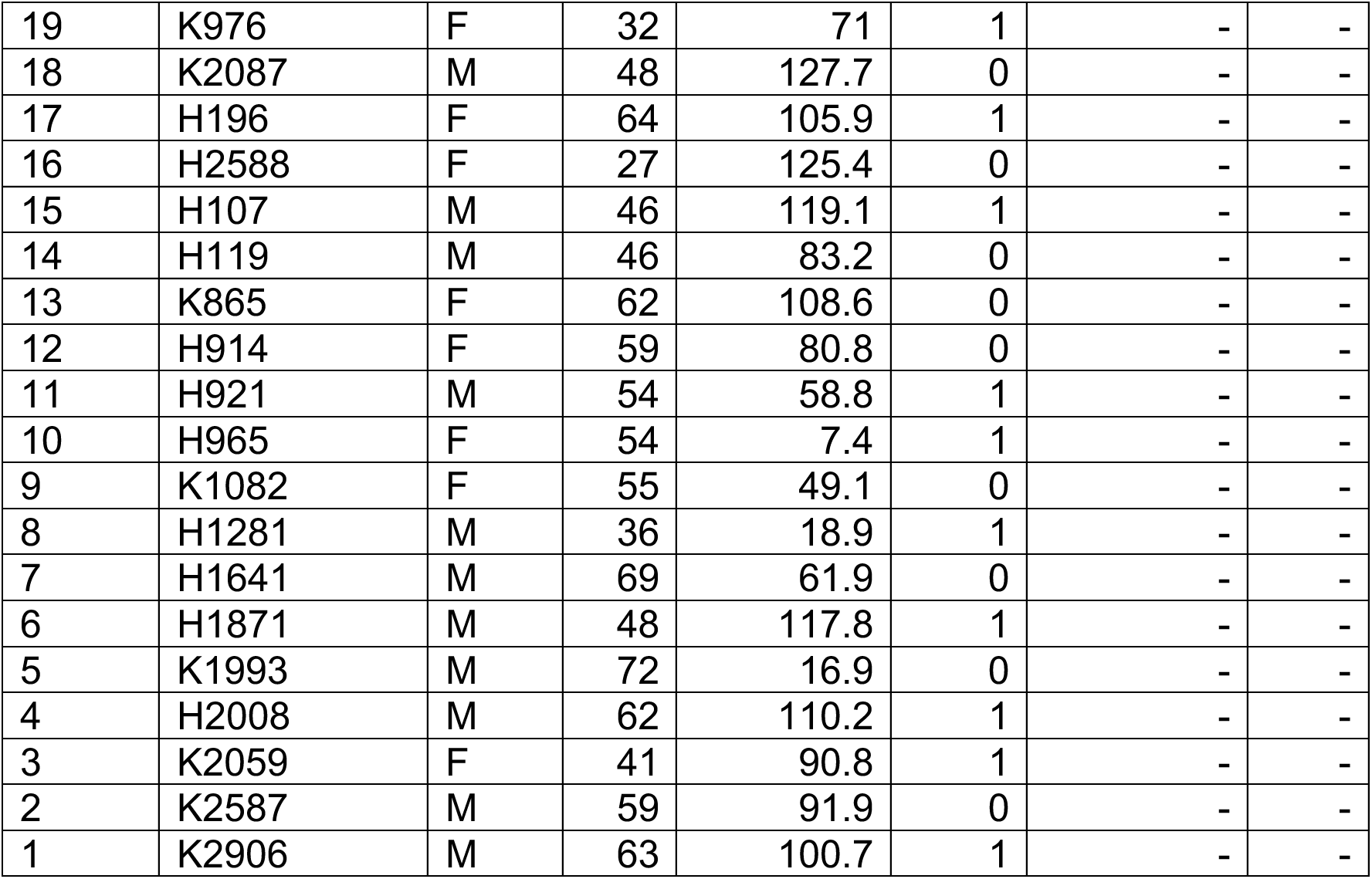
Characteristics of Individuals from Kalixanda Study, Sweden. The serum samples are displayed according to IT50s tested with *H. pylori* strain 17875/Leb. In addition, *H. pylori* strain J166 identified the series of IT50 titers as positive (+) or negative (–). The location for the background level (the start of the positive IT50 sample is indicated by the horizontal bar) was calibrated with sera from ELISA-negative individuals using *H. pylori* J166.

**Table S1C related to Figure S1 and Figure 3.**
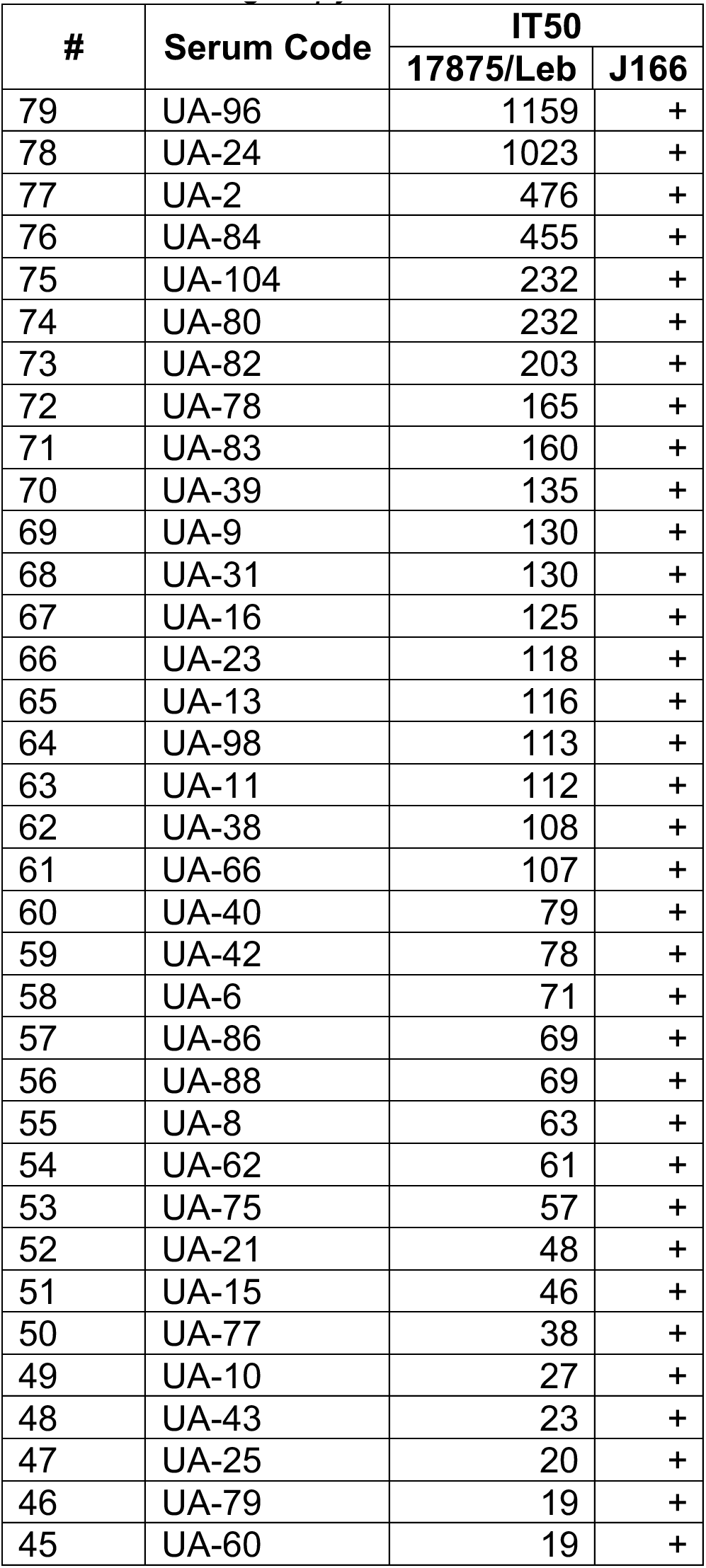
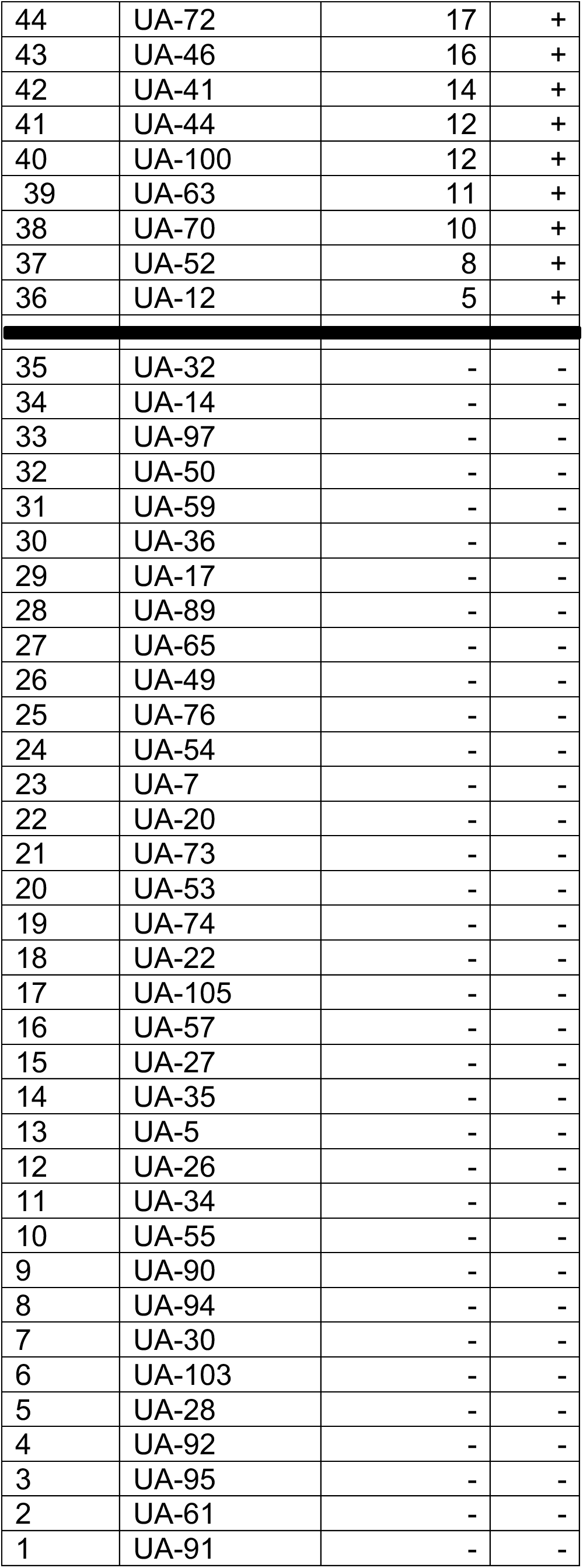
Characteristics of *H. pylori* ELISA-positive patients from Sumy Regional Clinical Hospital, Ukraine. The serum samples are displayed according to IT50s tested with *H. pylori* strain 17875/Leb. In addition, *H. pylori* strain J166 identified the series of IT50 titers as positive (+) or negative (–). The location for the background level (the start of the positive IT50 sample is indicated by the horizontal bar) was calibrated with sera from ELISA-negative individuals using *H. pylori* J166.

**Table S1D related to Figure S1 and Figure 3.**
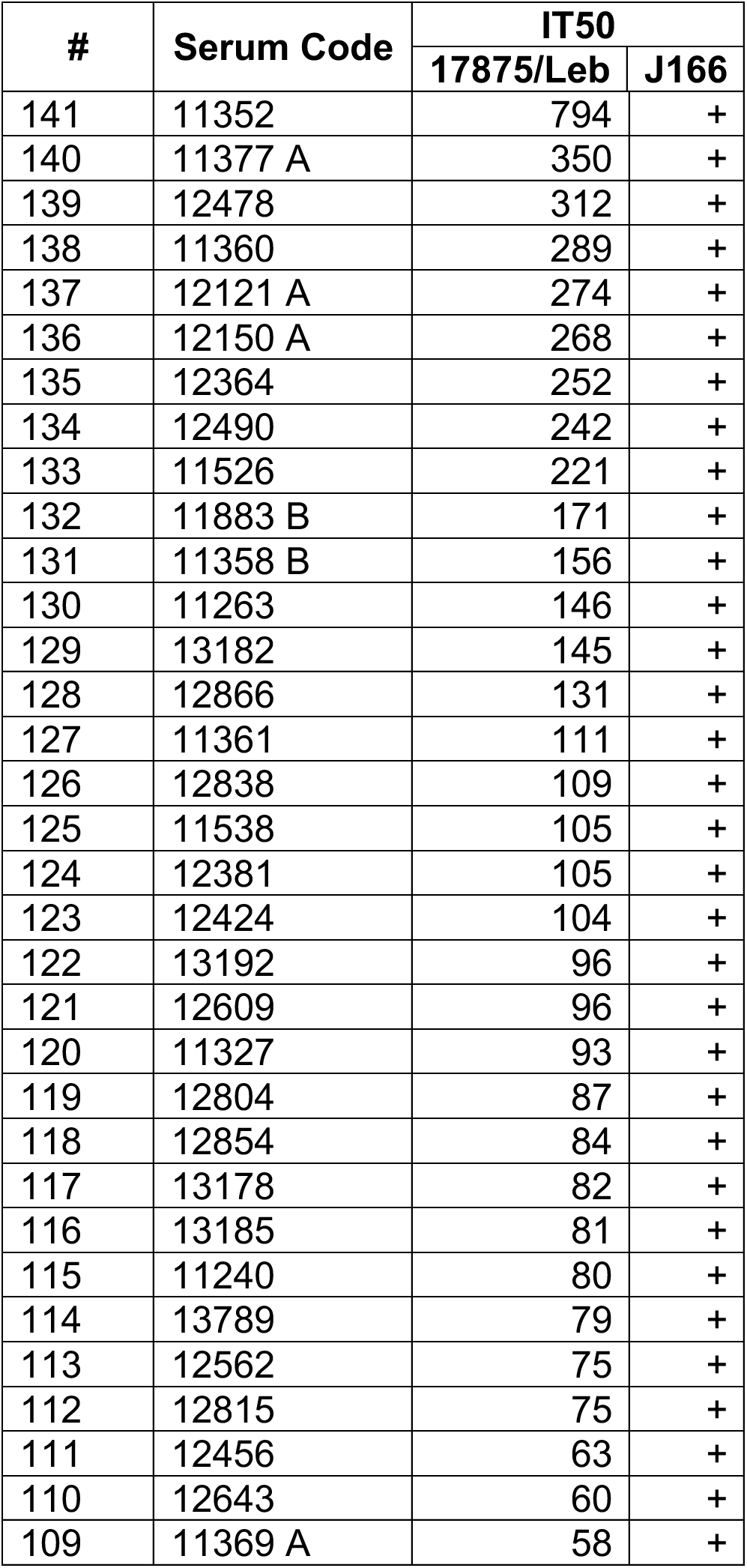
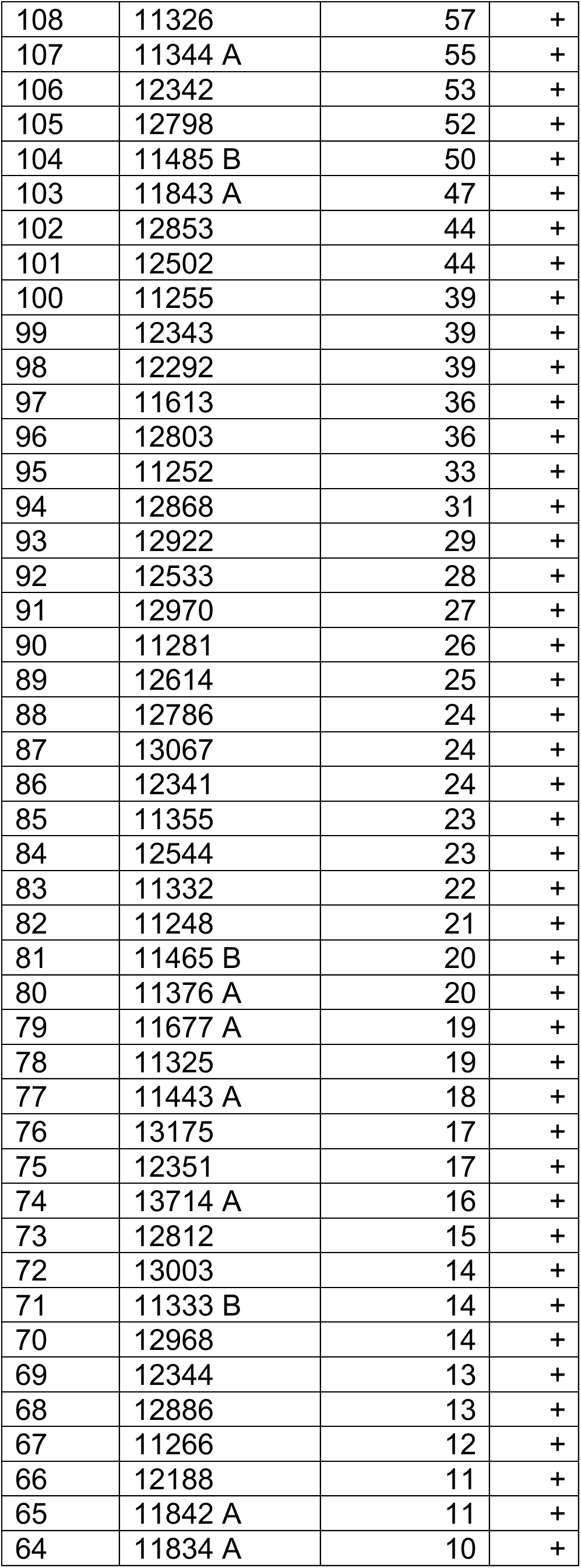
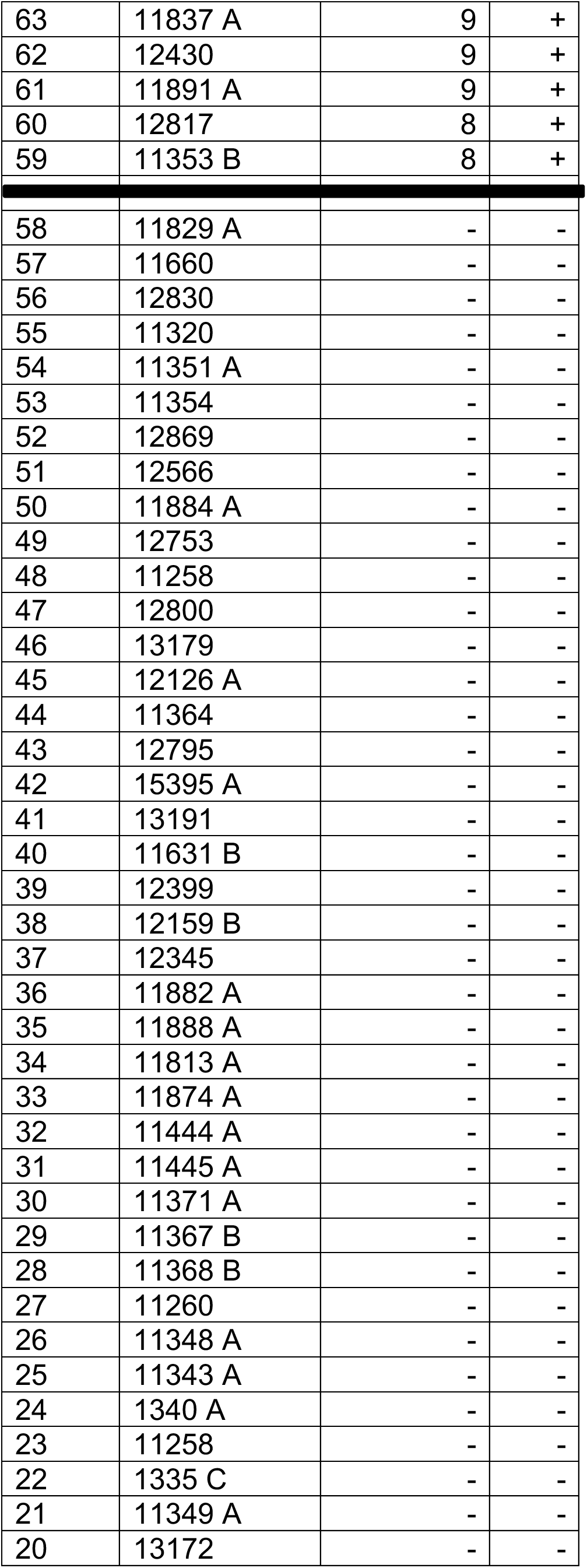
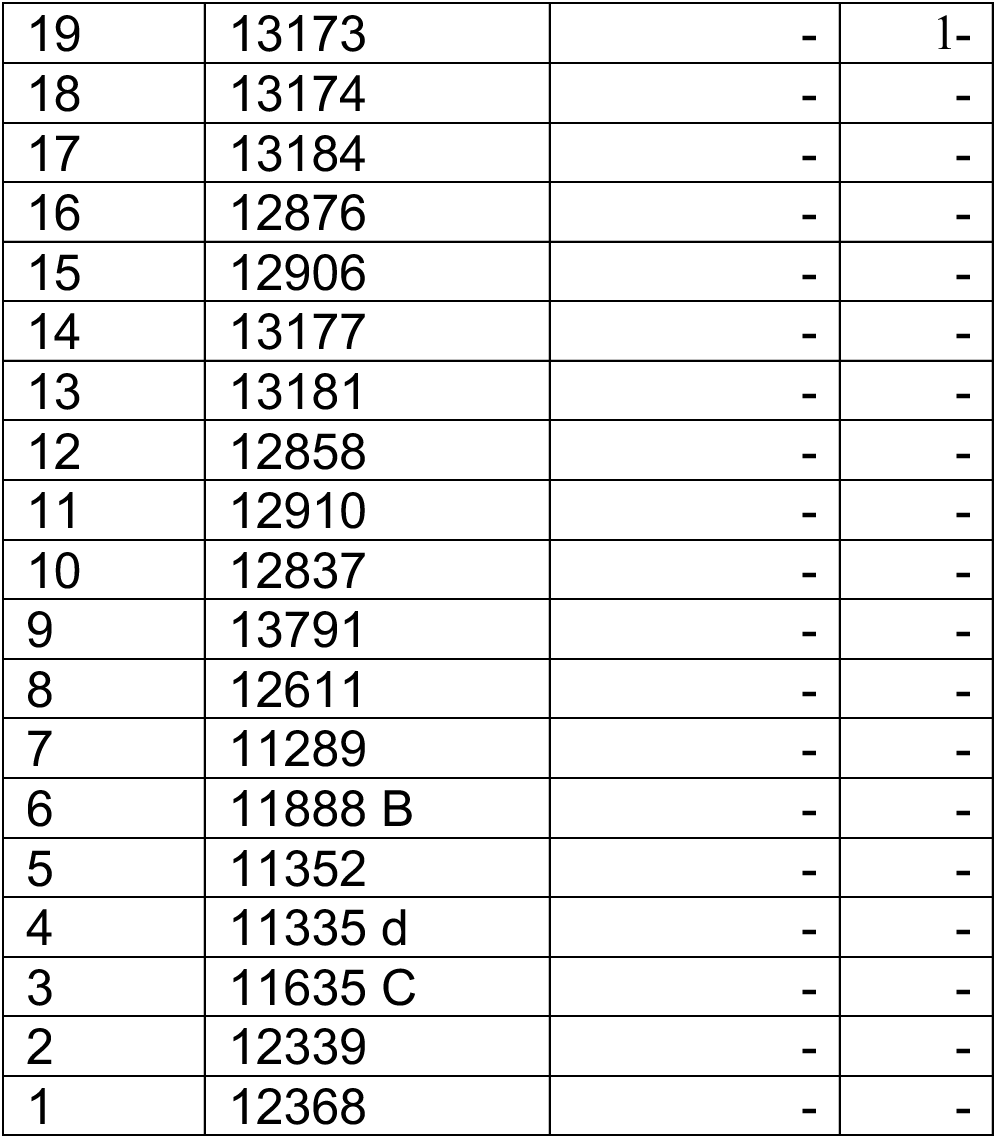
Characteristics of *H. pylori* ELISA-positive patients from Baylor College of Medicine, Houston, TX, USA. The serum samples are displayed according to IT50s tested with *H. pylori* strain 17875/Leb. In addition, *H. pylori* strain J166 identified the series of IT50 titers as positive (+) or negative (–). The location for the background level (the start of the positive IT50 sample is indicated by the horizontal bar) was calibrated with sera from ELISA-negative individuals using *H. pylori* J166.

**Table S1E related to Figure S1 and Figure 3.**
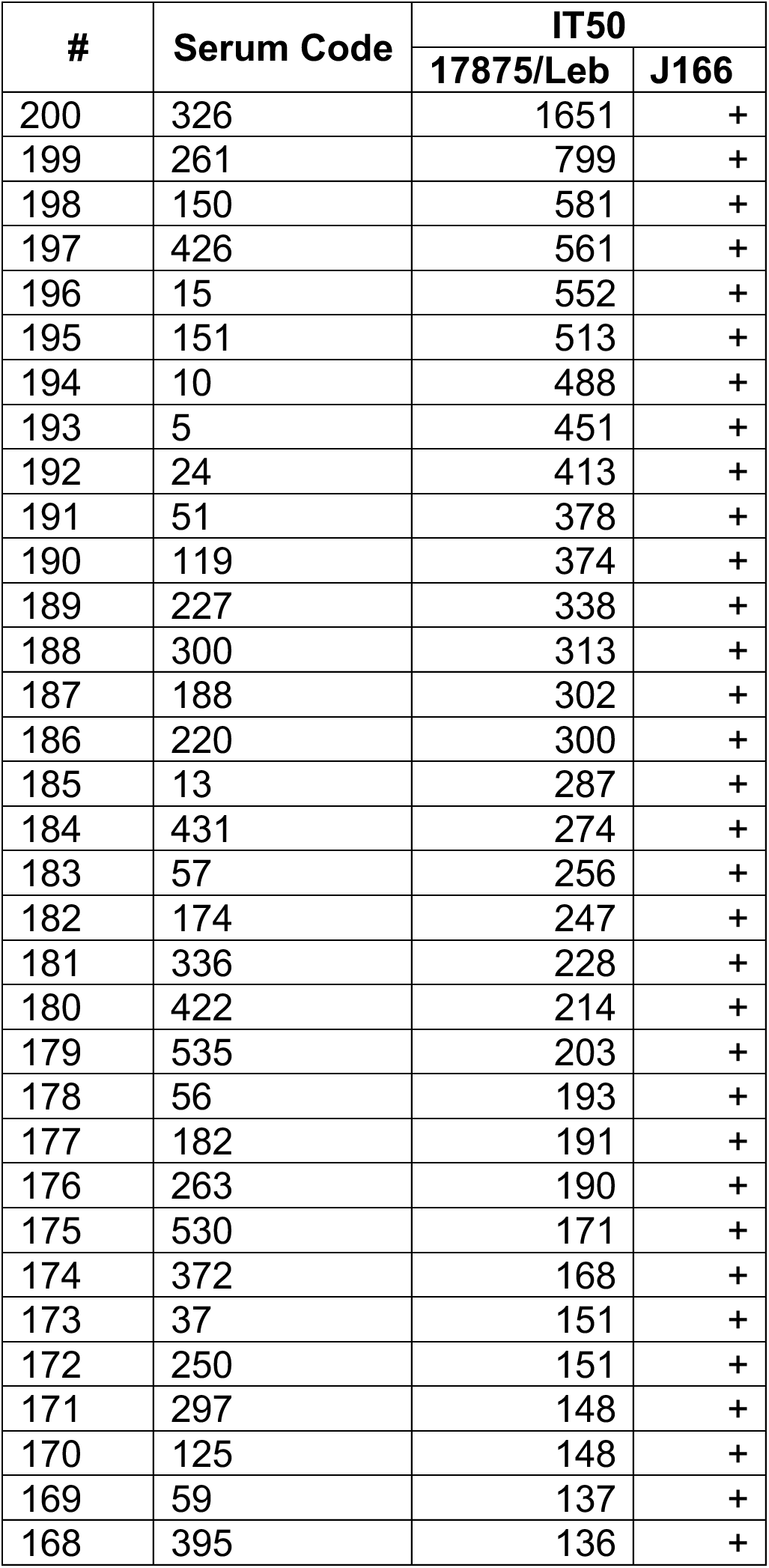
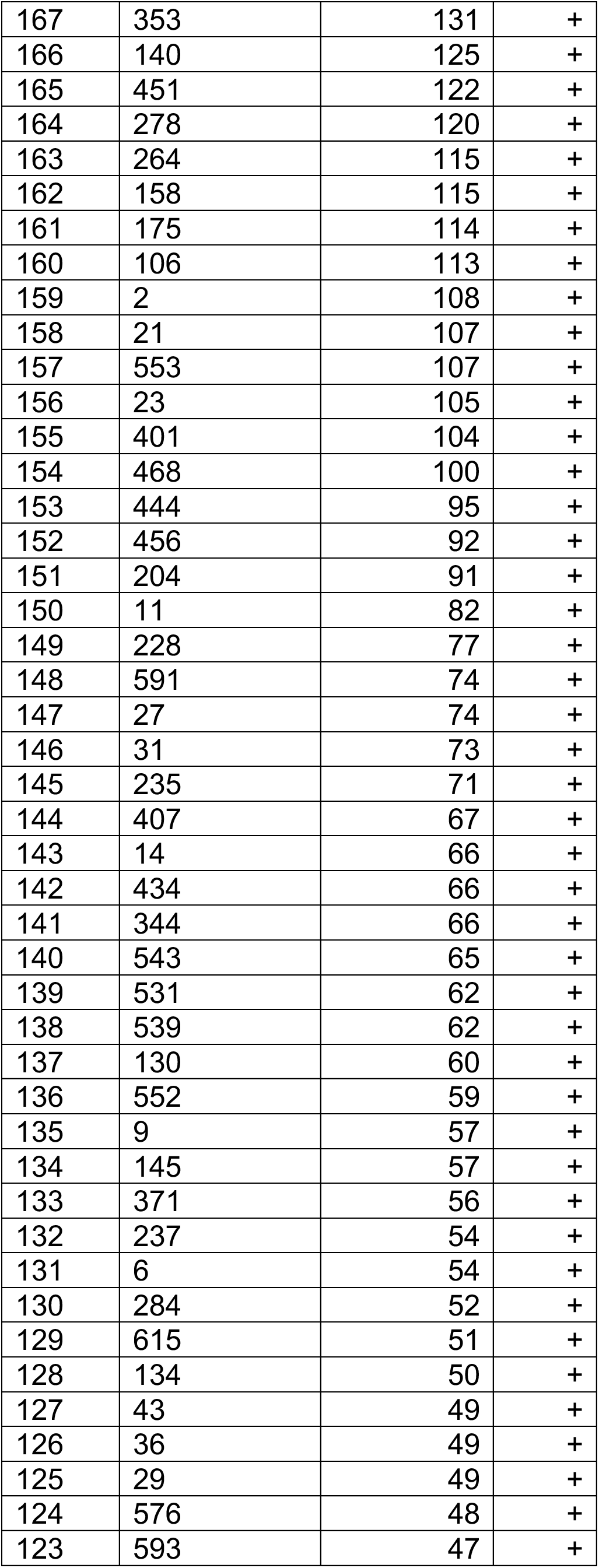
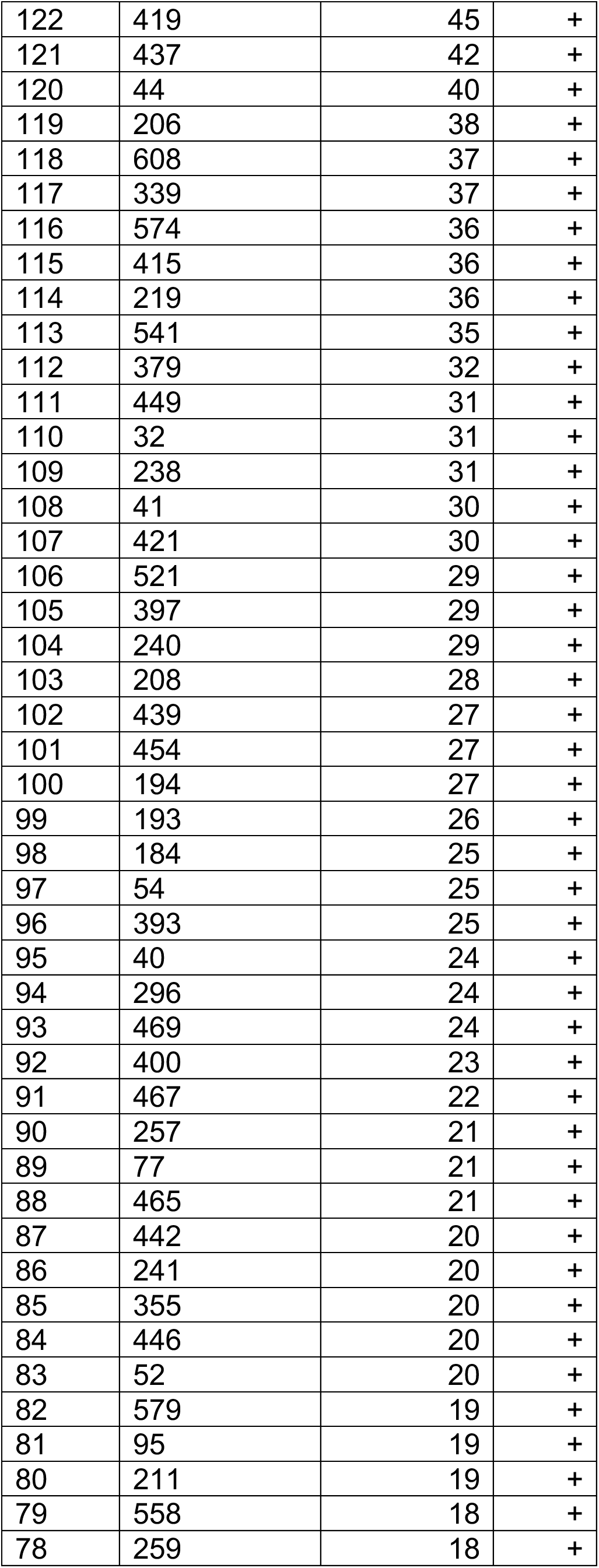
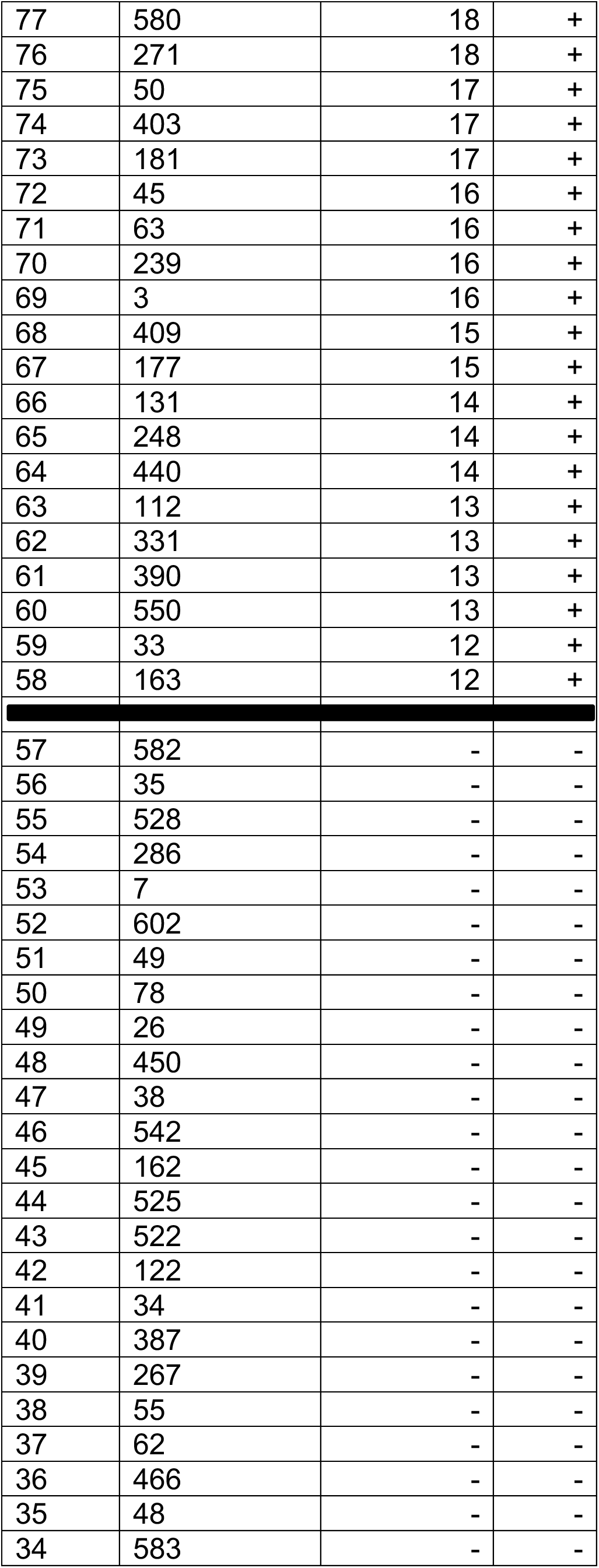
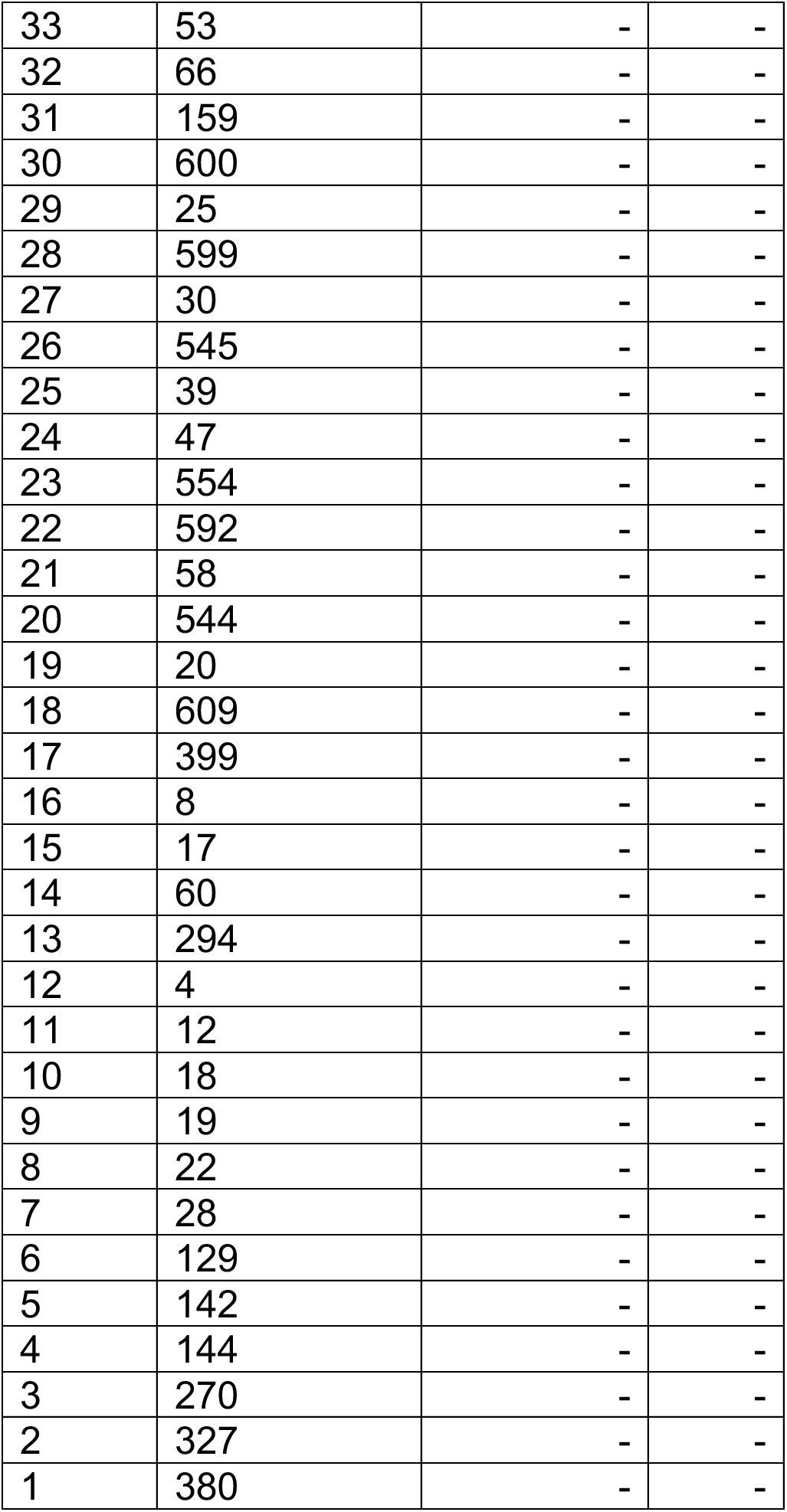
Characteristics of *H. pylori* ELISA-positive patients from Mexico UMAE Pediatria, Mexico City, Mexico. The serum samples are displayed according to IT50s tested with *H. pylori* strain 17875/Leb. In addition, *H. pylori* strain J166 identified the series of IT50 titers as positive (+) or negative (–). The location for the background level (the start of the positive IT50 sample is indicated by the horizontal bar) was calibrated with sera from ELISA-negative individuals using *H. pylori* J166.

**Table S1F related to Figure 1.**
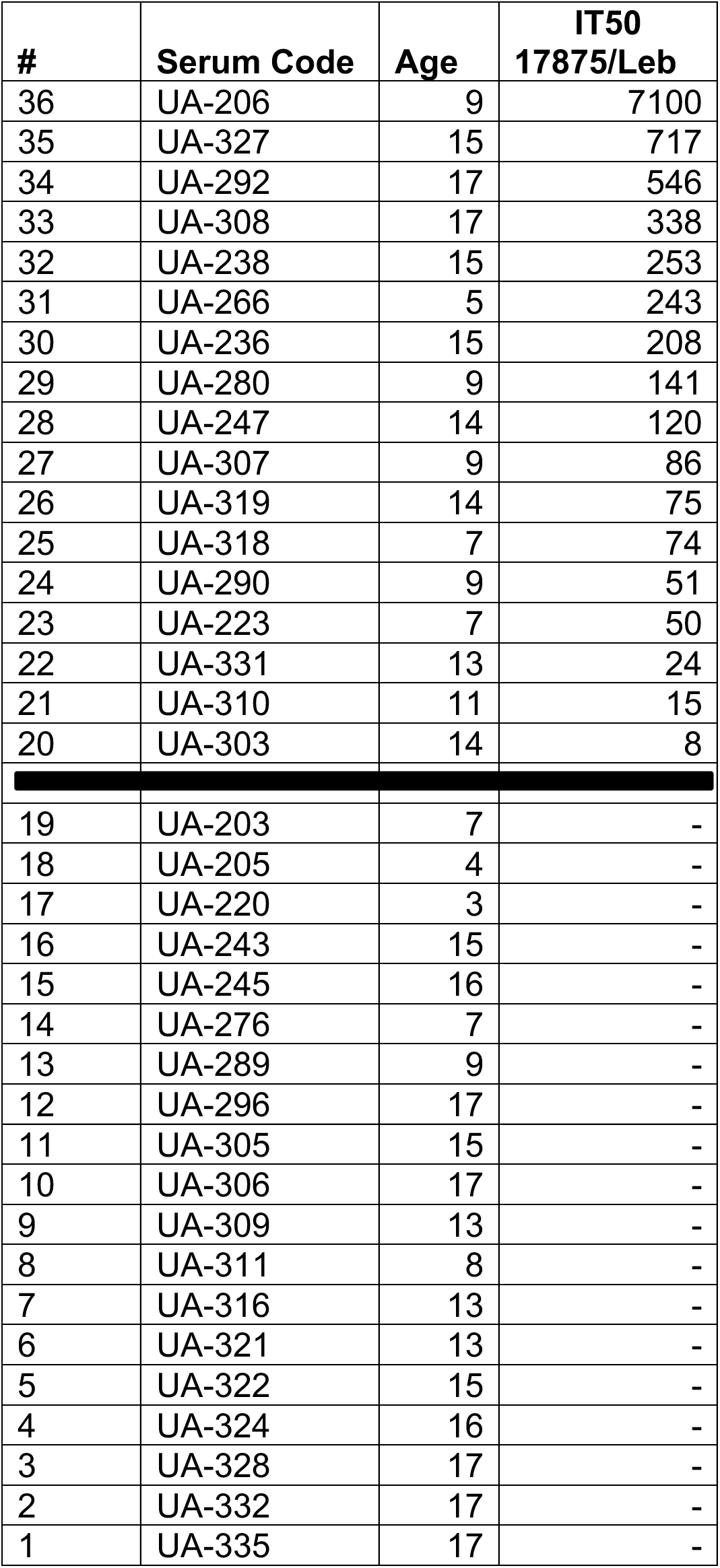
Characteristics of *H. pylori* ELISA-positive patients from Regional Children Hospital and St. Zinaida City Children Hospital, Sumy, Ukraine.

**Table S2A related to Figure 3A.**
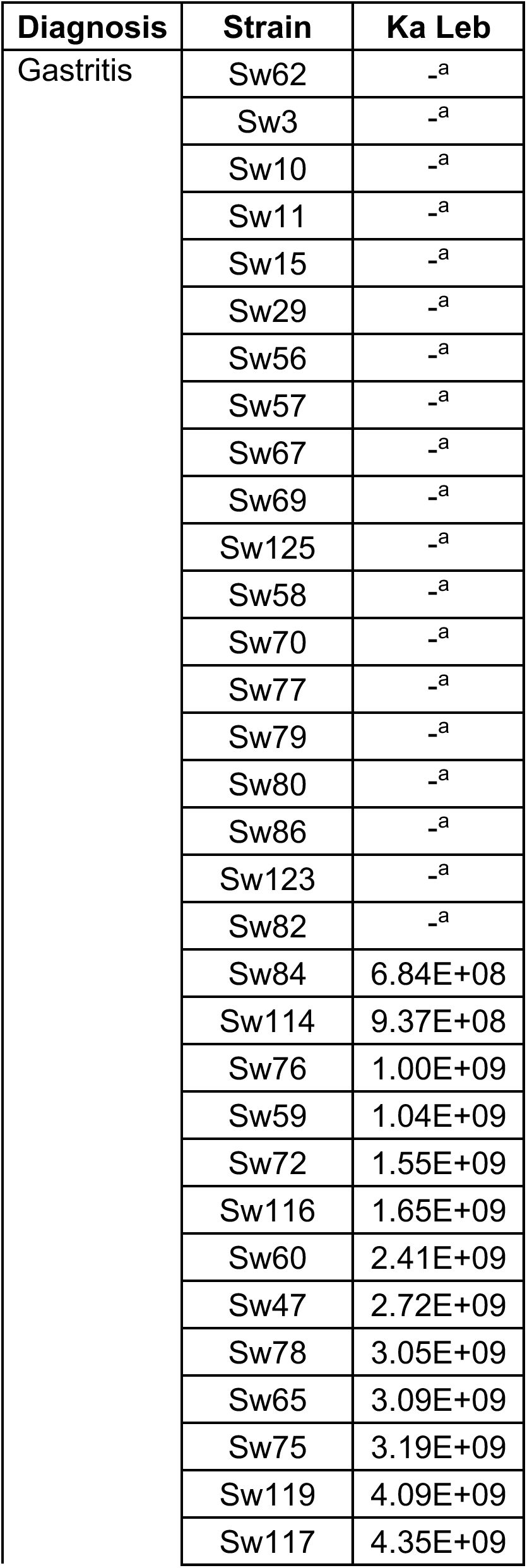
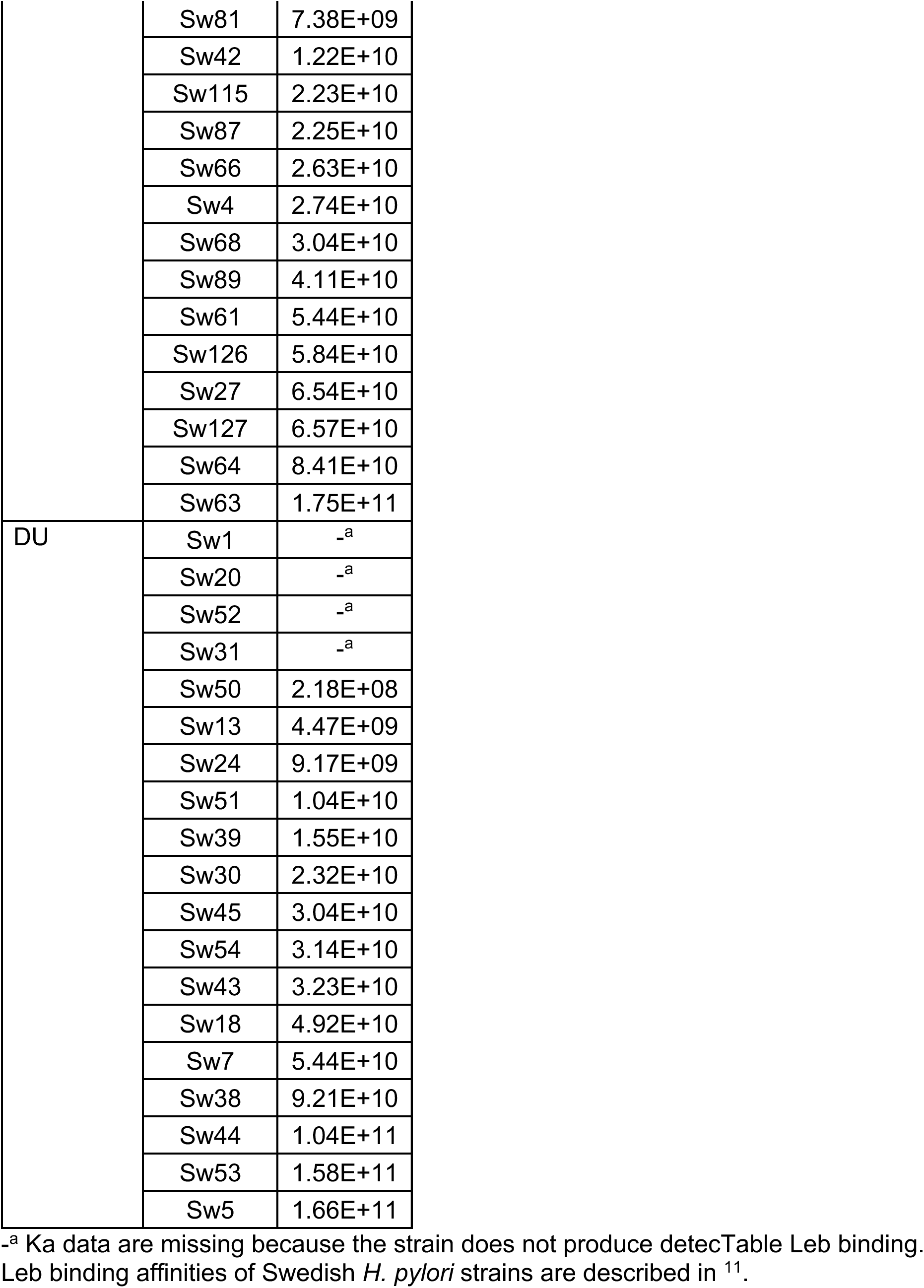
Leb-binding affinities of Swedish strains.

**Table S2B related to Figure S1G (Leb-Cocktail) and Figure S3A (Leb-Hot).**
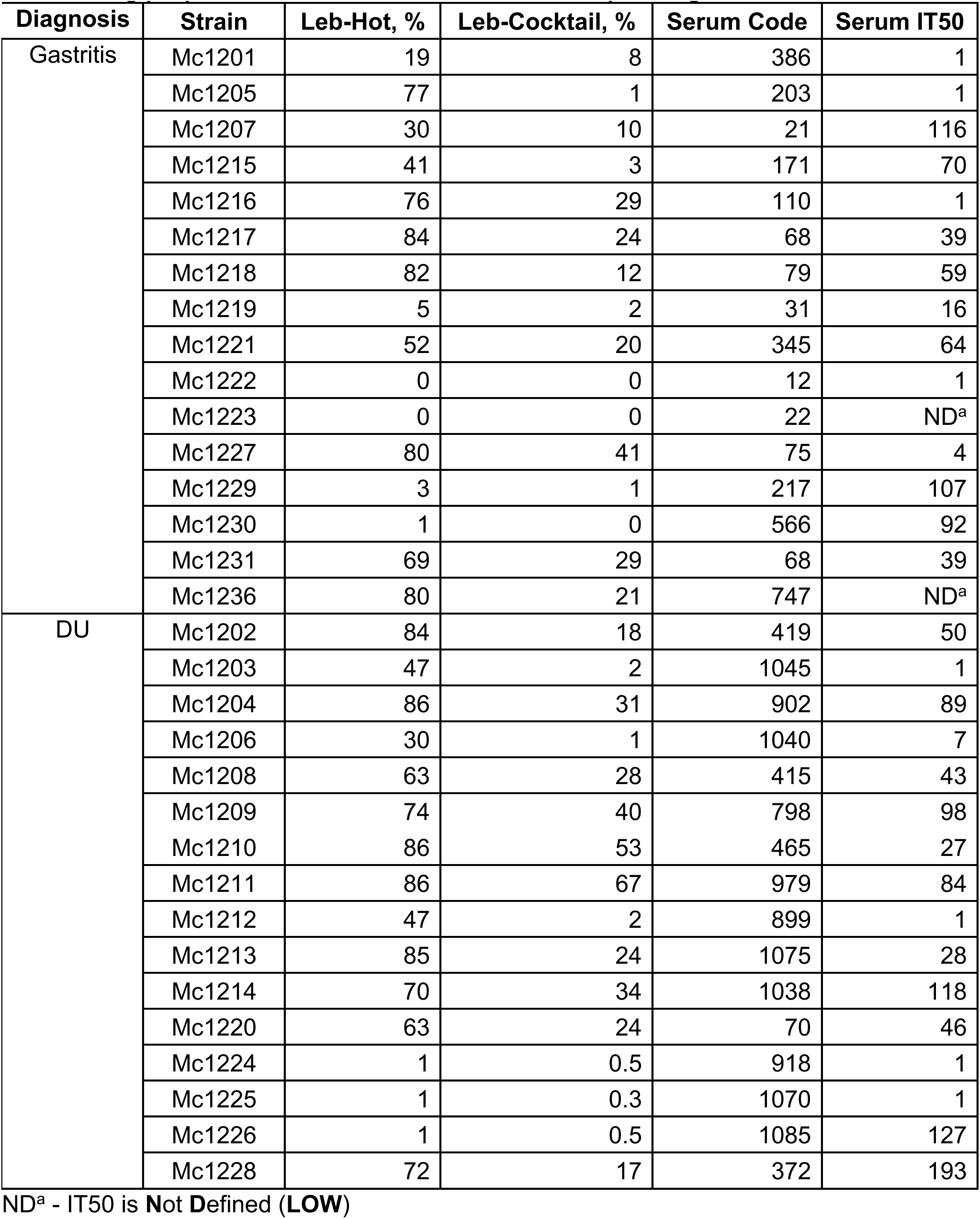
Leb-binding properties of Mexican strains and corresponding serum IT50s.

**Table S3 related to Figure 3 and Figure S3.**
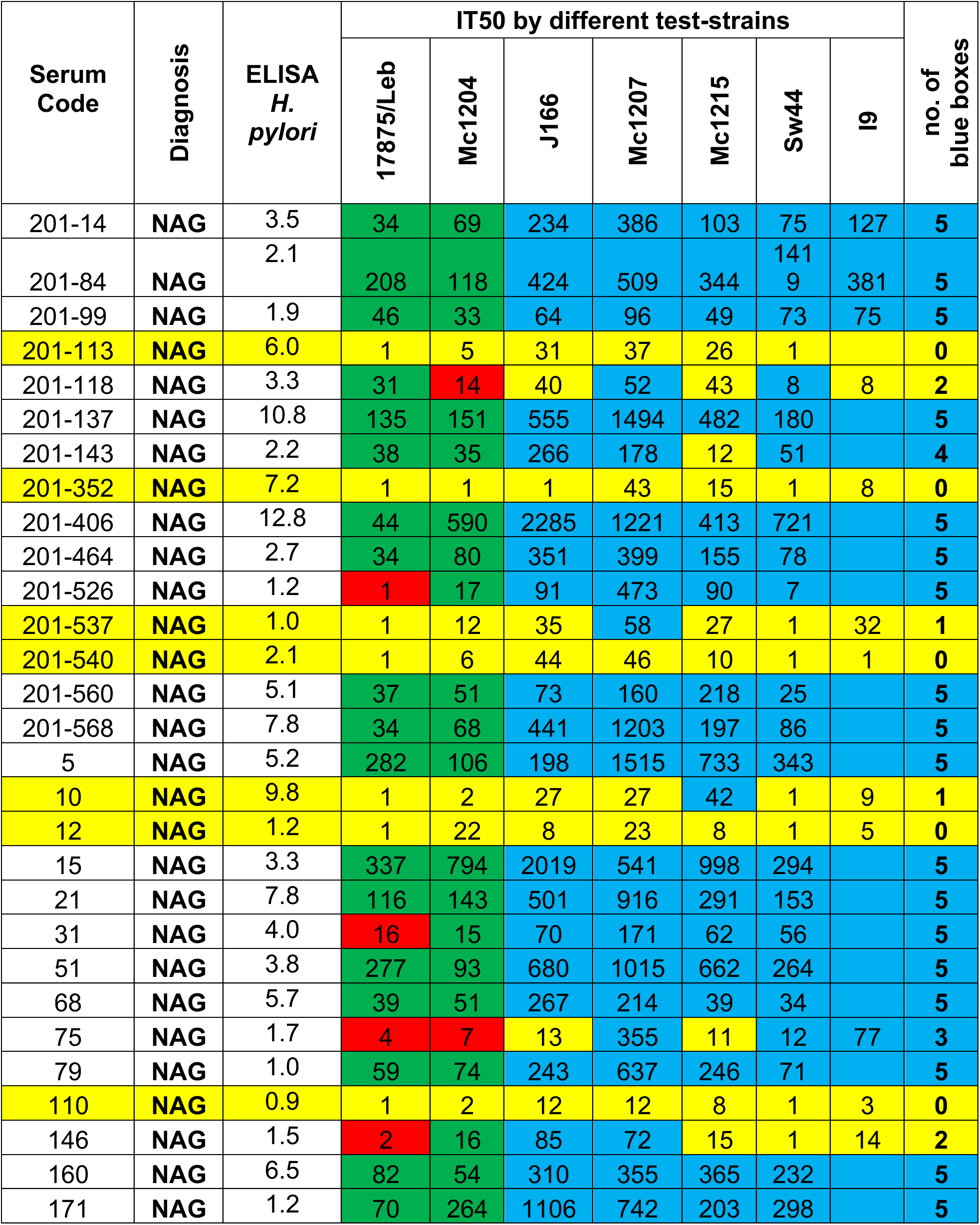
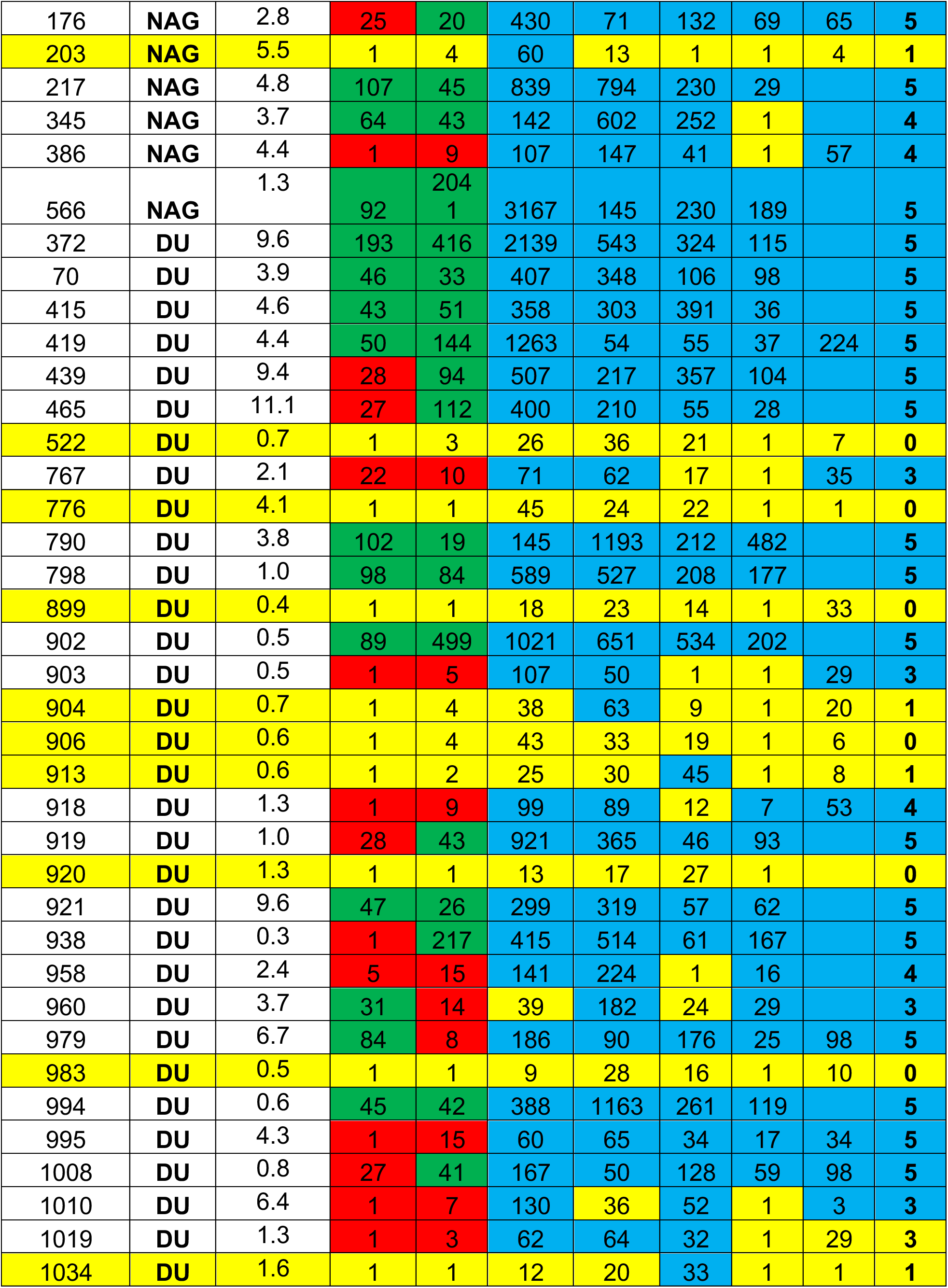
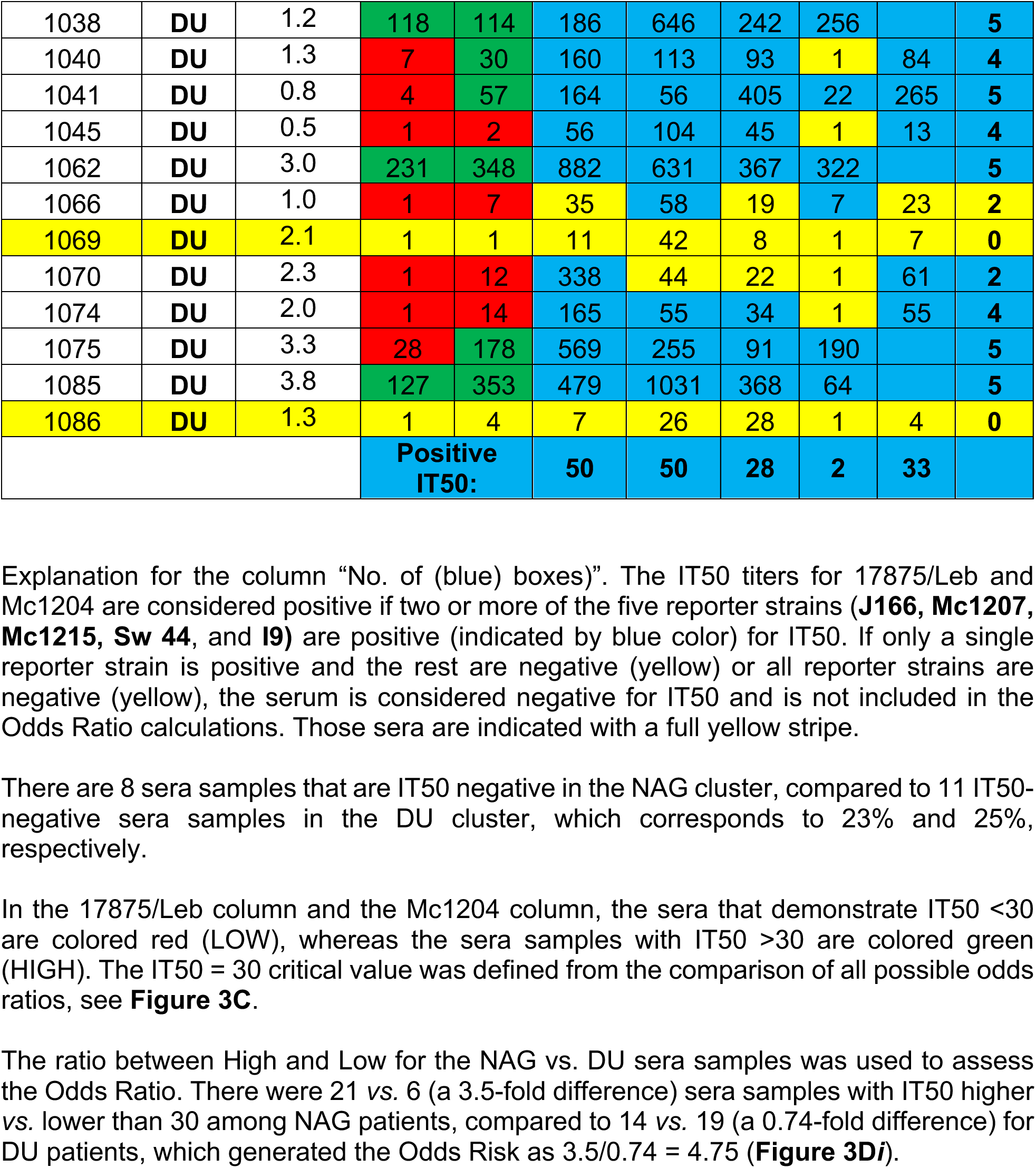
Sera IT50 from Non-Atrophic Gastritis (NAG) and Duodenal Ulcer disease (DU) patients.

**Table S4 related to Figure 1 and Figure S1 and Figure 3E.**
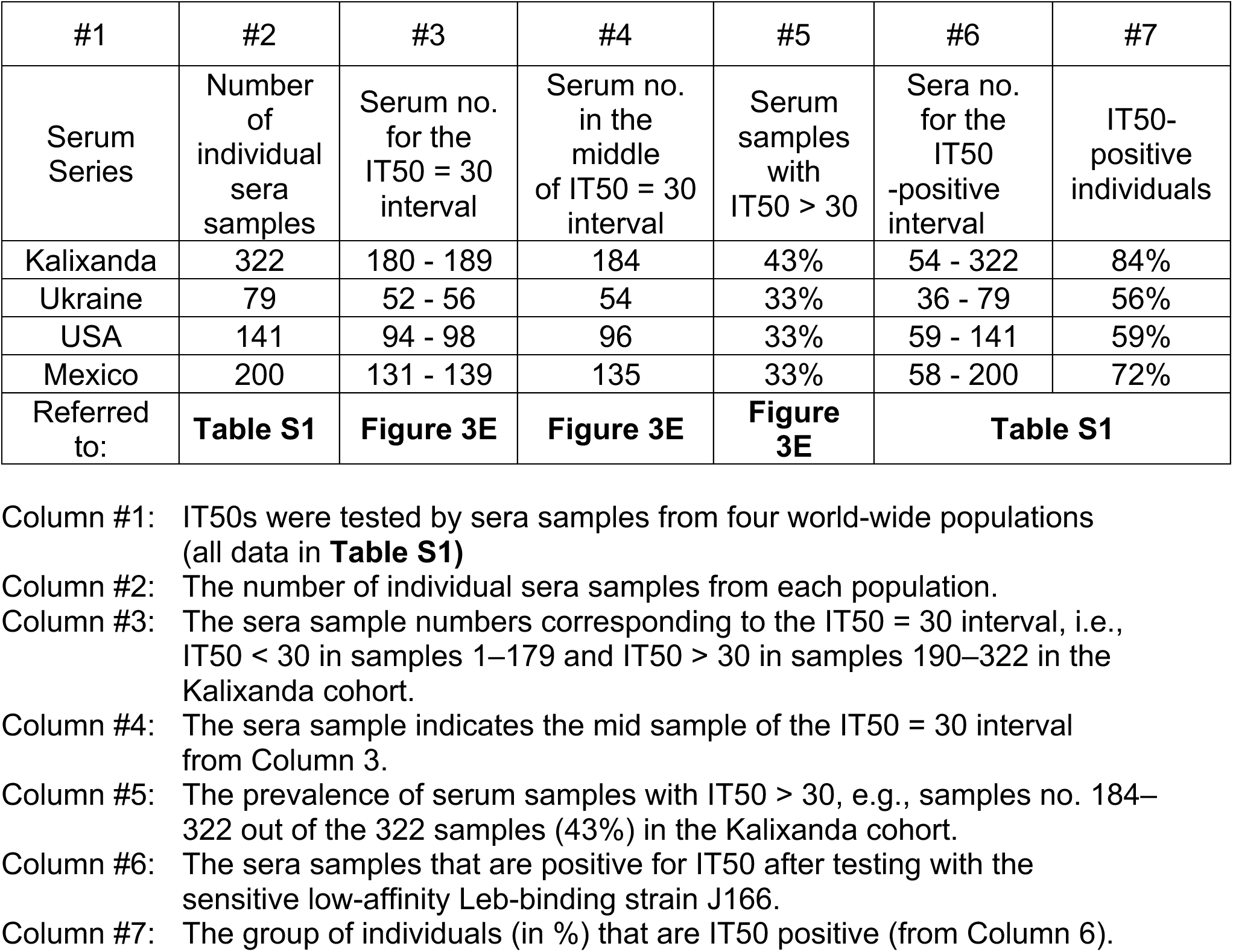
The locations of the IT50 = 30 interval in the different cohorts were identified by strain 17875/Leb, whereas the locations of IT50 positive *vs*. negative sera in the different cohorts were identified by strain J166.

**Table 5 related to Fig 5 and Figure S5.**
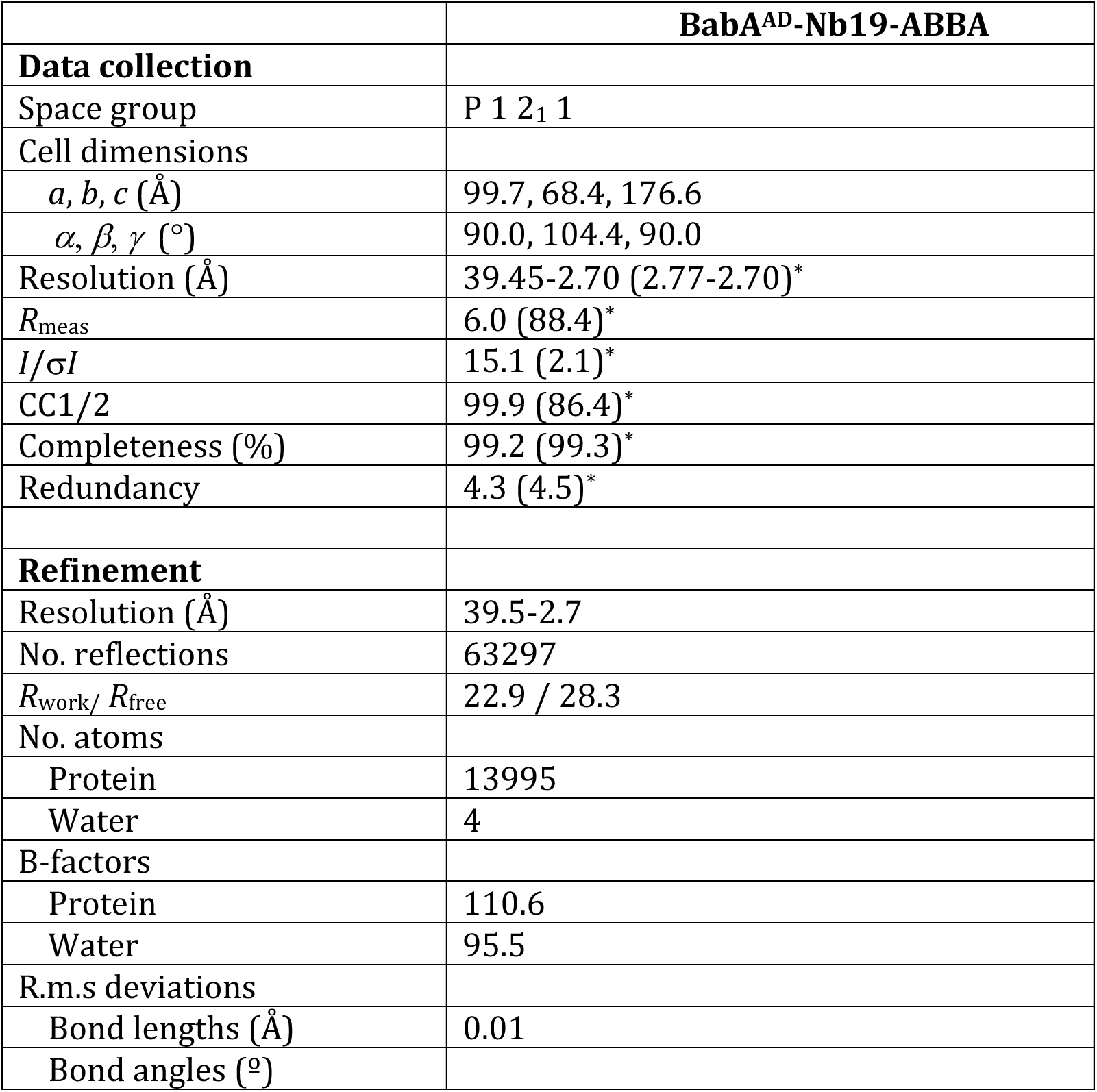
Crystallographic data collection and refinement statistics for the BabA protein co-crystallized with ABbA (Table 5).

**Table S6A related to Fig 6 and fig S6.**
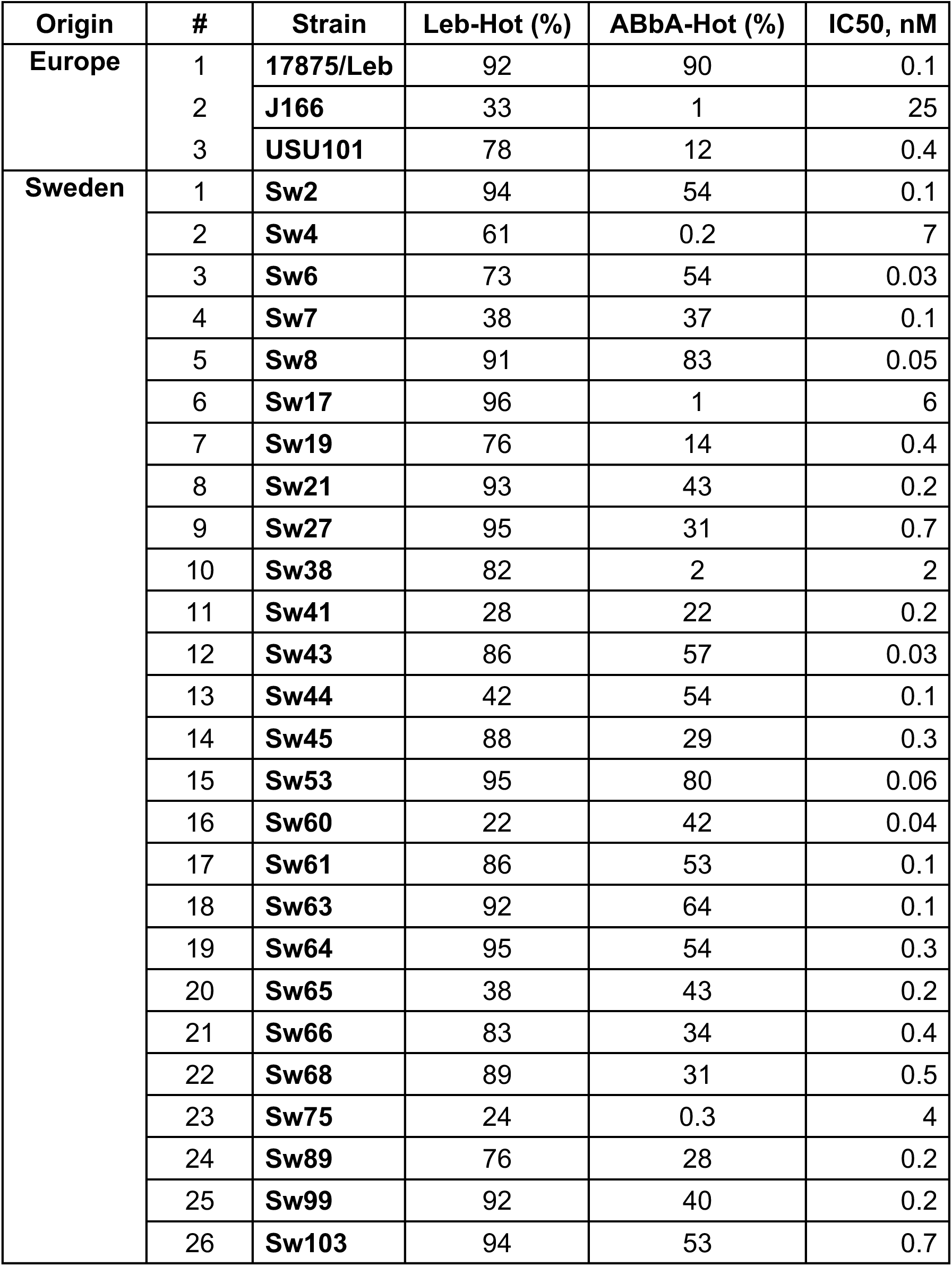
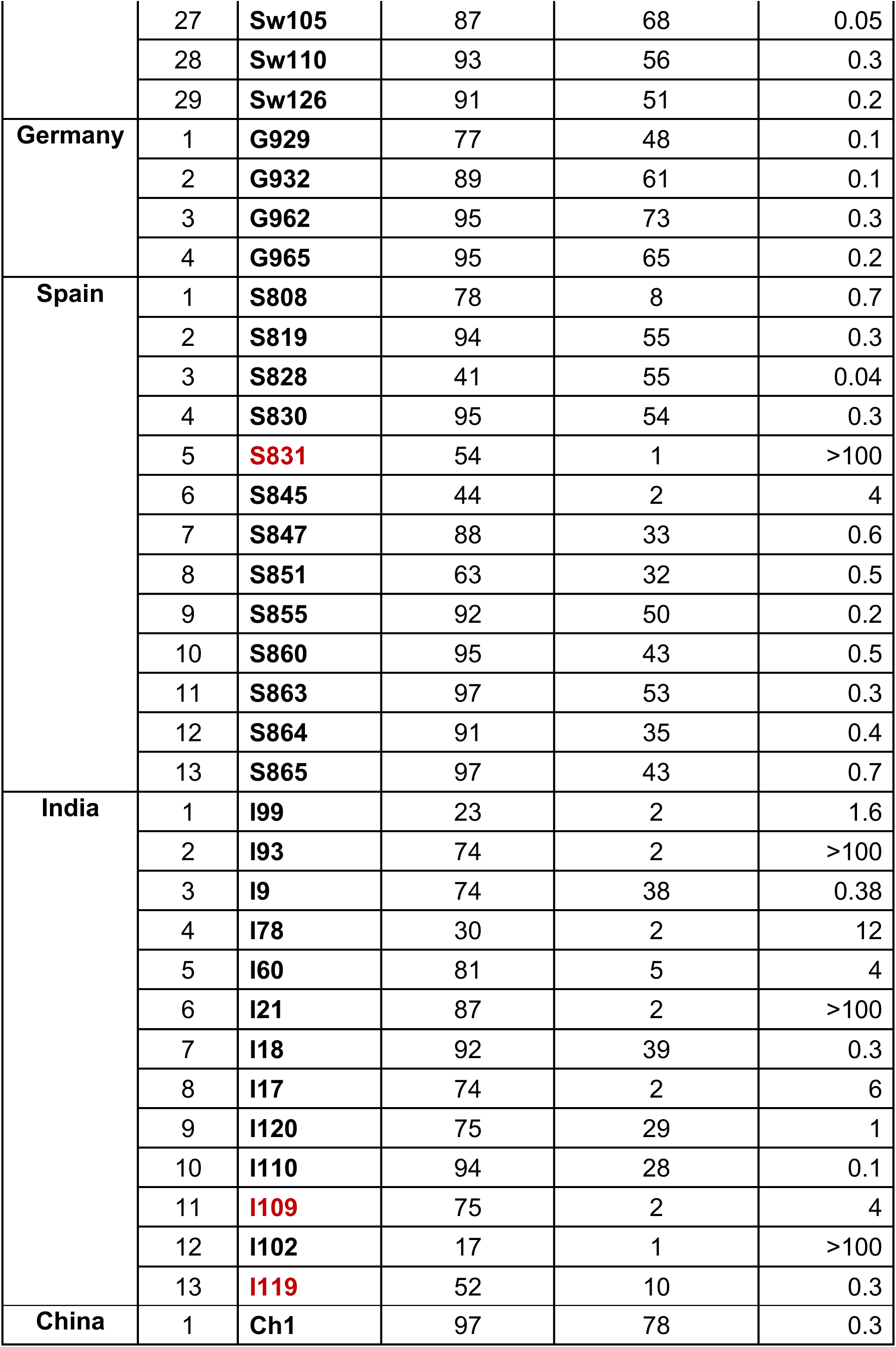
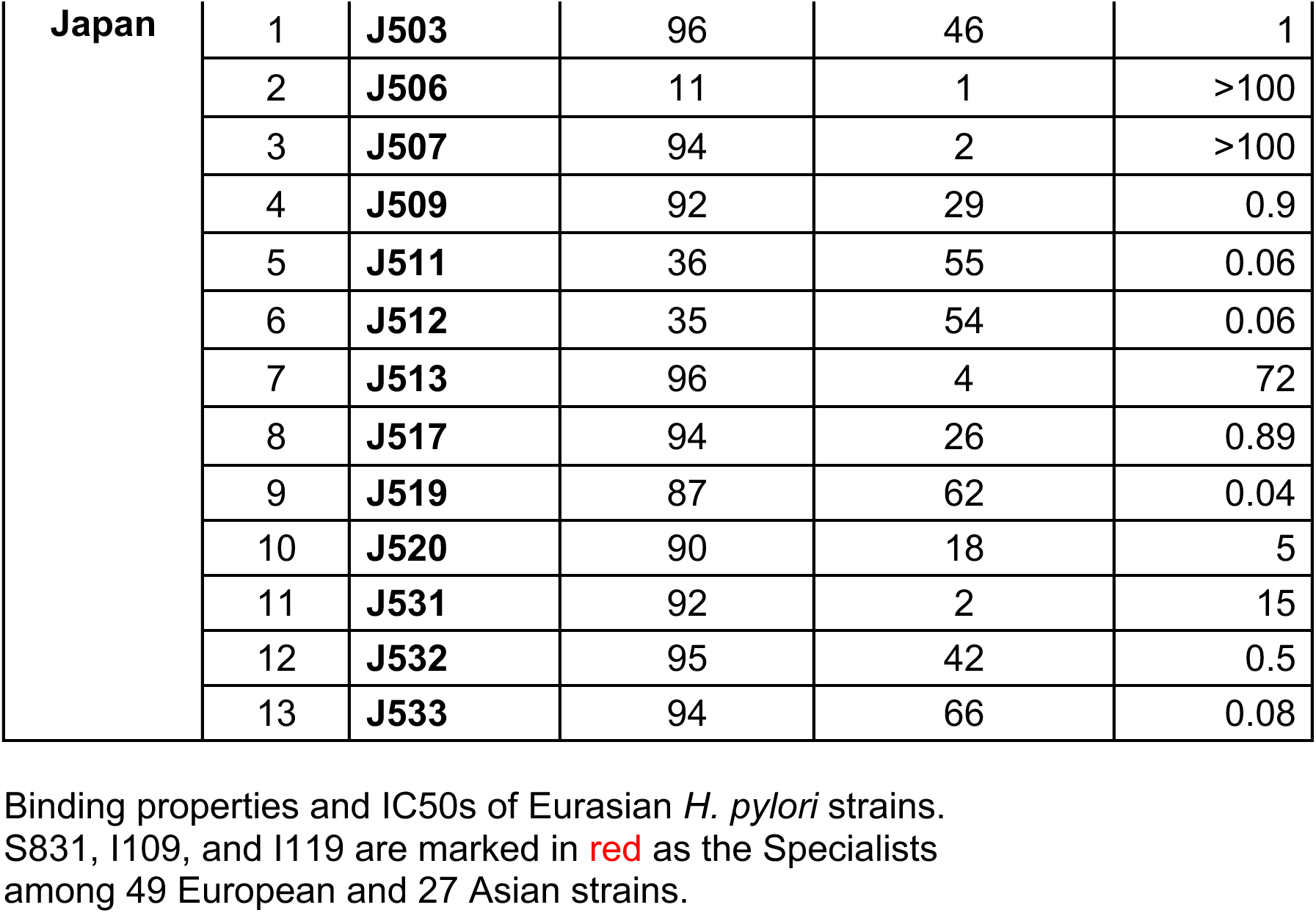
Binding properties and IC50 of Eurasian *H. pylori* strains. S831, I109, and I119 are marked in red as the Specialists among 49 European and 27 Asian strains.

**Table S6B related to Fig 6 and fig S6.**
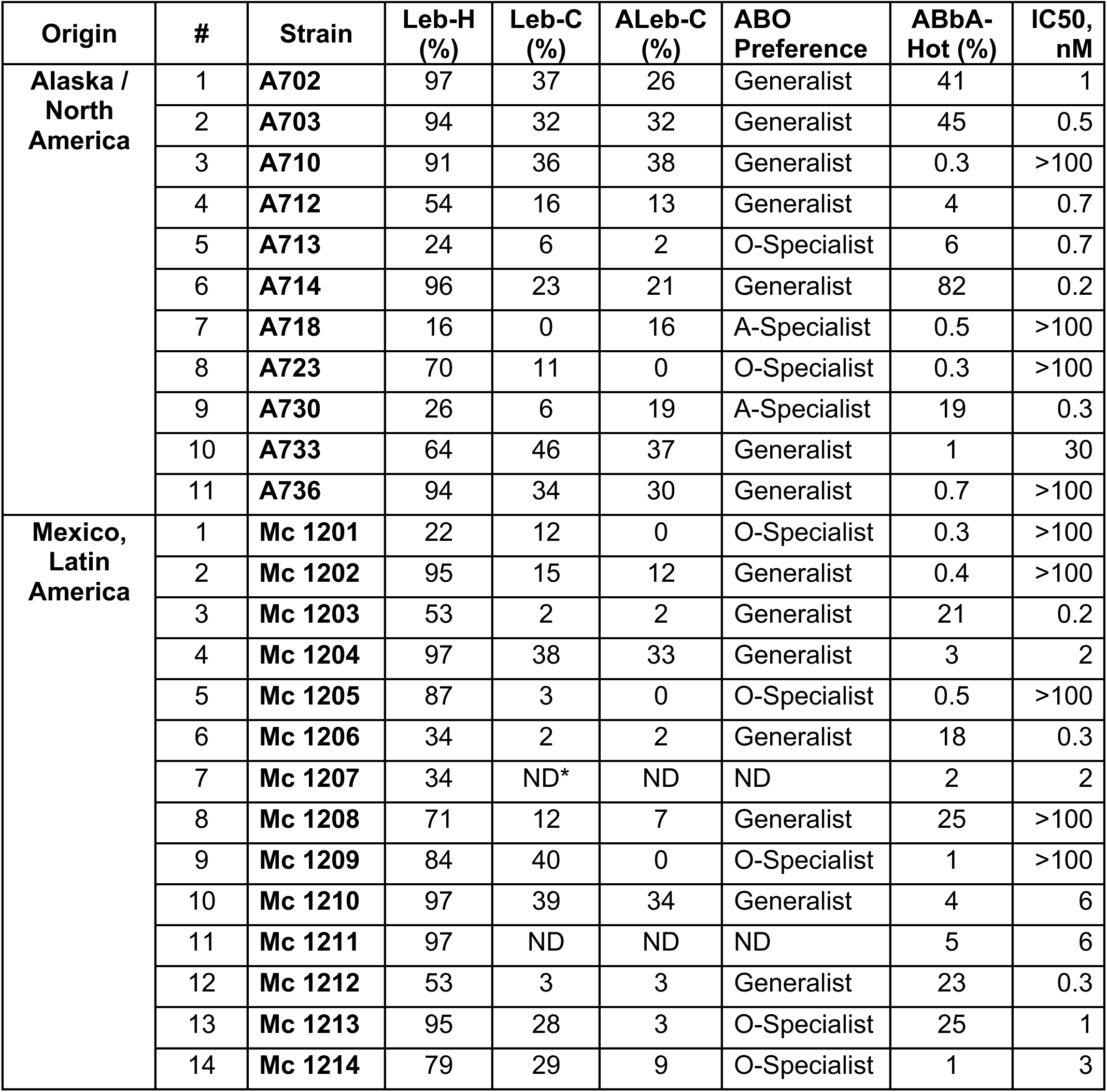
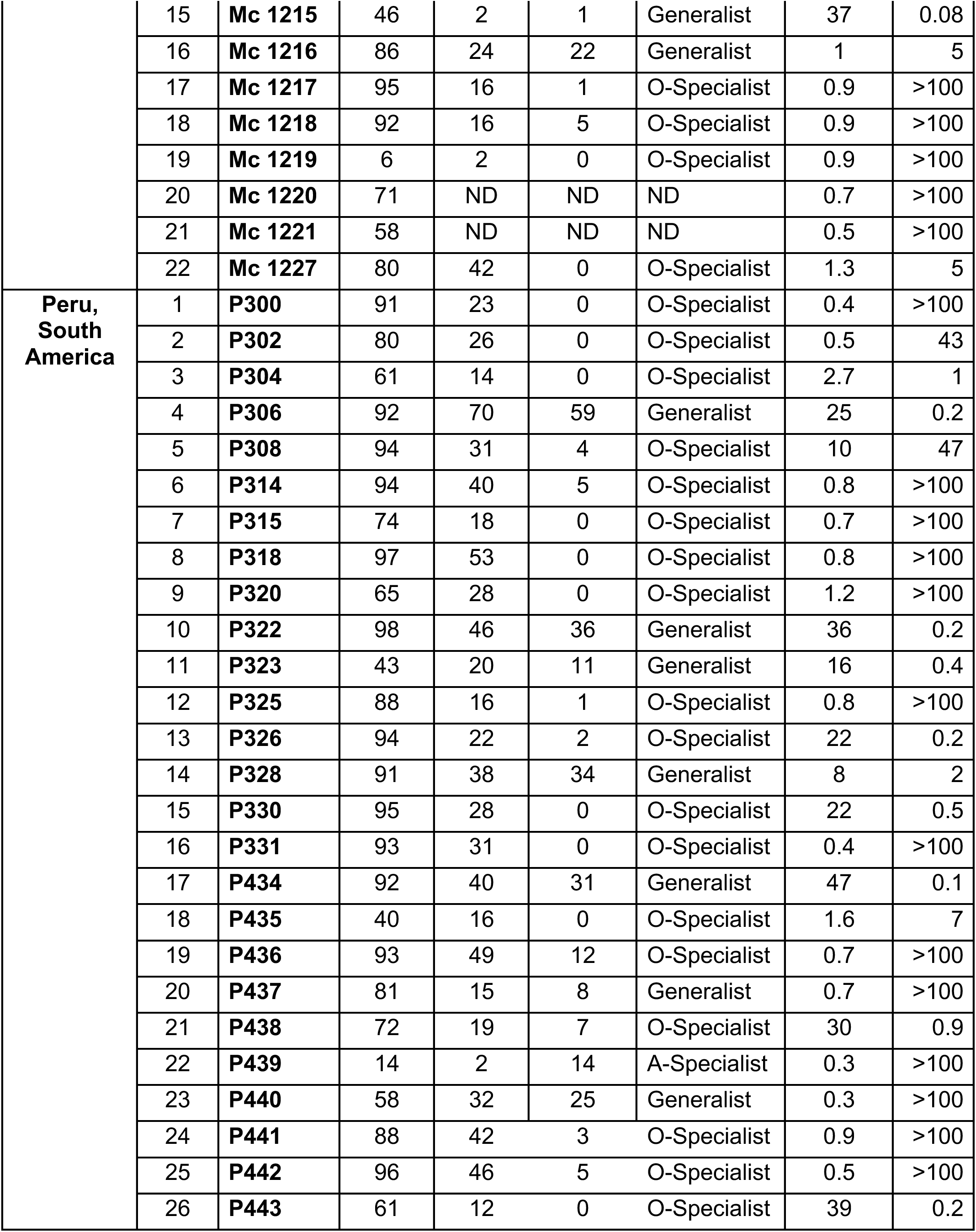
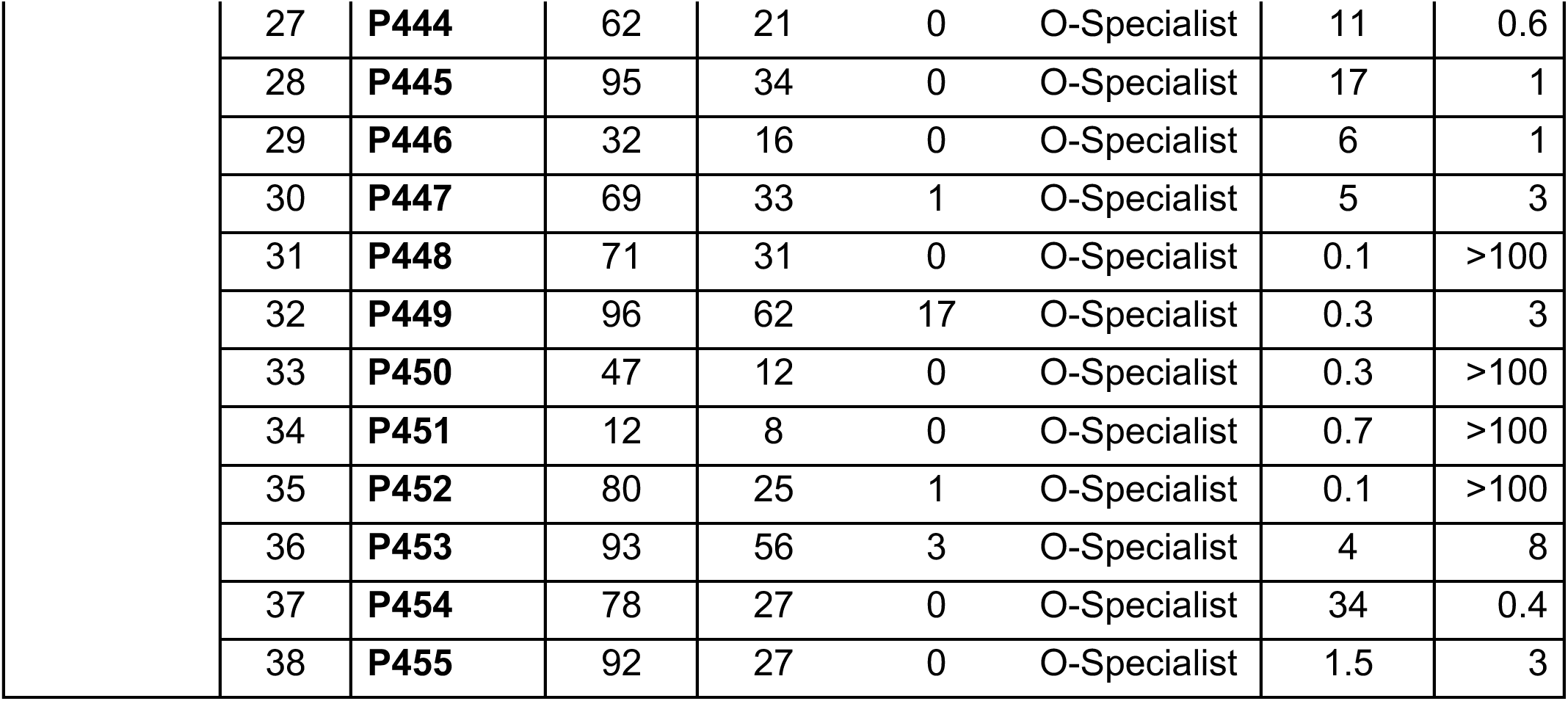
Binding properties and preferences and IC50s of North and South Indigenous American and Latin-American *H. pylori* strains. Leb-H, % - binding to **H**ot HSA-Leb conjugate Leb-C, % - binding to **C**ocktail-HSA-Leb conjugate ALeb-C, % - binding to **C**ocktail-HSA-ALeb conjugate ND - Not Determined O - Specialists are the strains with a >2.5 ratio in Leb/ALeb binding ^11^. A - Specialists are the strains with a <0.4 ratio in Leb/ALeb binding ^11^.

**Table S7 related to Fig 6 and fig S6.**
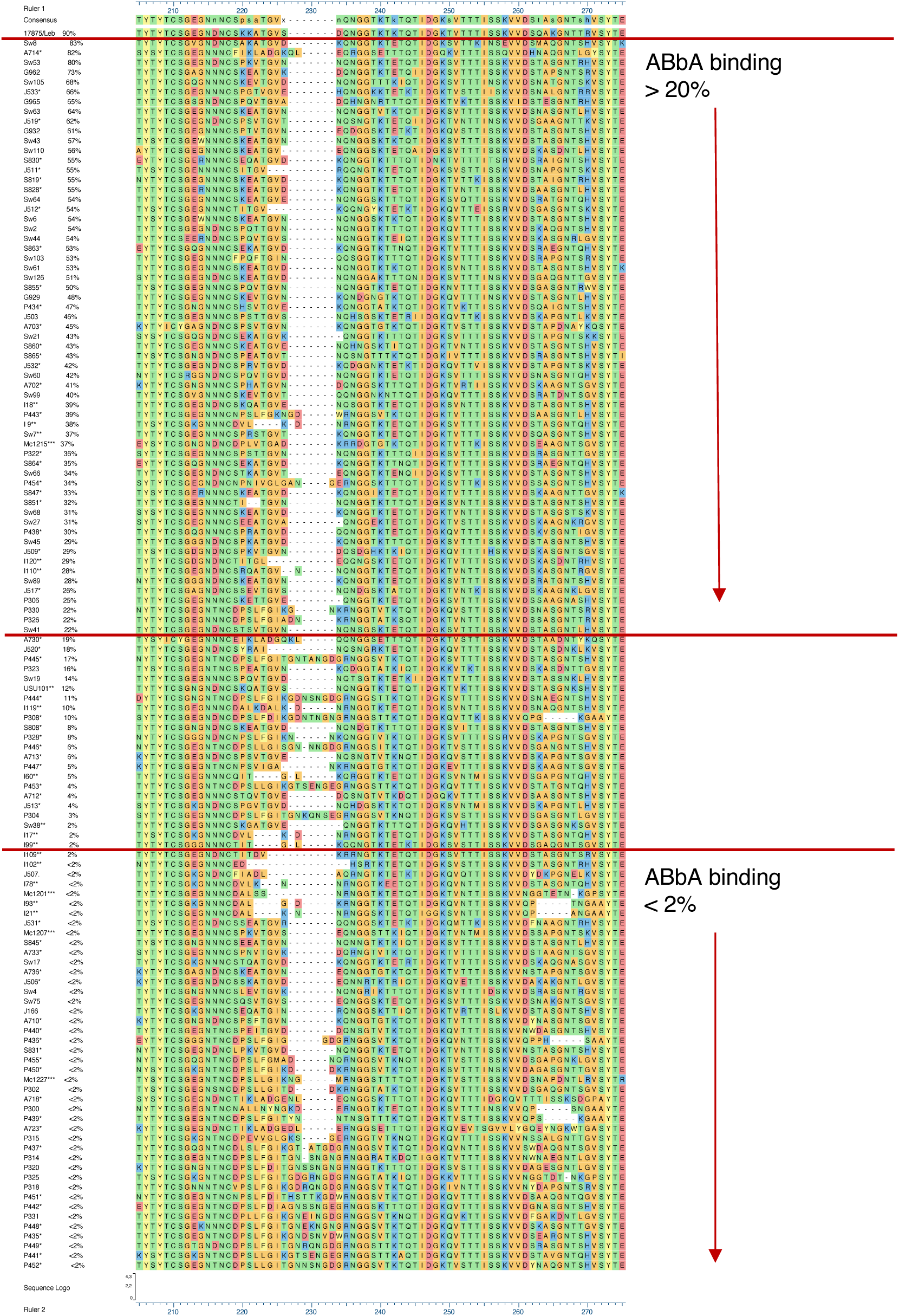
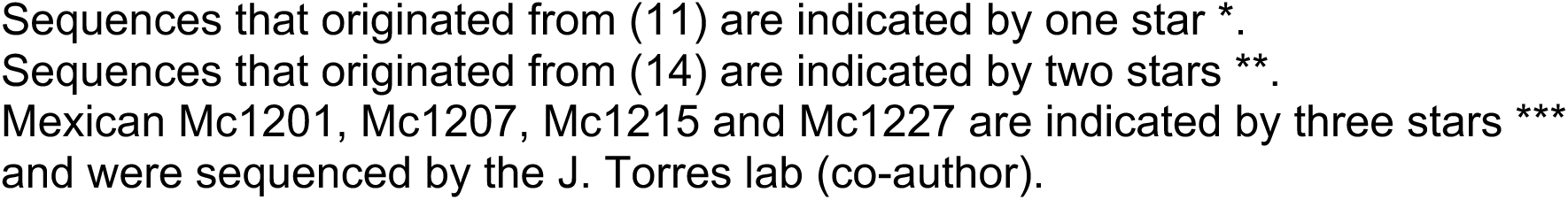
BabA alignment (aa185-247) according to ABbA binding strength.

**Table S8 related to Figure 6 and Figure S6.**
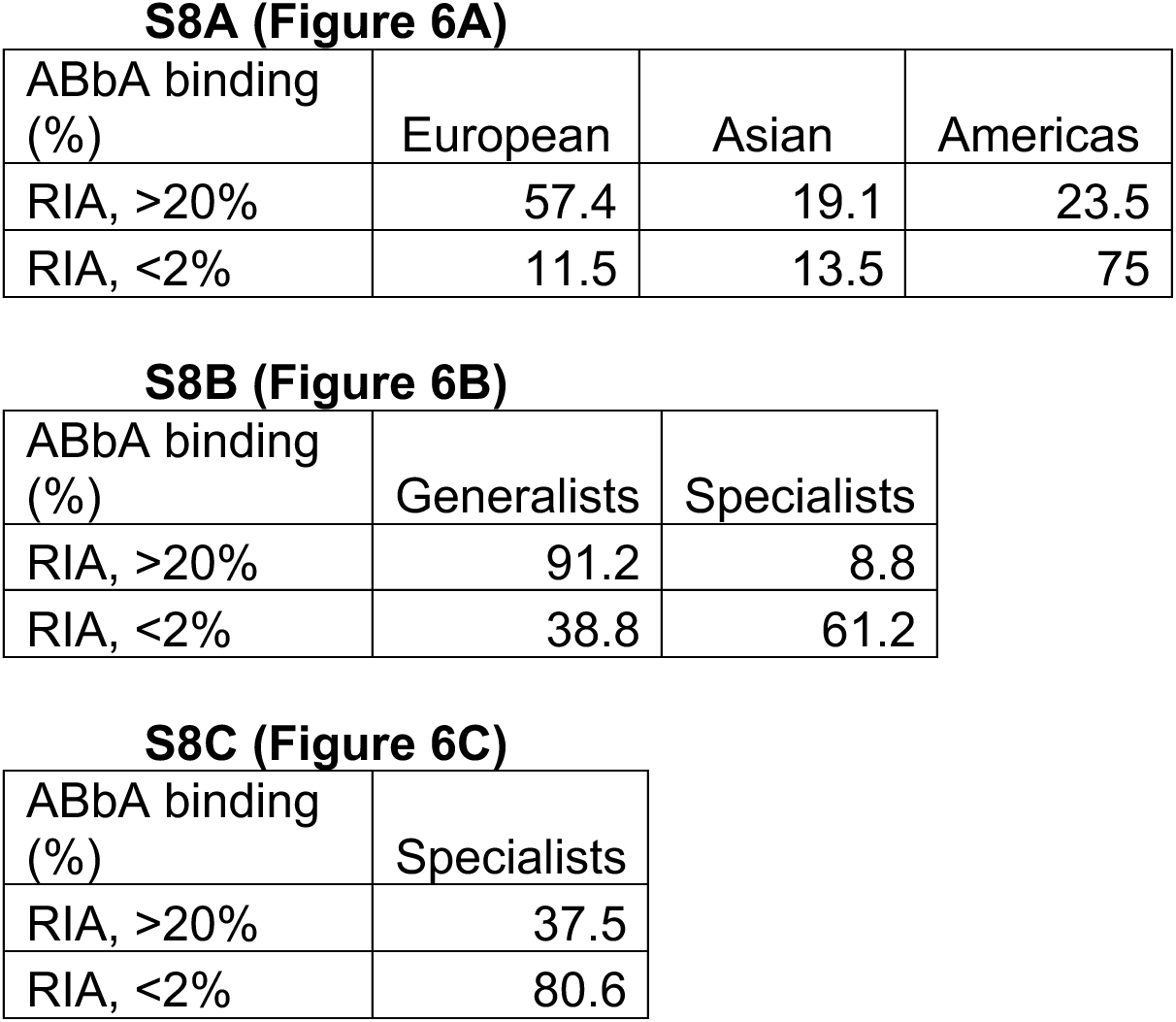
ABbA binding properties of global *H. pylori* strains and of Generalist *vs*. Specialist *H. pylori* strains.

